# The genome of the cereal pest *Sitophilus oryzae*: a transposable element haven

**DOI:** 10.1101/2021.03.03.408021

**Authors:** Nicolas Parisot, Carlos Vargas-Chavez, Clément Goubert, Patrice Baa-Puyoulet, Séverine Balmand, Louis Beranger, Caroline Blanc, Aymeric Bonnamour, Matthieu Boulesteix, Nelly Burlet, Federica Calevro, Patrick Callaerts, Théo Chancy, Hubert Charles, Stefano Colella, André Da Silva Barbosa, Elisa Dell’Aglio, Alex Di Genova, Gérard Febvay, Toni Gabaldon, Mariana Galvão Ferrarini, Alexandra Gerber, Benjamin Gillet, Robert Hubley, Sandrine Hughes, Emmanuelle Jacquin-Joly, Justin Maire, Marina Marcet-Houben, Florent Masson, Camille Meslin, Nicolas Montagne, Andrés Moya, Ana Tereza Ribeiro de Vasconcelos, Gautier Richard, Jeb Rosen, Marie-France Sagot, Arian F.A. Smit, Jessica M. Storer, Carole Vincent-Monegat, Agnès Vallier, Aurélien Vigneron, Anna Zaidman-Remy, Waël Zamoum, Cristina Vieira, Rita Rebollo, Amparo Latorre, Abdelaziz Heddi

## Abstract

Among beetles, the rice weevil *Sitophilus oryzae* is one of the most important pests causing extensive damage to cereal in fields and to stored grains. *S. oryzae* has an intracellular symbiotic relationship (endosymbiosis) with the Gram-negative bacterium *Sodalis pierantonius* and is a valuable model to decipher host-symbiont molecular interactions.

We sequenced the *Sitophilus oryzae* genome using a combination of short and long reads to produce the best assembly for a Curculionidae species to date. We show that *S. oryzae* has undergone successive bursts of transposable element (TE) amplification, representing 72% of the genome. In addition, we show that many TE families are transcriptionally active, and changes in their expression are associated with insect endosymbiotic state. *S. oryzae* has undergone a high gene expansion rate, when compared to other beetles. Reconstruction of host-symbiont metabolic networks revealed that, despite its recent association with cereal weevils (30 Kyear), *S. pierantonius* relies on the host for several amino acids and nucleotides to survive and to produce vitamins and essential amino-acids required for insect development and cuticle biosynthesis.

In addition to being an agricultural pest and a valuable endosymbiotic system, *S. oryzae* can be a remarkable model for studying TE evolution and regulation, along with the impact of TEs on eukaryotic genomes.

## Introduction

Beetles account for approximately 25% of known animals, with an estimated number of 400 000 described species (Hunt et al. 2007; Stork et al. 2015; Hammond 1992). Among them, Curculionidae (true weevils) is the largest animal family described, comprising about 70 000 species (Hunt et al. 2007; McKenna et al. 2009; Oberprieler et al. 2007). Despite being often associated with ecological invasion and ecosystem degradation, only three Curculionidae genomes are publicly available to date (Vega et al. 2015; Keeling et al. 2013; Hazzouri et al. 2020). Among the cereal weevils, the rice weevil *Sitophilus oryzae* is one of the most important pests of crops of high agronomic and economic importance (wheat, maize, rice, sorghum and barley), causing extensive quantitative and qualitative losses in field, stored grains and grain products throughout the world (Zunjare et al. 2016; Longstaff 1981; Grenier et al. 1997). Moreover, this insect pest is of increasing concern due to its ability to rapidly evolve resistance to insecticides such as phosphine, a fumigant used to protect stored grains from insect pests (Champ and Dyte 1977; Nguyen et al. 2015; Mills 2000).

Like other holometabolous insects, the life cycle of *S. oryzae* can be divided into four stages: egg, larva, pupa and adult (Figure 1). Females drill a small hole in the grain, deposit a single egg and seal it with secretions from their ovipositor. Up to six eggs can be laid daily by each female, totaling around 400 eggs over its entire lifespan (Campbell 2005). Larvae develop and pupate within the grain kernel, metamorphose, and exit the grain as adults. The whole process takes on average 30 days (Longstaff 1981). Like many insects living on nutritionally poor diets, cereal weevils permanently associate with nutritional intracellular bacteria (endosymbionts) that supply them with nutrients that are not readily available in the grains, thereby increasing their fitness and invasive power. The endosymbiont of *S. oryzae*, the gamma-proteobacterium *Sodalis pierantonius* (Oakeson et al. 2014; Heddi et al. 1998), is housed within specialized host cells, named bacteriocytes, that group together into an organ, the bacteriome (Heddi et al. 2001). Contrasting with most studied symbiotic insects, the association between *Sitophilus* spp. and *S. pierantonius* was established recently (less than 30 000 years ago), probably following the replacement of the ancestor endosymbiont, Candidatus *Nardonella*, in the Dryophthorinae subfamily (Lefèvre et al. 2004; Clayton et al. 2012). As a result, contrary to long-lasting endosymbiotic associations, the genome of *S. pierantonius* is GC rich (56.06%), and its size is similar to that of free-living bacteria (4.5 Mbp) (Oakeson et al. 2014). Moreover, it encodes genes involved in bacterial infection, including Type Three Secretion Systems (TTSS), as well as genes encoding Microbial Associated Molecular Patterns (MAMPs) that trigger Pattern Recognition Receptors (PRR), and are usually absent or reduced in bacteria involved in long-lasting associations (Oakeson et al. 2014; Akman et al. 2002; Shigenobu et al. 2000). Nevertheless, many features indicate that the genome of *S. pierantonius* is in a process of degradation, as it contains many pseudogenes (43% of the predicted protein-coding sequences) and a large number of mobile elements (18% of the genome size) (Oakeson et al. 2014; Gil et al. 2008). Finally, it is important to note that no other symbionts, with the exception of the familiar *Wolbachia* endosymbiont in some strains, have been described in *S. oryzae*.

**Figure 1.**
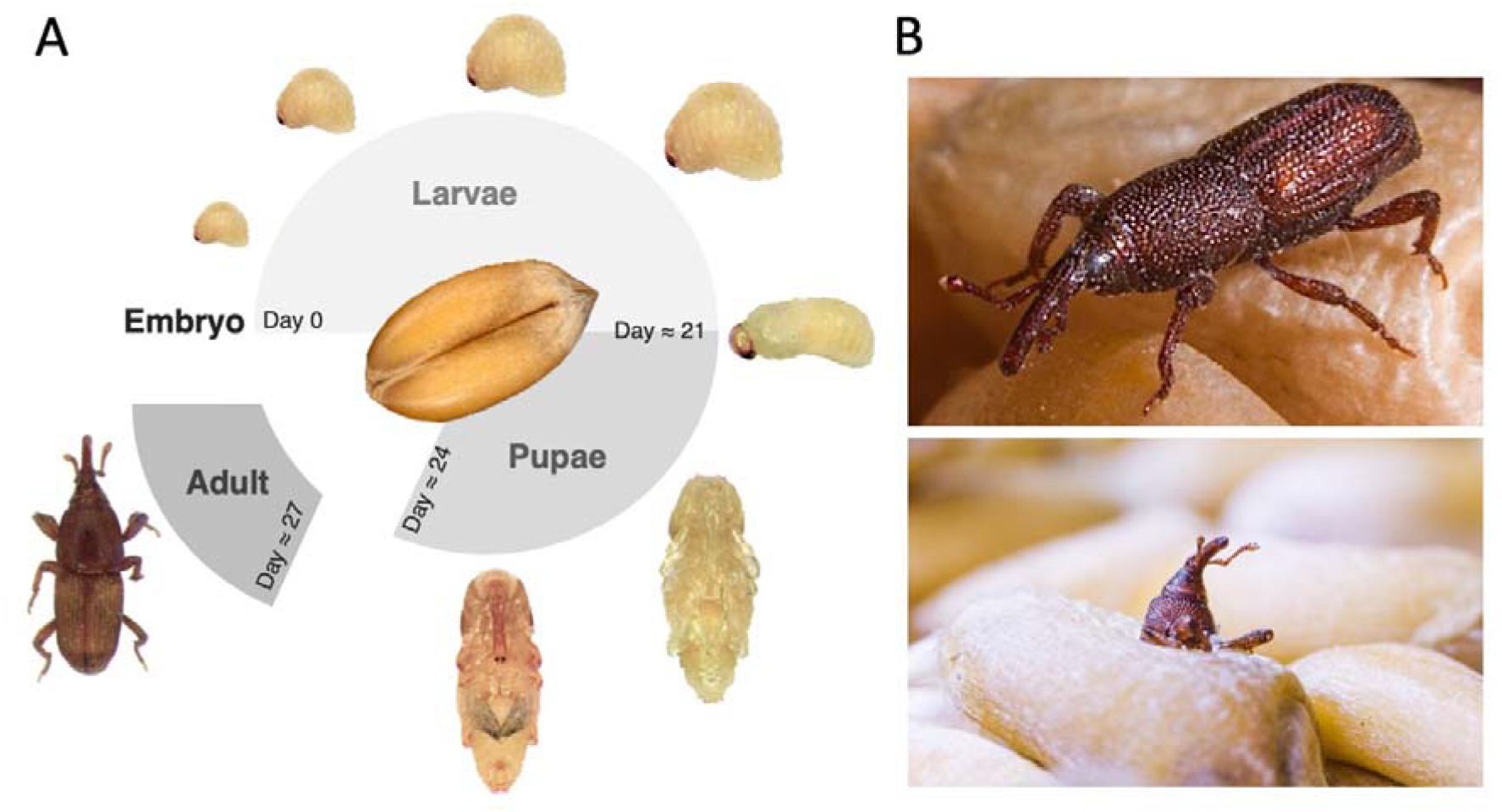
*Sitophilus oryzae* overview. A. Life cycle of cereal weevil *Sitophilus oryzae*. The embryo develops into a larva and pupa, and metamorphoses into a young adult, exiting the grain around 3 days after metamorphosis completion. The developmental times indicated are from a rearing condition at 27 °C and 70% relative humidity. B. Photos of adult *S. oryzae*. Lower panel shows an adult exiting the grain.

In order to help unravel potential adaptive functions and features that could become the basis for identifying novel control strategies for weevils and other major insect pests, we have undertaken the sequencing, assembly and annotation of the genome of *S. oryzae*. Strikingly, the repeated fraction of *S. oryzae*’s genome (repeatome), composed mostly of transposable elements (TEs), is among the largest found to date in insects. TEs, the most versatile DNA units described to date, are sequences present in multiple copies and capable of relocating or replicating within a genome. While most observed TE insertions evolve neutrally or are slightly deleterious, there are a number of documented cases where TEs may facilitate host adaptation (for reviews see (Rebollo et al. 2012; Bourque et al. 2018; Chuong et al. 2017)). For instance, gene families involved in xenobiotic detoxification are enriched in TEs in *Drosophila melanogaster* (Chen and Li 2007), *Plutella xylostella* (You et al. 2013), a major crop pest, and *Myzus persicae*, another phytophagous insect causing significant agronomic losses (Singh et al. 2020). TEs have also been frequently associated with insecticide-resistance in *Drosophila* species (Carareto et al. 2014; Rostant et al. 2012; Mateo et al. 2014). In addition, population genetics studies suggested that more than 84 TE copies in *D. melanogaster* may play a positive role in fitness-related traits (Rech et al. 2019), including xenobiotic resistance (Mateo et al. 2014) and immune response to Gram-negative bacteria (Ullastres et al. 2019).

In eukaryotes, TE content varies drastically and contributes significantly to the size and organization of the genome. From TE-rich genomes as maize (85% (Schnable et al. 2009)), humans (≈45% (Lander et al. 2001)), and the recently sequenced lungfish (≈90% (Meyer et al. 2021)) for instance, to TE-poor genomes, as *D. melanogaster* (12-15% (Adams et al. 2000)), or *Arabidopsis thaliana* (≈10% (The Arabidopsis Genome Initiative 2000)), repeatomes thrive on a high level of diversity. These drastic variations are also observed within animal clades, such as insects, where the proportion of TE ranges from 2% in the Antarctic midge (*Belgica antarctica*) to 65% in the migratory locust (*Locusta migratoria*) (Petersen et al. 2019; Wang et al. 2014; Kelley et al. 2014) and up to 75% in morabine grasshoppers (*Vandiemenella viatica* species) (Palacios-Gimenez et al. 2020). In addition to the overall TE content, the number of different TE families (homogeneous groups of phylogenetically related TE sequences), their size (number of copies per family) and sequence diversity are also very high among insect species (Gilbert et al. 2021). For instance, SINEs (Short INterspersed Elements) are almost absent from most insect genomes, but many lepidopterans harbor these elements (Gilbert et al. 2021). In flies, Long Terminal Repeats retrotransposons (LTRs) are a staple of the *Drosophila* genus, but such TEs are nearly absent from other dipteran genomes (*e.g*. *Glossina brevipalpis* and *Megaselia scalaris*) (Gilbert et al. 2021). Recent advances in sequencing have dramatically increased the level to which TEs can be studied across species and reveal that such variations can persist even within recently diverged groups, as observed within *Drosophila* species (Sessegolo et al. 2016) or among *Heliconius* butterflies (Ray et al. 2019). An increasing number of insect genomes are reported with large repeatomes (*e.g*. *Aedes aegypti* and *Ae. albopictus* 40-50% (Goubert et al. 2015; Nene et al. 2007), *L. migratoria* 60-65% (Wang et al. 2014; Petersen et al. 2019), *Dendrolimus punctatus* 56% (Zhang et al. 2020), *Vandiemenella viatica* species 66-75% (Palacios-Gimenez et al. 2020)).

Here we present the genome of *S. oryzae*, with a strong focus on the repeatome, its largest genomic compartment, spanning over ≈74% of the assembly. *S. oryzae* represents a model system for stored grain pests, host-TE evolutionary biology, and the study of the molecular mechanisms acting at the early steps of symbiogenesis. Moreover, the features uncovered suggest that *S. oryzae* and its relatives have the potential to become a platform to study the interplay between TEs, host genomes and endosymbionts.

## Results and discussion

### Genome assembly and annotation

We have sequenced and assembled the genome of the rice weevil *S. oryzae* at a base coverage depth of 142X using a combination of short and long read strategies (see Methods). The karyotype of *S. oryzae* comprises 22 chromosomes (Silva et al. 2018), and the genome assembly consists of 2 025 scaffolds spanning 770 Mbp with a N50 of 2.86 Mbp, demonstrating a high contiguity compared to other Coleopteran genomes (Table 1). The assembly size is consistent with the genome size measured through flow cytometry (769 Mbp in females and 768 Mbp in males (Silva et al. 2018)). We assessed the completeness of the genome assembly using BUSCO (Seppey et al. 2019) (97.9% complete and 0.7% fragmented), and along with the aforementioned statistics, *S. oryzae* is the best assembled Curculionidae genome to date (Keeling et al. 2013; McKenna et al. 2016; Tribolium Genome Sequencing Consortium et al. 2008) (Table 1). The complete analysis of gene content and function can be found in Supplemental file S1.

**Table 1.**
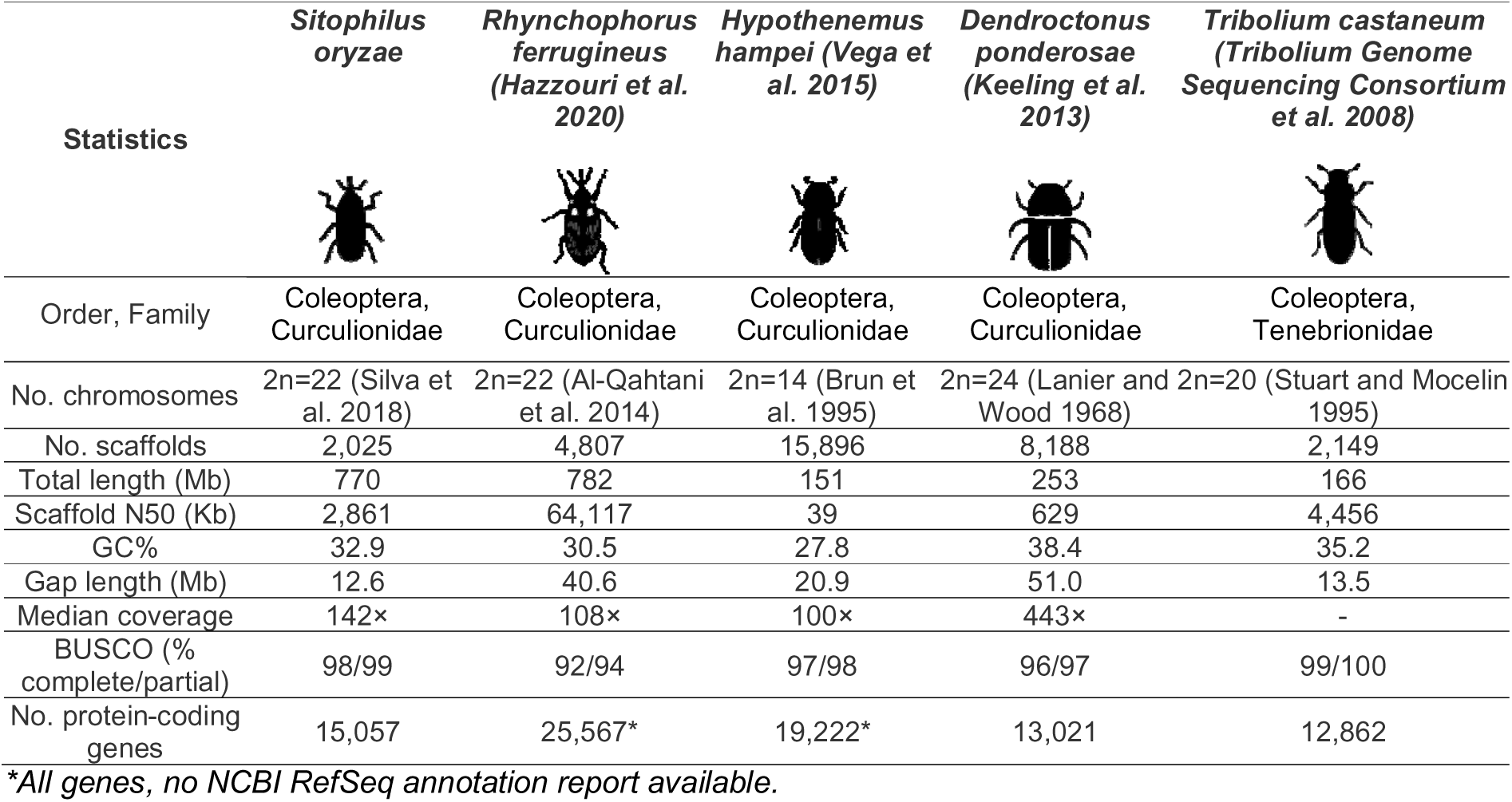
Assembly statistics of *S. oryzae*’s genome in comparison to Curculionidae genomes and *T. castaneum*

| Statistics | *Sitophilus oryzae* | *Rhynchophorus ferrugineus* (Hazzouri et al. 2020) | *Hypothenemus hampei* (Vega et al. 2015) | *Dendroctonus ponderosae* (Keeling et al. 2013) | *Tribolium castaneum* (Tribolium Genome Sequencing Consortium et al. 2008) |
| --- | --- | --- | --- | --- | --- |
| Order, Family | Coleoptera, Curculionidae | Coleoptera, Curculionidae | Coleoptera, Curculionidae | Coleoptera, Curculionidae | Coleoptera, Tenebrionidae |
| No. chromosomes | 2n=22 (Silva et al. 2018) | 2n=22 (Al-Qahtani et al. 2014) | 2n=14 (Brun et al. 1995) | 2n=24 (Lanier and Wood 1968) | 2n=20 (Stuart and Mocelin 1995) |
| No. scaffolds | 2,025 | 4,807 | 15,896 | 8,188 | 2,149 |
| Total length (Mb) | 770 | 782 | 151 | 253 | 166 |
| Scaffold N50 (Kb) | 2,861 | 64,117 | 39 | 629 | 4,456 |
| GC% | 32.9 | 30.5 | 27.8 | 38.4 | 35.2 |
| Gap length (Mb) | 12.6 | 40.6 | 20.9 | 51.0 | 13.5 |
| Median coverage | 142x | 108x | 100x | 443x | - |
| BUSCO (% complete/partial) | 98/99 | 92/94 | 97/98 | 96/97 | 99/100 |
| No. protein-coding genes | 15,057 | 25,567* | 19,222* | 13,021 | 12,862 |

\*All genes, no NCBI RefSeq annotation report available.

### Annotation of the *Sitophilus oryzae* genome

Among the different pathways we were able to decipher in the genome of *S. oryzae*, we present here highlights of the main annotation efforts, followed by a detailed analysis of the TE content and impact on the host genome. A comprehensive analysis for each highlight is presented as Supplemental Notes in Supplemental file S1.

#### Phylome and horizontal gene transfer

*Sitophilus oryzae* has a high gene expansion rate when compared to other beetles. Some of the families with the largest expansions include genes coding for proteins with DNA binding motifs, potentially regulating functions specific to this clade. Olfactory receptors, antimicrobial peptides (AMPs) and P450 cytochromes were expanded as well, probably in response to their ecological niche and lifestyle. Additionally, we noticed an expansion of plant cell wall degrading enzymes that originated from horizontal gene transfer (HGT) events from both bacteria and fungi. Given the intimate relationship between *S. oryzae* and its endosymbiont, including the permanent infection of the female germline, we searched for evidence for HGT in the weevil genome possibly coming from *S. pierantonius*. Contrary to the genome of the tsetse fly *Glossina,* where at least three HGT events from *Wolbachia* have been reported (Initiative 2014), we were unable to pinpoint any HGT event from either the ancient endosymbiont *Nardonella, Wolbachia,* or the recently acquired *S. pierantonius.* A detailed description is reported in Supplemental file S1: Supplemental Note 1, and Note 3 for digestive enzymes.

#### Global analysis of metabolic pathways

Using the CycADS (Vellozo et al. 2011) pipeline and Pathway Tools (Karp et al. 2019) we have generated BioCyc metabolism reconstruction databases for *S. oryzae* and its endosymbiont *S. pierantonius*. We compared *S. oryzae* metabolism to that of other arthropods available in the ArthropodaCyc collection and we explored the metabolic exchanges between weevils and their endosymbionts (see Supplemental file S1: Supplemental Note 2). The metabolic reconstruction reveals that, despite its large genome for an endosymbiotic bacterium, *S. pierantonius* relies on its host for several central compounds, including alanine and proline, but also isocitrate, Inosine MonoPhosphate (IMP) and Uridine MonoPhosphate (UMP), to produce essential molecules to weevils, including the essential amino acids tryptophan, phenylalanine, lysine and arginine, the vitamins pantothenate, riboflavin and dihydropteroate as a folate precursor, as well as nicotinamide adenine dinucleotide (NAD) (Supplemental file S1: Supplemental Note 2). Among the amino acids listed above, phenylalanine, in particular is an essential precursor for the cuticle synthesis in emerging adults (Vigneron et al. 2014). In addition, several studies have shown that *S. pierantonius* improves host fitness, including fertility, developmental time and flight capacity, in part by supplying the host with vitamins and improving its mitochondrial energy metabolism (Heddi et al. 1999; Grenier et al. 1994; Rio et al. 2003).

#### Development

The annotation of developmental genes uncovered a high level of conservation in comparison to the red flour beetle *Tribolium castaneum*, a model coleopteran. When compared to *D. melanogaster*, several key coordinate group genes are absent in *T. castaneum* and *S. oryzae*. Moreover, a number of genes with two homologs in the *Drosophila* genome are represented by a single ortholog in *T. castaneum* and *S. oryzae*. We also observed that homologs for several signaling pathway ligands could not be identified, which, given the presence of conserved receptors, is probably due to divergent primary sequence of the ligands. A detailed description is reported in Supplemental file S1: Supplemental Note 4.

#### Cuticle protein genes

Among the distinctive biological features of coleopterans is the ability to generate a hard and thick cuticle that protects them against dehydration and represents the first physical barrier from infections and topical insecticide penetration. The analysis of cuticle proteins (CPs) showed that *S. oryzae* has an average number of CPs, but with an enrichment of members of the CPAP1 family. While some members of this family are known to be involved in molting and maintaining the integrity of the cuticle in *T. castaneum*, most are still uncharacterized (Jasrapuria et al. 2010, 2012). Thus, these proteins might be involved in the development of specific cuticular tissues in *S. oryzae* or other weevils. The total number of CPs did not follow the taxonomy of beetles, suggesting instead that it might be an adaptation to their diverse lifestyles. For details see Supplemental file S1: Supplemental Note 5.

#### Innate immune system

The analysis of immunity-related genes revealed that the genome of *S. oryzae* encodes the canonical genes involved in the three main antimicrobial pathways Toll, Imd and JAK-STAT, suggesting functional conservation of these pathways in cereal weevils. The conservation of the Imd pathway in the *S. oryzae* genome is of particular interest as its degradation in other symbiotic insects (*Acyrthosiphon pisum* (Gerardo et al. 2010), *B. tabaci* (Zhang et al. 2014), or *Rhodnius prolixus* (Salcedo-Porras et al. 2019) was initially correlated to their symbiotic status. The Imd pathway is not only present in *S. oryzae*, but it is also functional (Maire et al. 2018, 2019), and has evolved molecular features necessary for endosymbiont control (Maire et al. 2018) and host immune homeostasis (Maire et al. 2019). Thus, not only is the Imd pathway conserved in cereal weevils, contrary to aphids and some other hemimetabolous insects, but it seems to have been evolutionary “rewired” toward additional functions in symbiotic homeostasis (Maire et al. 2018). A detailed description can be seen in Supplemental file S1: Supplemental Note 6.

#### Detoxification and insecticide resistance

Fumigation using phosphine, hydrogen phosphide gas (PH_3_), is by far the most widely used treatment for the protection of stored grains against insect pests due to its ease of use, low cost, and universal acceptance as a residue-free treatment (Chaudhry 2000, 1997). However, high-level resistance to this fumigant has been reported in *S. oryzae* from different countries (Athié et al. 1998; Rajendran 1998; Zeng 1999; Benhalima et al. 2004; Pimentel et al. 2010; Nguyen et al. 2015, 2016; Holloway et al. 2016; Agrafioti et al. 2019). Hence, we searched for genes associated with detoxification and resistance to insecticide and more generally to toxins, including plant allelochemicals. The *S. oryzae* repertoire of detoxification and insecticide resistance genes includes more than 300 candidates, similar to what is seen in other coleopteran genomes. For more details see Supplemental file S1: Supplemental Note 7.

#### Odorant receptors

One promising pest management strategy relies on modifying insect behavior through the use of volatile organic compounds that act on odorant receptors (ORs) (Carey and Carlson 2011; Andersson and Newcomb 2017). ORs play a significant role in many crucial behaviors in insects by mediating host-seeking behavior, mating, oviposition, and predator avoidance (Leal 2013). Interfering with the behavior of pest insects and modulating their ability to find suitable hosts and mates has been shown to reduce population numbers, notably using plants that are capable of producing attractants and repellents (Hassanali et al. 2008; Hatano et al. 2015). *Sitophilus* spp. are known to use kairomones for host detection (Ukeh et al. 2012; Germinara et al. 2008), as well as aggregation pheromones (Phillips et al. 1985; Schmuff et al. 1984). We annotated 100 candidate OR genes in *S. oryzae* (named SoryORs), including the gene encoding the co-receptor Orco. Of these genes, 46 were predicted to encode a full-length sequence. The global size of the SoryOR gene repertoire is in the range of what has been described in other species of the coleopteran suborder Polyphaga (between 46 in *Agrilus planipennis* and more than 300 in *T. castaneum*) and close to the number of OR genes annotated in the closely related species *Dendroctonus ponderosae* (85 genes, (Mitchell et al. 2020)) (Supplemental file S1: Supplemental Note 8).

### Massive expansion of TE copies in the genome of *S. oryzae*

#### Detection and annotation of the repeatome

The repeatome represents the fraction of the genome categorized as repetitive. It encompasses TEs, satellites, tandem, and simple repeats. Eukaryotic TEs can be separated into two classes, depending on their replication mode (Makałowski et al. 2019). DNA (Class II) based elements are able to directly move within a genome, and include terminal inverted repeat (TIR), Crypton, rolling-circle (RC/Helitron), and large composite elements (Maverick). Conversely, retrotransposons (Class I) have an RNA intermediate and replicate through RNA retrotranscription. Retrotransposons can be further divided into long terminal repeat (LTR), and non-LTR elements, including long and short interspersed nuclear repeat elements (LINEs and SINEs). Other retrotransposons include Penelope-like (PLEs) and DIRS-like elements. Each one of these TE orders can be further classified into specific superfamilies (as for instance Copia or Gypsy LTR elements, and hAT or Tc1/Mariner TIR elements), that may encompass hundreds of TE families, each containing thousands of copies. The intrinsic diversity of TEs complicates their identification and annotation, especially in understudied species genera.

We used multiple state-of-the-art TE detection tools, including RepeatModeler2 and EDTA (Flynn et al. 2020; Ou et al. 2019), to generate consensus sequences of the TE families in *S. oryzae*. After an initial discovery step, more than 10 000 likely redundant TE families were identified by the dedicated programs; we combined their results using multiple sequence alignments and clustering (see Methods and Supplemental file S1: Figure S1) to reduce this number to 3 399. Due to the evolutionary distance between *S. oryzae* and other known coleopterans, the consensus sequences obtained were further classified using a thorough combination of sequence homology and structure (see Methods).

#### The *S. oryzae* genome is among the most TE-rich insect genomes to date

We uncovered 570 Mbp of repeat sequences, corresponding to ≈74% of the *S. oryzae* genome: ≈2% of satellite sequences, simple or low-complexity repeats, and ≈72% of other mobile elements, including TEs, (Figure 2A, Supplemental table S2). Given the limitation of the sequencing technologies, the proportion of satellites and TEs usually abundant in the heterochromatin is likely underestimated. We took advantage of a recent comparative analysis of TE content in 62 insect species (Petersen et al. 2019) to contrast with the *S. oryzae* TE compartment. The *S. oryzae* genome ranks among those with the highest TE fraction observed in insects (Figure 2B and 2C). Within the largest insect order, Coleoptera, very little is known regarding TE distribution and evolution. *T. castaneum* harbors only 6% of TEs (Tribolium Genome Sequencing Consortium et al. 2008), *Hypothenemus hampei* contains 8.2% of TEs (Vega et al. 2015; Hernandez-Hernandez et al. 2017), while *Dichotomius schiffleri* harbors 21% (Ic et al. 2020). The species closest to *S. oryzae*, *Rhynchophorus ferrugineus,* has a TE content of 45% (Hazzouri et al. 2020). Therefore, while TE content has been described to follow phylogenetic relationships in most insects (Gilbert et al. 2021; Sessegolo et al. 2016) there is a large variation among the few Coleoptera species with available genomes. It is important to note that the pipeline we used to detect and annotate TEs in *S. oryzae* differs from the method implemented by Petersen and colleagues (Petersen et al. 2019), as we incorporated 31 manually curated TE references for *S. oryzae*, and specifically annotated DNA/TIR elements based on their sequence structure (see Methods), increasing the annotation sensitivity.

**Figure 2.**
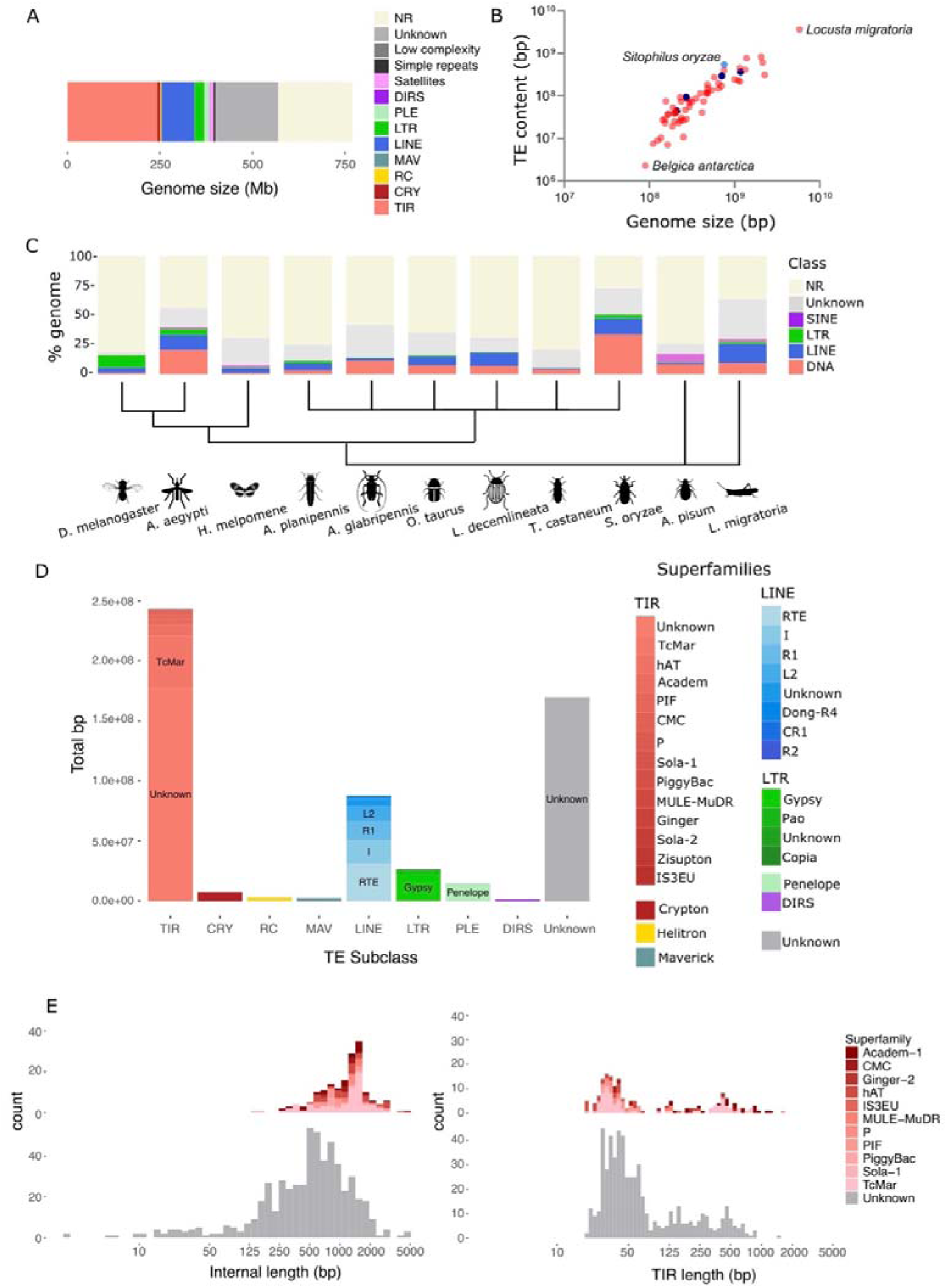
A. Proportion of repeat content in *S. oryzae*’s genome. The majority of repeats detected in *S. oryzae* are represented by Class II (TIR) elements, LINEs (Class I), and unclassified repeats (unknown). NR: non repetitive. B. Variation of genome size and TE content in 62 insect species from (Petersen et al. 2019) and *S. oryzae*. Coleopteran species are depicted in dark blue, and *S. oryzae* in light blue. *S. oryzae* is clearly a TE-rich genome. C. TE proportion across 11 insect species, including six coleoptera. In agreement with the data used for comparison (Petersen et al. 2019), PLEs are included in the LINE superfamilies, DIRS in LTRs, and RC, CRY, MAV and TIR in the DNA superfamilies. NR: non repetitive. *S. oryzae* harbors the largest TE content among Coleopterans and most insect species studied to date. Within Coleoptera, there is a large variation in TE content and type, with *A. planipennis, L. decemlineata* and *O. taurus* carrying an abundant LINE content, while *S. oryzae, T. castaneum* and *A. glabripennis* show larger DNA content. Cladogram based on (Misof et al. 2014). D. Classification of the 570 Mbs of TEs present in the *S. oryzae* genome. Most TIR families detected were not classified into known superfamilies. RTE LINE and Gypsy LTR elements are the most abundant superfamilies among retrotransposons. Around 22% of repeats in *S. oryzae’*s genome were not classified by our pipeline, and remain unknown (grey). E. Distribution of TIR length sequences (right) detected by einverted, and the internal region present between both TIRs (left) for complete consensus of TIR superfamilies (color) and unknown TIR families (grey).

#### Class II (DNA) elements dominate *S. oryzae*’s genome

The most striking feature of the genome of *S. oryzae* is the high abundance of Class II (DNA) elements (≈32% of the genome, ≈43% of the TE content) (Figure 2A), which is the highest observed among all 62 insect species included in this analysis (Petersen et al. 2019; Wang et al. 2014; Kelley et al. 2014). The most DNA transposon-rich genomes include mosquito *Culex quinquefasciatus,* and *Ae. aegypti,* harboring 25% and 20% of DNA transposon content in their genome, amounting to 54% and 36% of the total TE compartment, respectively (Vega et al. 2015). The TE-rich grasshopper *L. migratoria* repeatome comprises only 14% of DNA transposons, while LINE retroelements (Class I) amount to 25%. Morabine grasshoppers, with up to 75% of TE content, show equivalent amounts of DNA, LINE and Helitrons (Palacios-Gimenez et al. 2020). Finally, among Coleoptera, a large diversity of repeatomes is observed (Figure 2C) with *A. planipennis, Leptinotarsa decemlineata* and *Onthophagus taurus* carrying an abundant LINE content, while *S. oryzae, T. castaneum* and *Anoplophora glabripennis* show larger DNA transposon content.

Among the Class II elements present in *S. oryzae*, the majority belongs to the TIR subclass but has not been assigned a known superfamily (Figure 2D), while Tc Mariners make up ≈6% of DNA elements. Among the consensus sequences we were able to assemble from 5’TIR to 3’TIR (highest confidence, see Methods), the length distribution shows a continuum starting at a couple of hundred bases to a maximum of ∼5 Kbp (see Figure 2E). We hypothesize that most of the smaller TIR families observed are miniature inverted repeat elements (MITEs). MITEs are non-autonomous elements, deriving from autonomous Class II/TIR copies, comprising two TIRs flanking a unique, non-coding, region (sometimes absent) of variable length. While the TE detection pipeline used was able to detect and annotate most Class II/TIR elements based on transposase homologies, we also specifically searched for non-autonomous TIR sequences, allowing the detection of putative MITEs that lack protein coding regions (Supplemental file S1: Figure S1). Among all Class II/TIR superfamilies, TIR length varies between tens of base pairs to ≈1 Kbp (Figure 2E). We identified short elements, composed mostly of their TIR sequences (Figure 2E), typical of MITEs. Interestingly, the unknown TIR families show an average size smaller than 1 Kbp, while TIRs with an annotated superfamily, show larger sizes (Supplemental file S1: Figure S3), suggesting that most unknown families could be indeed non-autonomous MITEs. MITE size ranges were previously described from around 100 bp to copies reaching more than 1 Kbp (Feschotte et al. 2002). Finally, the distribution of the proportions of TIR relative to the consensus length appears superfamily-specific (Figure 2E and Supplemental file S1: Figure S3), and unknown families recapitulate these patterns. In conclusion, while most unknown TIR families seem to be composed of MITEs, we cannot exclude that our homology database is limited, likely missing some unknown protein domains. The most abundant TE family detected in the *S. oryzae* genome is indeed a MITE element (TE2641_SO2_FAM0704), with 10 486 genomic hits (or the equivalent of ≈4 117 copies based on the consensus size), corresponding to 1.3% of the genome. Large fractions of MITEs were also reported in Class II-rich genomes, such as the aforementioned mosquitoes (Nene et al. 2007; Feschotte and Mouchès 2000) and the invasive *Ae. albopictus* (Goubert et al. 2015), but also in many plant species such as the rice *Oryza sativa* (Feschotte et al. 2003; Lu et al. 2012; Feng 2003). Among Class II elements, we have also detected Crypton (0.9% of the genome), RC/Helitrons (0.4% of the genome) and Mavericks (0.3% of the genome).

LINE elements are the second most abundant TE subclass, representing ≈11% of the *S. oryzae* genome, among which ≈35% are assigned to RTE elements and ≈22% to I elements (Figure 2D). No SINE families have been detected. LTRs are rather scarce, representing only ≈3% of the genome (Figure 2D), and the vast majority belongs to the Gypsy superfamily (≈30%). Another retrotransposon order detected are Penelope (PLEs), reaching nearly 2% of *S. oryzae*’s genome, and DIRS (Tyrosine recombinase retrotransposons, 0.14% of the genome).

Finally, around 22% of the genome is composed of repeats for which our pipeline could not assign a known TE class (Figure 2D). These unknown families highlight the wealth and diversity of TEs among insects and Coleopteran genomes in particular, and could represent an overlooked reservoir of genomic innovations.

#### TE copies make up most of non-coding sequences of *S. oryzae*’s genome

TE copies are interspersed around the *S. oryzae* genome. TEs are less frequently found close to gene transcription start sites (TSS), 5’ and 3’ untranslated regions (5’ and 3’ UTRs) and exons (Figure 3A), as expected. On the contrary, introns and intergenic sequences harbor the highest TE content (Figure 3A), amounting to around 50% of TE density, close to the general TE proportion in the genome (72%), suggesting that most non-coding DNA sequences in the *S. oryzae* genome are virtually made of TEs. To grasp the impact of TEs on intron size, we compared intron length in *S. oryzae* with two very well assembled genomes: *D. melanogaster* with a very compact and small genome, and the large, TE-rich human genome (Figure 3B). In *D. melanogaster*, introns are small and harbor few TEs, while in humans, introns are much larger potentially due to high TE accumulation (Sela et al. 2010). *S. oryzae* intron sizes also seem to be due, at least partly, to TE accumulation. Interestingly, the *S. oryzae* genome presents a bimodal distribution, with a large proportion of small introns, as found in *D. melanogaster,* but also a noticeable amount of larger, TE packed and more human-like introns. This could suggest that specific regions of the genome could be more prone to TE elimination, and be associated with high rates of recombination and/or signature of purifying selection.

**Figure 3.**
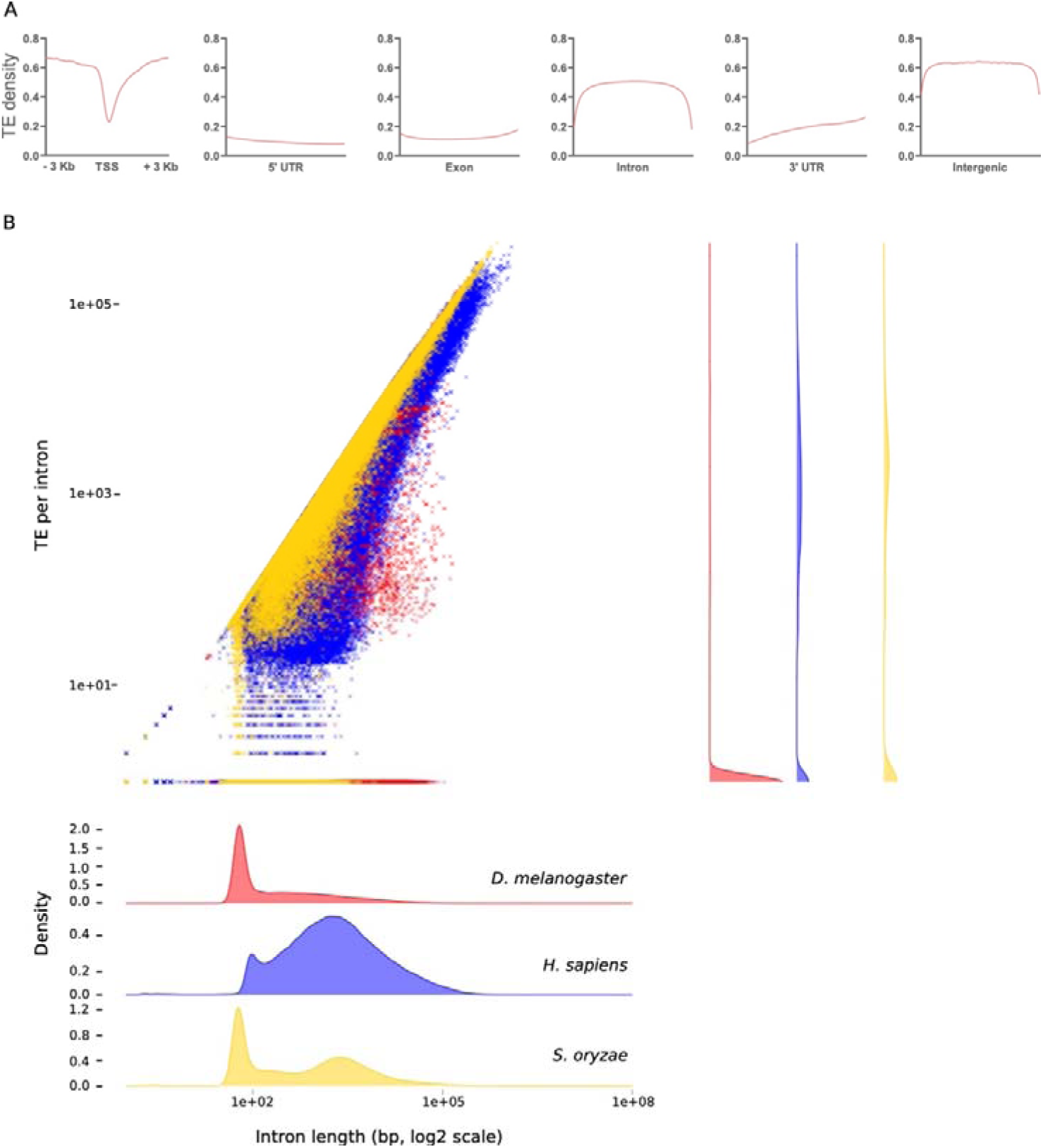
TE distribution in *S. oryzae*’s genome. A. Density of TE copies within gene regions. TE copies are the least abundant within TSSs, 5’ and 3’ UTRs and exons, while introns and intergenic regions are riddled with TEs. TSS: transcription start site, UTR: untranslated regions. B. Relationship between intron length and TE per intron in *D. melanogaster* (red), *H. sapiens* (blue) and *S. oryzae* (yellow). *S. oryzae* shares characteristics of both *Drosophila* with short and TE poor introns and Humans with a significant number of large, TE-packed introns.

#### TE activity inferred by evolutionary history

Within reconstructed TE families, nucleotide substitution levels (Kimura 2 Parameters, K2P) between copies and their consensus sequences allowed estimation of their relative ages and identified potentially active ones (Figure 4A). Such “TE landscapes’’ are extremely helpful to pinpoint potential TE amplifications (modes in the distribution) and extinctions (valleys) within the 0-30% K2P range (beyond, the increased divergence between copies affects negatively the sensitivity of the alignments, such that TE-derived sequences are no longer recognisable). The landscape analysis revealed a heterogeneous distribution of the TE copy divergence to their consensus within and between the main TE subclasses (Figure 4A). Most identified TE copies have a K2P divergence under 10, which is often observed in insects, and strikingly distinguishes itself from TE-rich mammalian genomes (RepeatMasker.org, (Petersen et al. 2019)). While *S. oryzae’*s TE density and distribution evoques the architecture of mammalian genomes, this relatively younger TE landscape suggests higher deletion rates, and possibly a higher TE turnover rate, as observed in *Drosophila* (Petrov 2002; Petrov and Hartl 1998). LINEs and DNA transposons have the wider spectrum of divergence levels, suggesting an aggregation of distinct dynamics for the TE families present in *S. oryzae.* By contrast, the rare LTR copies identified appear to be the most homogeneous within families, with only a few substitutions between copies and their consensuses, suggesting a very recent amplification in this subclass. Finally, unknown TEs share a large part of their K2P distribution with TIR elements, though relatively less divergent from their consensus sequences as a whole. A breakdown of the K2P distributions at the superfamily level reveals specific evolutionary dynamics (Figure 4B). Diverse superfamilies, such as Tc-Mar and hAT (TIR) or RTE (LINE), show more uniform distributions, suggesting sustained activity of some of its members throughout *S. oryzae*’s genome evolution, though this could also indicate that these subfamilies could be subdivided further. As observed at the class level, all three identified LTR superfamilies (Pao, Gypsy and Copia) show families within the lowest K2P range.

**Figure 4.**
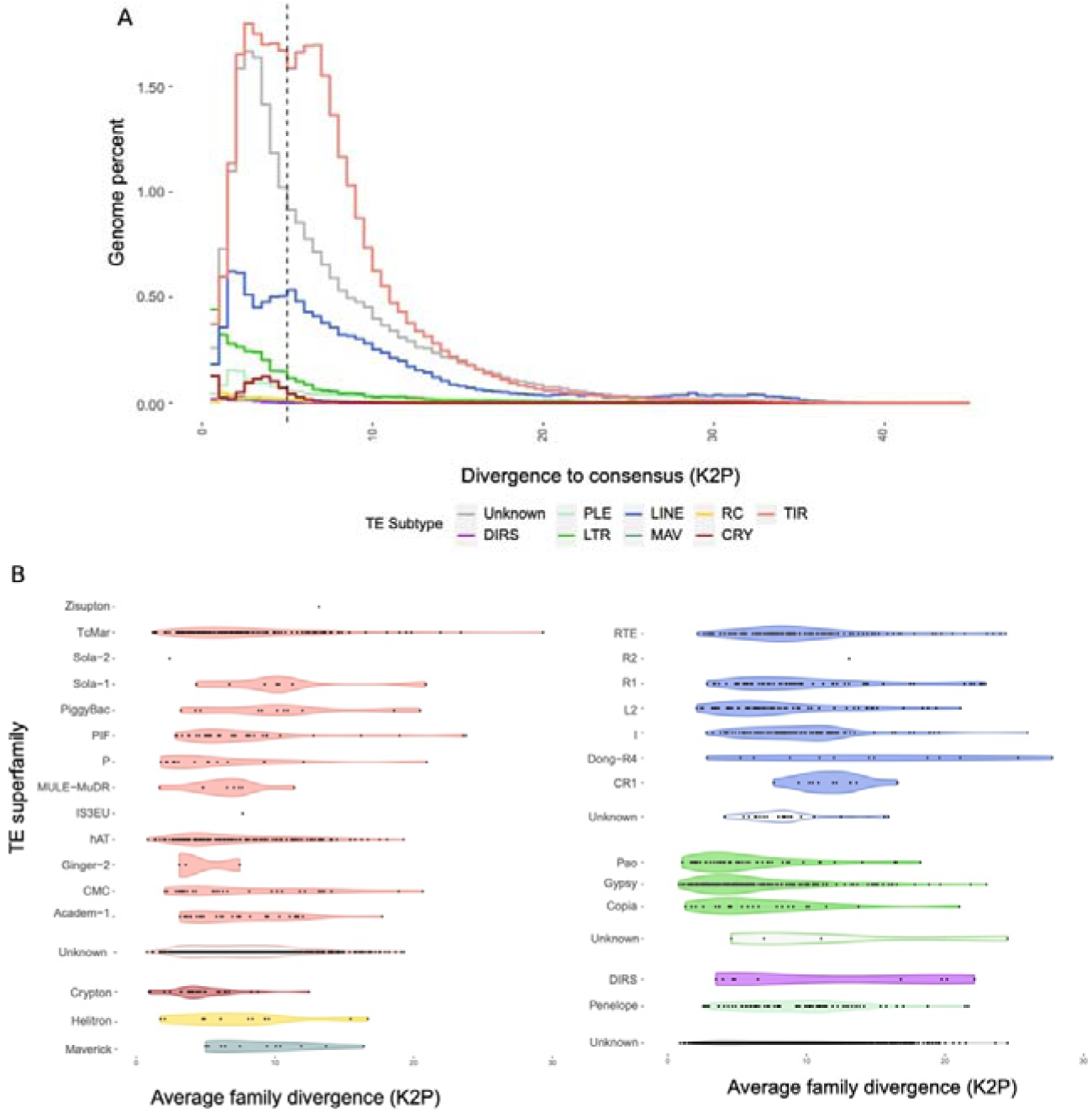
A. TE divergence landscape. Distribution of the divergence (Kimura two parameters, K2P) between TE copies and their consensus, aggregated by TE class reported in percent of the genome. The less divergent superfamilies are distributed to the left and suggest recent activity. Strikingly, most of the TE copies have less than 10% divergence to their consensus, with a large number of copies under 5% (dotted line). The distribution of the “unknown” class overlaps with the leftmost mode of the TIR distribution, suggesting that many more TIR families are yet to be described in *S. oryzae.* Strikingly, LTR elements are the least diverged altogether with the mode of the distribution on the 0-1% divergence bin. B. Mean K2P distributions within TE superfamilies. Left panel depicts Class II families, and all Class I (retrotransposons) and unknown families are on the right panel. LTR superfamilies harbor some of the least divergent TE families, suggesting that this class may host some of the youngest TE.

#### TEs are transcriptionally active in somatic and germline tissues

The TE K2P landscape suggests that LTR elements as well as some LINE families and several Class II subclasses are among the youngest, and thus potentially active. In order to estimate the transcriptional activity of *S. oryzae*’s TE families, we have produced somatic (midgut) and germline (ovary) transcriptomic data. While germline tissues allow identification of potential TE families capable of producing vertically transmitted new copies, TE derepression in somatic tissues represents the potential mutational burden due to TEs. The expression of TE families varied extensively within a class and the proportion of transcriptionally active/inactive TE families between classes was also distinct (Figure 5A). In total, 1 594 TE families were differentially expressed between ovary and midgut tissues (Figure 5B, Supplemental table S3); of which 329 have an absolute Log2 fold change higher than 2 (71 downregulated and 258 upregulated in midgut). In total, we detected 360 TE families downregulated in midgut when compared to ovaries: A much larger set of upregulated TE families was detected in midgut when compared to ovaries (1 236), illustrating the tighter regulation of TE copies in germline tissues. Moreover, the distribution of Log2 fold changes were similar between TE subclasses but different for LTRs, which had a higher proportion of upregulated TE families in ovaries compared to other classes (Figure 5C. Kruskall and Wallis rank-sum test: H = 36.18, *P* < 0.01; LTR vs. LINE, Class II or Unknown: Dunn’s test: *P-adj* < 0.01). In conclusion, the large TE compartment in *S. oryzae* shows abundantly expressed TE families, and tissue-specific expression patterns.

**Figure 5.**
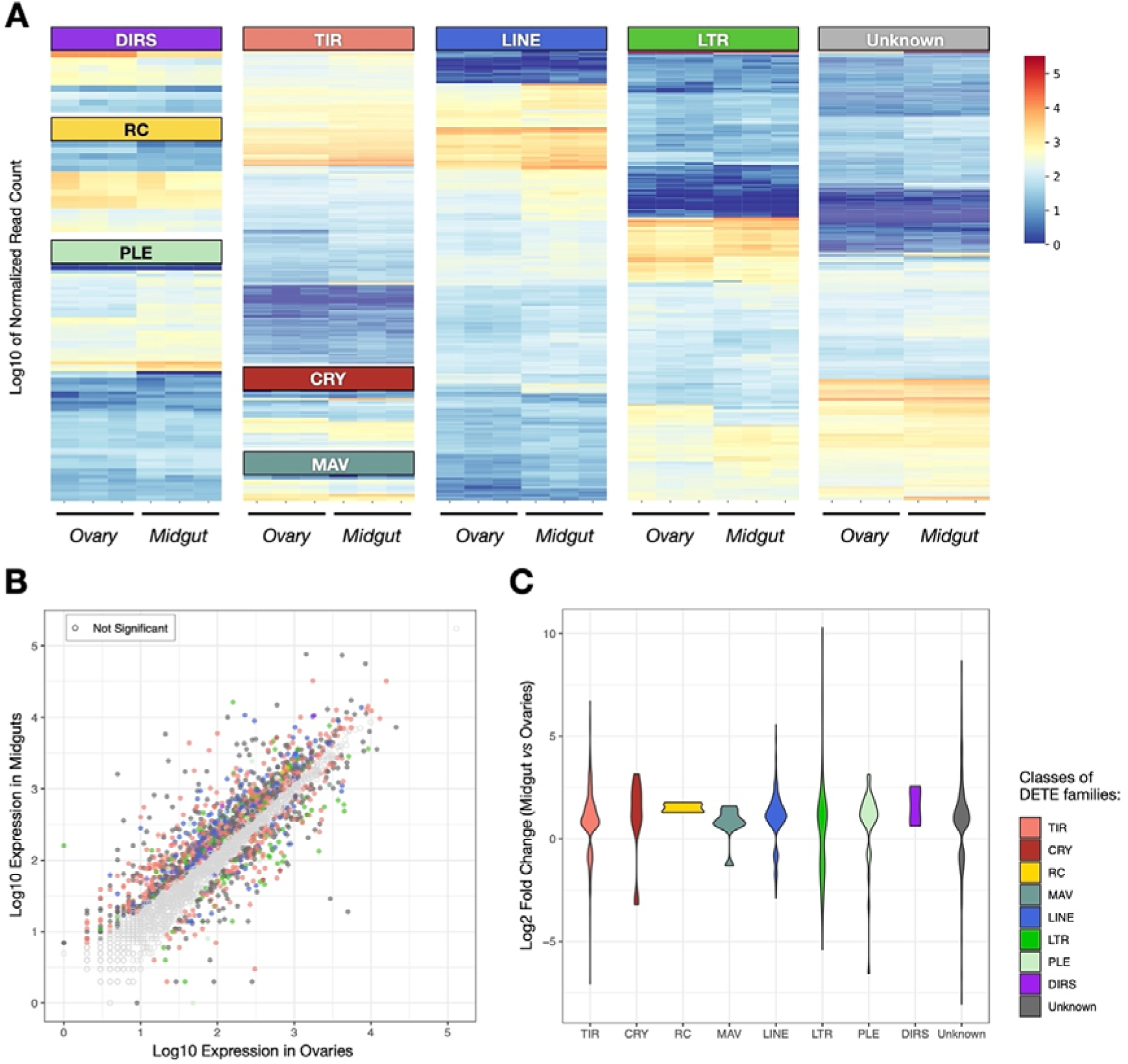
TE family expression in midguts and ovaries from *S. oryzae*. A. Log10 normalized counts in midguts and ovaries triplicates. Normalized counts show different proportions of transcriptionally active TE families in different TE classes. B. Log10 of base mean average expression of TE families in ovaries and midguts from three biological replicates. Depicted in color only TE families which had differential expression between ovary and gut tissues (padj<0.05, |log2FC|>2). Most TE families are upregulated in midguts compared to ovaries. C. Distribution of all significant (padj<0.05). Log2FC depicts specifically deregulated TE classes in each tissue. LTR elements are predominantly upregulated in ovaries.

To estimate the TE transcriptional load imposed on *S. oryzae,* we computed the percentage of total RNA-seq poly-A enriched reads mapping to TE consensus sequences (Supplemental file S1: Figure S4). Around 5% of the midgut transcriptome corresponds to TE sequences. We compared such transcriptional burden to a TE-poor (*D. melanogaster,* ≈12%) and a TE-rich (*Ae. albopictus* ≈50%) genome, using similar technology in equivalent tissues (adult midgut, see Methods). It is important to note that, despite being a TE-poor genome, *D. melanogaster* harbors many young LTR elements that have been recurrently shown to transpose (Pasyukova and Nuzhdin 1993). We did not detect a direct correlation between genomic TE content and TE expression (Supplemental file S1: Figure S4). *S. oryzae* bears the highest proportion of RNAseq reads mapped against TE consensus sequences (≈5%), followed by *D. melanogaster* (≈1%) and *Ae. albopictus* (≈0.01%). Henceforth, not only is *S. oryzae* a TE-rich genome, but the transcriptional load from TEs is higher than in other TE-rich genomes (*Ae. albopictus)*, and in genomes harboring young and active TE copies (*D. melanogaster,* (Adams et al. 2000; Ashburner and Bergman 2005)).

Finally, it is important to note that while transcriptional activation of TE copies may have an impact on the host genome, it does not indicate high transposition and therefore higher mutation rates. The high transcriptional load of *S. oryzae* compared to other species might stem from differences in TE regulation. In insects, TEs are mainly silenced by small RNAs and repressive chromatin marks (Czech and Hannon 2016). More specifically, piwi-interacting RNAs (piRNAs) are able to target post-transcriptional repression of TEs, and guide chromatin silencing complexes to TE copies (Sienski et al. 2012; Andersen et al. 2017; Czech and Hannon 2016). Therefore, we have annotated genes implicated in small RNA biogenesis and found that all three pathways (piRNAs but also microRNAs and small interfering RNAs biogenesis pathways) are complete (Supplemental file S1: Supplemental Note 9). Genes involved in piRNA biosynthesis are expressed mainly in ovaries and testes, while somatic tissues (midgut) show smaller steady-state levels (Supplemental file S1: Supplemental Note 9), suggesting the piRNA pathway is potentially functional in *S. oryzae* ovaries, and could efficiently reduce transposition.

#### TE content is variable among *Sitophilus* species

Cereal weevils are part of the Dryophthoridae family that includes more than 500 species. Very little is known about genome dynamics in this massive phylogenetic group. Because of the unusual high TE copy number and landscape observed in *S. oryzae,* we analyzed three other closely related species namely *Sitophilus zeamais, Sitophilus granarius* and *Sitophilus linearis*. We produced low coverage sequencing and estimated the TE content from raw reads using our annotated *S. oryzae* TE library with dnaPipeTE (Goubert et al. 2015). Remarkably, among *Sitophilus* species, repeat content is variable (Figure 6A), with *S. linearis* harboring the smaller repeat load (≈54%) compared to *S. oryzae* (≈80%), *S. zeamais* (≈79%), and *S. granarius* (≈65%). Most importantly, Class II (DNA) elements of *S. oryzae* are nearly absent from *S. linearis*, and no recent burst of LTR elements is observed, contrary to the other *Sitophilus* species, suggesting alternative TE evolutionary histories (Figure 6B). It is important to note that our analysis is biased towards *S. oryzae,* as the library used to annotate the TEs in the other *Sitophilus* species stems from automatic and manual annotation of the *S. oryzae* genome. Finally, the relatively higher dnaPipeTE estimations of the LTR content in *S. oryzae* compared to the assembled genome supports the hypothesis that LTR elements have seen a recent burst of transposition, as young elements tend to collapse in genome assemblies and eventually diminish their estimated copy number.

**Figure 6.**
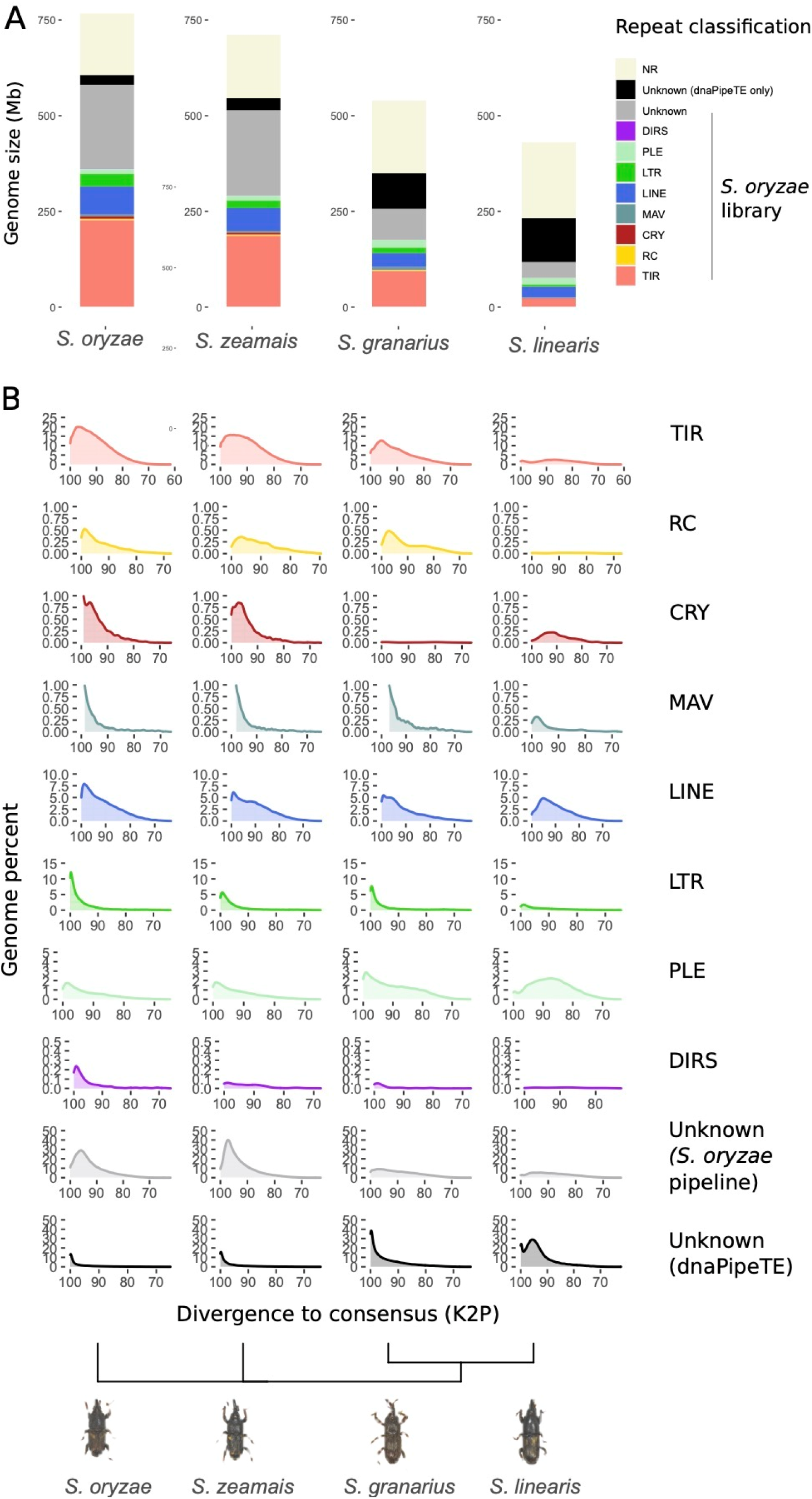
TE landscape across *Sitophilus* species. A. Proportion of TE per species estimated from short reads with dnaPipeTE and a custom TE library including Repbase (release 2017) and annotated TE consensus discovered in *S. oryzae*. *S. oryzae*, *S. zeamais* and *S. granarius* harbor similar TE content, while *S. granarius* presents a smaller TE load, and *S. linearis* harbors the smallest TE content and the higher proportion of unknown repeats. The proportion of unknown repeats only found by dnaPipeTE (black) increases from *S. oryzae* to *S. linearis* with the phylogenetic distance. B. Distribution of divergence values between raw reads and repeats contig assembled with dnaPipeTE (blastn) across four *Sitophilus* species. *S. oryzae* appears to share its TE landscape with *S. zeamais* and *S. granarius*, but the three species display a distinct repeatome than *S. linearis,* in spite of their phylogenetic proximity. SO2: *S. oryzae*’s TE library produced in this analysis, DPTE: DNApipeTE TE annotation (repeats only found by dnaPipeTE).

Overall, the comparison of TE content in closely related species highlights the influence of phylogenetic inertia, but reveals a possible TE turnover in the *S. linearis* lineage. In addition to the regulation mechanisms that strongly contribute to TE amount and variation, TE accumulation is conditioned by the drift/selection balance in populations. Indeed, effective population size has been suggested to be a major variable influencing TE content, as small, inbred or expanding populations suffer drift, allowing detrimental insertions to stay in the gene pool and thus favor TE fixation (Lynch and Conery 2003). Such hypotheses should be addressed in the future, especially on recently sequenced TE-rich but rather small (<1 Gbp) genomes such as *S. oryzae*.

#### Endosymbionts impact TE transcriptional regulation

The four *Sitophilus* species studied have different ecologies. *S. oryzae* and *S. zeamais* infest field cereals and silos, while *S. granarius* is mainly observed in cereal-containing silos. *S. linearis,* however, lives in a richer environment, *i.e.* tamarind seeds. In association with their diets, the interaction of *Sitophilus* species with endosymbiotic bacteria differs: the cereal weevils (*S. oryzae, S. zeamais* and *S. granarius*) harbor the intracellular gram-negative bacteria *S. pierantonius*, albeit at very different loads. While *S. oryzae* and *S. zeamais* show high bacterial load, *S. granarius* has a smaller bacterial population (Vigneron et al. 2014). In contrast, *S. linearis* has no nutritional endosymbionts, in correlation with its richer diet. We wondered whether the presence of intracellular bacteria impacts TE regulation, and took advantage of artificially obtained aposymbiotic *S. oryzae* animals to search for TE families differentially expressed in symbiotic versus aposymbiotic ovaries. There were 50 TE families upregulated in symbiotic ovaries compared to artificially obtained aposymbiotic ones, while 15 families were downregulated (Figure 7 and Supplemental table S3). Only three families presented an absolute Log2 fold change higher than 2: one LINE and two LTR/Gypsy elements. The three of them were upregulated both in symbiotic *versus* aposymbiotic ovaries, and in ovaries *versus* midgut (Supplemental table S3), suggesting that such elements have tissue specificity, and their expression is modulated by the presence of intracellular bacteria. Such TE families would be ideal candidates to further study the crosstalk between host genes, intracellular bacteria and TE transcriptional regulation.

**Figure 7.**
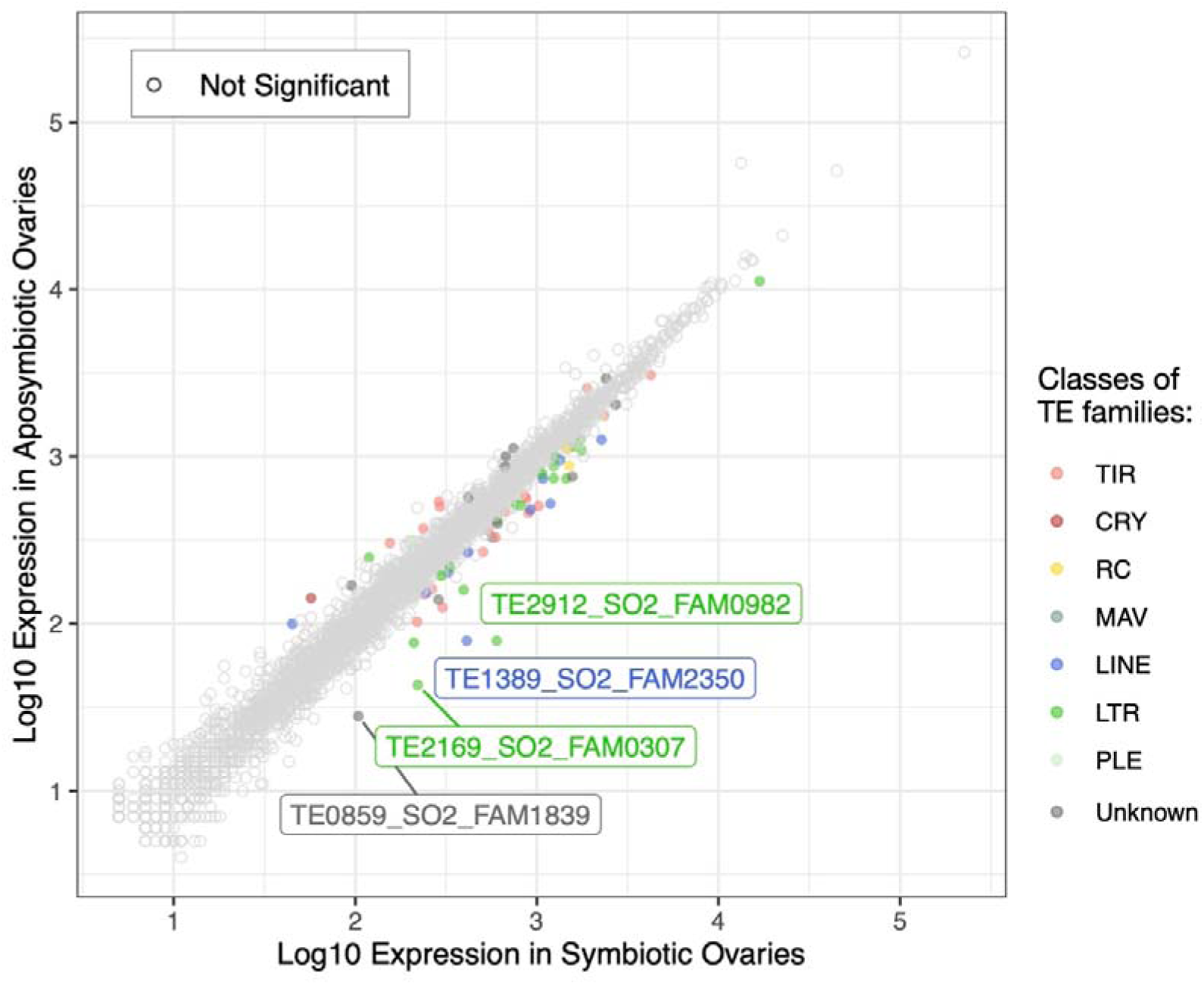
Differentially expressed TE families between symbiotic and aposymbiotic *S. oryzae* ovaries. Log10 of base mean average expression of TE families in symbiotic vs aposymbiotic ovaries from two biological replicates. Depicted in color only TE families which had differential expression between both ovary types (padj<0.05, |log2FC|>2). Two LTR elements and one LINE element are upregulated (log2FC > 2) in symbiotic ovaries.

## Conclusion

The success of obtaining a TE-rich genome assembly complete enough to understand genome architecture and regulatory networks relies on the use of multiple sequencing platforms (Peona et al. 2021). Here, we describe the first assembly of the repeat-rich (74%) *S. oryzae* genome, based on a combination of long and short read sequencing, and a new assembly method, WENGAN (Di Genova et al. 2020). While this first assembly reaches quality standards similar to other coleopteran species (Table 1), it is important to stress that new sequencing methods have emerged in order to improve genome assemblies, including linked-reads and optical mapping (Peona et al. 2021).

We uncovered around 74% of repeated sequences in the *S. oryzae* genome, mostly TE families. While the TE landscape is marked by a wealth of Class II elements, e specially non-autonomous MITE elements, 22% of the genome is composed of unknown repeats. Large duplicated gene families can be present in such a category, but it is tempting to speculate that the majority is composed of novel Class II elements. Indeed, Unknown and TIR elements share the same K2P landscapes, and many Class II elements have only been detected through an inverted repeat search for TIRs, and not proteins, excluding therefore TE copies old enough that TIRs are too divergent to be recognized. Moreover, we have shown that many TE families in *S. oryzae* are present in the transcriptome, suggesting that several families can be transcriptionally active. How such TE families are able to escape host silencing remains unknown. It seems obvious today that insect models such as *D. melanogaster* only represent a small window on the complex biology and evolution of TEs, and the sequencing and annotation of species with high TE content -- while challenging (Platt et al. 2016) -- is key to understanding how genomes, their size, their structure and their function evolve. In conclusion, *S. oryzae* constitutes an excellent model to understand TE dynamics and regulation and the impact on genome function.

*Sitophilus* species not only differ in their TE landscape, but also in their ecology and as a consequence, their association with intracellular bacteria. Comparison of TE content within the *Sitophilus* genus shows variable TE amount and diversity. In addition, intracellular bacterium impacts transcription of specific TE families in ovaries. The molecular mechanisms behind the co-evolution between an insect, its endosymbiotic bacterium and TEs remains unexplored. The impact of intracellular bacteria on host genomes is poorly studied, and the *Sitophilus* genus offers a simpler experimental setting, with a single intracellular bacterium present within specific host cells (Lefèvre et al. 2004; Heddi et al. 1999), and a well established knowledge of host-bacteria interaction (Vigneron et al. 2014; Maire et al. 2018, 2019, 2020; Login et al. 2011).

## Methods

### DNA extraction and high-throughput sequencing

Individuals of both sexes of *S. oryzae* were reared on wheat grains at 27.5 °C with 70% relative humidity. The aposymbiotic strain was obtained by treating the symbiotic strain during one month at 35 °C and 90% relative humidity as previously described (Nardon 1973). This strain is viable, fertile and was raised in the same conditions as the symbiotic strain. The aposymbiotic status was confirmed by PCR and histology. Male and female adults of *S. oryzae* were used for DNA extraction. Only the gonads were used to minimize DNA contamination from its diet, which could be still present in the gut. The reproductive organs were obtained from aposymbiotic adults and a DNA extraction protocol specific for *Sitophilus* weevils was performed. DNA extractions were performed using a STE buffer (100 mM NaCl, 1 mM Na_2_EDTA pH 8, 10 mM Tris HCl pH 8). Tissues were homogenized in STE buffer, then treated successively by SDS 10%, proteinase K and RNase. Briefly, genomic DNA was purified by two successive extractions with phenol:chloroform:isoamyl alcohol (25/24/1) followed by extraction with 1 vol of chloroform:isoamyl alcohol (24/1). Genomic DNA was then precipitated by 0.7 vol isopropanol. After washing the pellet with 70% ethanol, genomic DNA was recovered in TE (1 mM EDTA, 10 mM Tris HCl pH8) buffer. Using this protocol, we obtained six different DNA samples: four from males and two from females. Each sample corresponds to the genomic DNA from 20 individuals. Five additional DNA samples were obtained using a high molecular weight DNA extraction protocol consisting of a single phenol:chloroform:isoamyl alcohol (25/24/1) extraction step from the genomic DNA of 100 males. The DNA concentration in each of these samples was quantified using a NanoDrop spectrophotometer (ThermoFisher Scientific, Waltham, MA, USA).

Sequencing was performed using a combination of Illumina, PacBio and Nanopore technologies (Supplemental table S4). For each sex, two Illumina libraries were generated: one paired-end library with an average fragment size of 500 bp and one mate pair library with an average fragment size of 5 Kbp. The libraries were sequenced using an Illumina HiSeq 2000 platform with the V3 chemistry and a read size of 101 bp; the paired-end (PE) libraries were sequenced at the “Génomique & Microgénomique” service from ProfileXpert (Lyon, France) while the mate paired (MP) were sequenced at Macrogen (Seoul, South Korea). Two male samples were used to build (i) an Illumina library with an average fragment size of 200 bp which was sequenced on a HiSeq 2500 instrument using the V4 chemistry and a read size of 125 bp, and (ii) a PacBio library sequenced on seven SMRT cells using the P6-C4 chemistry. These two libraries were sequenced at KeyGene (Wageningen, The Netherlands). Finally, five male samples were used to build Nanopore libraries with the SQK-LSK109 kit and without DNA fragmentation step. The libraries were independently sequenced on five MinION R9.4 flow cells. These libraries were built and sequenced at the sequencing platform of the IGFL (Institut de Génomique Fonctionnelle de Lyon, Ecole Normale Supérieure de Lyon, France). Statistics and accession numbers from all the sequencing runs are listed in the Supplemental table S3.

### Genome assembly and annotation

First, the Illumina reads were error-corrected using BFC release 181 (Li 2015). The PacBio and Nanopore reads were error-corrected using LORDEC v0.9 (Salmela and Rivals 2014) with the error-corrected Illumina overlapping PE reads, a k-mer size of 19 and solidity threshold of 3. Overlapping reads were then merged using FLASH2 v2.2 (Magoc and Salzberg 2011). Based on the merged Illumina reads, a first short-read assembly was produced using a modified version of MINIA v3.2.1 (Chikhi and Rizk 2012) with a k-mer length of 211. A hybrid assembly was then performed using WENGAN v0.1 (Di Genova et al. 2020) on the MINIA short-read assembly and the raw Nanopore reads. The resulting assembly was polished using two rounds of PILON v1.23 (Walker et al. 2014) using the error-corrected Illumina overlapping PE reads and the --diploid option. A first scaffolding was then performed with two rounds of FAST-SG v06/2019 (Di Genova et al. 2018) and SCAFFMATCH v0.9 (Mandric and Zelikovsky 2015) with the error-corrected Illumina MP, Illumina PE, PacBio and Nanopore libraries. The LR_GAPCLOSER algorithm v06/2019 (Xu et al. 2019) was used for the gap-filling step using the error-corrected PacBio and Nanopore libraries. An additional scaffolding step was performed using RASCAF v1.0.2 (Song et al. 2016) with the available RNA-seq libraries from the Sequence Read Archive (SRX1034967-SRX1034972 and SRX3721133-SRX3721138). The resulting scaffolds were then gap-filled using a new round of LR_GAPCLOSER as previously described followed by two rounds of SEALER v2.1.5 (Paulino et al. 2015) using the error-corrected Illumina overlapping PE reads and k-mer sizes of 64 and 96. Two rounds of PILON, as previously described, were performed to produce the final assembly. Quality of the assembly was assessed by computing several metrics using i) QUAST v5.0.2 (Mikheenko et al. 2018) with a minimal contig size of 100 bp and the --large and -k options, ii) BUSCO v4.0.5 (Seppey et al. 2019) using the Insecta ODB10 database and the -geno option, and iii) KMC v3.0.0 (Kokot et al. 2017) to evaluate the percentage of shared 100-mers between the assembly and the merged Illumina reads.

Three contaminant scaffolds corresponding to the mitochondrial genome and an artefact were removed from the assembly prior to the annotation step. The ‘NCBI *Sitophilus oryzae* Annotation Release 100’ was produced using the NCBI Eukaryotic Genome Annotation Pipeline v8.2.

### Low-coverage genome sequencing of other *Sitophilus* species

Twenty pairs of ovaries were dissected from *S. oryzae, S. zeamais, S. granarius and S. linearis* females. Ovaries were homogenized in 100 mM NaCl, 1 mM EDTA pH 8, 10 mM Tris-HCl pH 8 using a small piston. Proteinase K digestion followed in the presence of SDS for 2 h at 55 °C with shaking and for 1 h at 37 °C with RNAse A. A typical phenol chloroform extraction was then performed and genomic DNA was isopropanol precipitated. Eight whole genome sequencing libraries with a median insert size of 550 bp were constructed using the Illumina TruSeq DNA PCR-free sample preparation kit (Illumina, San Diego, CA, USA), according to manufacturer’s protocols. Briefly, 2 µg of each gDNA were sheared using a Covaris M220 Focused-ultrasonicator (Covaris, Inc. Woburn, MA, USA), end-repaired, A-tailed, and adapter ligated. Library quality control was performed using the 2100 Bioanalyzer System with the Agilent High Sensitivity DNA Kit (Agilent Technologies, Santa Clara, CA, USA). The libraries were individually quantified via qPCR using a KAPA Library Quantification Kits (Kapa Biosystems, Wilmington, MA, USA) for Illumina platforms, then they were pooled together in equimolar quantities and sequenced in a MiSeq sequencing system. 2×300 paired-end reads were obtained using a MiSeq Reagent Kits (600-cycles).

### TE library construction

In order to annotate the *S. oryzae* repeatome, we collected and combined cutting-edge bioinformatic tools to (i) create and (ii) classify a non-redundant library of repeated elements (Supplemental file S1: Figure S1). First, we separately ran RepeatModeler2 (Flynn et al. 2020) and EDTA (Ou et al. 2019) on the assembled genome. Together, these programs include most of the recent and long-trusted tools used to detect generic repeats, but also include specific modules, such as for LTR and TIR elements. Preliminary analyses of the *S. oryzae* genome with RepeatModeler1 (Smit et al. 2013) and dnaPipeTE (Goubert et al. 2015) suggested a rather large fraction of Class II DNA elements with terminal inverted repeats (TIRs). Thus, MITE-Tracker (Crescente et al. 2018) was incorporated in our pipeline and ran independently on the genome assembly using 1- and 2-Kbp size cutoffs to detect Class II elements harboring TIRs with high sensitivity. Following this initial step, 15 510 consensus sequences obtained from RM2, EDTA and the two runs of MITE-tracker were successively clustered using MAFFT (Katoh and Standley 2013), Mothur (Schloss et al. 2009), and Refiner (Smit et al. 2013) to reduce redundancy in the repeat library to a total of 2 754 consensus sequences (Supplemental file S1: Figure S1A, https://github.com/clemgoub/So2). Then, we inspected the quality of the raw library by calculating the genomic coverage of each consensus. We ran the library against the genome using RepeatMasker (52) and implemented a simple algorithm “TE-trimmer.sh” to trim or split a consensus sequence wherever the genomic support drops below 5% of the average consensus coverage (Supplemental file S1: Figure S1A, https://github.com/clemgoub/So2). To mitigate any redundancy generated by the splitting, the newly trimmed library was clustered before being re-quantified using RepeatMasker (Smit et al. 2013). At this step, we removed any consensus under 200 bp and represented by less than the equivalent of two full-length copies (in total bp). In addition, TAREAN (Novák et al. 2017) was used to detect and quantify candidate satellite repeats. We obtained an *ab-initio* repeat library of 3 950 consensus sequences automatically generated (Supplemental file S1: Figure S1A).

To refine and improve the quality of the TE consensus sequences, we then turned it over to DFAM (Storer et al. 2021) who processed the *ab initio* library. First, any sequences mostly composed of tandem repeats were removed using a custom script to remove any sequences that were greater than 80% masked and/or had a sequence less than 100 bp. To generate seed alignments for each consensus, the consensus sequences were used as a search library for RepeatMasker to collect interspersed repeats. Seed alignments in the form of stockholm files were generated using the RepeatMasker output. To extend potentially truncated elements, the instances in the stockholm file for each model were extended into neighboring flanking sequences until the alignment was below a threshold equivalent to ∼3 sequences in agreement. More specifically, all sequences are extended using full dynamic programming matrices using an improved affine gap penalty (default: -28 open, -6 extension) and a full substitution matrix (default: 20 percent divergence, 43% GC background). The termination of extension occurs when the improvement by adding a further column to the multiple alignment does not exceed 27 (with default scoring system). This is equivalent to a net gain of ∼3 sequences in agreement. Following extension, the new consensus were collected and consensus sequences greater than 80% similar for 80% of their length were considered duplicates and only one consensus was kept.

Upon completion, we used RepeatMasker to quantify the improved library. We selected the top 50 elements (by abundance in the genome) represented in each of the “LTR”, “LINE”, “Class II” and “Unknown” classes for manual inspection (these categories represent the 4 most abundant classes of repeats in the *S. oryzae* genome). While most consensus sequences where correctly extended and annotated (200) we noticed some cases of over-extension with LTR (consensus doubled in size) and flagged others with non-supported fragments for further trimming (Supplemental table S2 | tab 1). Once our quality check completed and the sequences curated, we removed fragments with 100% identity against a previously established consensus (Supplemental table S2 | tab 2). The final TE library contains 3 399 sequences to classify.

The classification of the final repeat library was done in successive rounds combining homology and structure methods (Supplemental file S1: Figure S1B). Before the final TE library was completed, we manually curated and annotated the sequences of 31 transposable elements and satellites among the most represented in *S. oryzae*. These high-confidence references were added to the default libraries used by the following programs and Repbase v.2017 (Bao et al. 2015). We searched for nucleotide homology using RepeatMasker (V.4.1.1 (Smit et al. 2013)) with -s “-slow” search settings. Best hits were chosen based on the highest score at the superfamily level allowing non-overlapping hits of related families to contribute to the same hit. In addition we used blastx (Camacho et al. 2009) to query each consensus against a curated collection of TE proteins (available with RepeatMasker), as well as those identified in the 31 manual consensus sequences. We kept the best protein hit based on the blastx score. Based on the 200 consensus sequences manually inspected (see above), we set a hit length / consensus size threshold of 0.08 (RepeatMasker) and 0.03 (blastx) to keep a hit. In our hands, these thresholds were conservative to automate the classification. As an alternate homology-based method, we also ran RepeatClassifier (RepeatModeler2). Finally, because DNA elements are often represented by non-autonomous copies (unidentifiable or absent transposase) we further used einverted to flag terminal inverted repeats located less than 100 bp of the ends of each sequence. The complete library of 3 399 consensus sequences was first annotated at the subclass level (see DFAM taxonomy: https://dfam.org/classification/tree) if two out of RepeatMasker, RepeatClassifier and blastx annotations agreed. Further, the same rule was applied for the superfamilies if possible. At this stage, consensus sequences without annotation by homology but with TIRs as flagged by einverted, were classified as TIR and all other sequences classified as Unknown. We further divided the subclass “DNA” into “MAV” (Mavericks), “RC” (Rolling circle/Helitron), “CRY” (Crypton) and “TIR” (terminal inverted repeats). Finally, the classifications automatically given as “Unknown” to 16/274 manually inspected consensus sequences were replaced to match the manually reported classification.

### Estimation of the repeat content

The total repeat content of the *S. oryzae* genome was analyzed using RepeatMasker (v.4.1.1) and our classified library of 3 399 consensus sequences and the following parameters: -s -gccalc -no_is-cutoff 200. The subsequent alignments were parsed with the script “parseRM.pl” (Kapusta and Suh 2017) https://github.com/4ureliek/Parsing-RepeatMasker-Outputs) to remove hits overlap and statistically analyzed with R version 4.0.2.

### Genomic distribution of TE copies

The distribution of TE copies across the *S. oryzae* genome was assessed using two different approaches over six different genomic regions namely TSS ± 3 Kbp, 5’ UTRs, exons, introns, 3’ UTRs and intergenic regions. Briefly, the coverage of all TE copies was computed over a sliding window of 100 bp across the whole genome sequence using the makewindows and coverage tools from the bedtools package (Quinlan and Hall 2010) and the bedGraphToBigWig UCSC gtfToGenePred tool. Then the different genomic regions were retrieved from the *S. oryzae* annotation file (GFF format) using the gencode_regions script (https://github.com/saketkc/gencode_regions) and the UCSC gtfToGenePred tool (https://github.com/ENCODE-DCC/kentUtils). A matrix containing the TE coverage per genomic region was generated using the computeMatrix tool from deepTools (Ramírez et al. 2016) and used to generate metaplots using the plotProfile tool.

### TE landscapes

The relative age of the different TE families identified in the genome assembly was drawn performing a “TE-landscape” analysis on the RepeatMasker outputs. Briefly, the different copies of one TE family identified by RepeatMasker are compared to their consensus sequence and the divergence (Kimura substitution level, CpG adjusted, see RepeatMasker webpage: http://repeatmasker.org/webrepeatmaskerhelp.html) is calculated. The TE landscape consists of the distribution of these divergence levels. In the end, the relative age of a TE family can be seen as its distribution within the landscape graph: “older” TE families tend to have wider and flatter distribution spreading to the right (higher substitution levels) than the “recent” TE families, which are found on the left of the graph and have a narrower distribution. TE landscapes were drawn from the RepeatMasker output parsed with the options -l of “parseRM.pl”. We report here the TE landscape at the level of the TE subclass (LINE, LTR, TIR, CRY, MAV, DIRS, PLE, RC and Unknown).

### dnaPipeTE comparative analysis in *Sitophilus* species

To compare the TE content of *S. oryzae* to four related species of *Sitophilus* (*S. granarius, S. zeamais, S. linearis*) we used dnaPipeTE v.1.3 (Goubert et al. 2015). dnaPipeTE allows unbiased estimation and comparison of the total repeat content across different species by assembling and quantifying TE from unassembled reads instead of a linear genome assembly. Reads for *Sitophilus* species were produced as described above. Using our new classified library (3 390 consensus) as TE database in dnaPipeTE, we were further able to identify the phylogenetic depth of the repeat identified in *S. oryzae*.

### RNA sequencing and TE expression analysis

Adapter sequences and low quality reads were filtered out with Trimmomatic (v0.36) (Bolger et al. 2014) and clean reads were aligned to the *S. oryzae* genome with STAR aligner (v2.5.4b, (Dobin et al. 2013)) and featureCounts from subread package (Liao et al. 2014) to obtain gene counts. We also used the STAR aligner to map the clean reads against all TE copies extracted from the genome with the following options: --outFilterMultimapNmax 100 -- winAnchorMultimapNmax 100 --outMultimapperOrder Random --outSAMmultNmax 1. The mapped bam files were used as input to TEtools software (Lerat et al. 2017) to determine TE family expression. Genes and TE family counts were used as input for DESEq2 package (Love et al. 2014) to determine differential TE expression between Ovary vs Gut tissues as well as Ovaries from symbiotic and aposymbiotic weevils. Differentially expressed TEs were defined whenever the adjusted p-value was smaller than 0.05 and Log2 fold change was higher than 1 or smaller than -1. We used the aforementioned STAR alignment parameters to map transcriptomic sequencing reads from midgut of *S. oryzae* (Accession: SRX1034971, and SRX1034972), *D. melanogaster* (Accession: SRX029389, and SRX045361), and *Ae. albopictus* (Accession: SRX1512976, SRX1898481, SRX1898483, SRX1898487, SRX3939061, and SRX3939054) against the TE consensus sequences for each species.

## Abbreviations

AMPs: AntiMicrobial Peptides
CPs: cuticle proteins
HGT: horizontal gene transfer
IMP: Inosine MonoPhosphate
K2P: Kimura 2 Parameters
LINE: long INterspersed Element
LTR: Long Terminal Repeat
MAMPs: Microbial Associated Molecular Patterns
MITEs: miniature inverted repeat elements
ORs: odorant receptors
PRR: Pattern Recognition Receptors
PLE: Penelope-like
RC: rolling circle
SINE: Short INterspersed Element
TIR: terminal inverted repeat
TSS: transcription start sites
TEs: transposable elements
TTSS: Type Three Secretion Systems
UTR: untranslated regions
UMP: Uridine MonoPhosphate

## Data access

This Whole Genome Shotgun project has been deposited at DDBJ/ENA/GenBank under the accession PPTJ00000000. The version described in this paper is version PPTJ02000000. The assembly can be visualised, along with gene models and supporting data, on a dedicated genome browser (https://bipaa.genouest.org/sp/sitophilus_oryzae/). Raw reads from low coverage genome sequencing of *S. zeamais*, *S. granarius* and *S. linearis* have been deposited at NCBI Sequence Read Archive (SRA) under the BioProject accessions PRJNA647530, PRJNA647520 and PRJNA647347 respectively. TE annotation (GFF) and consensus sequences can be found at https://dx.doi.org/10.5281/zenodo.4570415. Bisulfite-seq reads have been deposited at NCBI SRA, under the BioProject accession PRJNA681724.

## Acknowledgments

The authors acknowledge supercomputing resources made available by the Rhône-Alpes Bioinformatics center (PRABI-AMSB, http://www.prabi.fr) to perform the NGS data analyses. The authors would like to thank Stéphanie Robin, Fabrice Legeai and Anthony Bretaudeau at the INRAE BioInformatics Platform for Agro-ecosystems Arthropods (BIPAA) (https://bipaa.genouest.org) for the *S. oryzae* genome integration. The authors would like to thank Fabienne Barbet, Séverine Croze and Nicolas Nazaret from profileXpert for Illumina sequencing of *Sitophilus oryzae* libraries. The authors thank the network REacTION and its coordinators (N. Ponts, G. Le Trionnaire, I. Fudal, M. Jubault), funded by INRAE-SPE, for organizing a Bisulfite-seq workshop that allowed the production of the data presented here. We also thank J.-Y. Rasplus for collecting *Sitophilus linearis* individuals from Niger. The authors would like to thank Cédric Feschotte at Cornell University for the support, insights and the occasional use of bioinformatic resources.

## Authors’ contributions

AH and AL conceived the original sequencing project and were joined by NP, RR, CG, CV-C, AM and CV who participated in the coordination of the project. AVa, CV-M, ED, JM, FM and AVi reared the inbred lines and AVa extracted genomic DNA and RNA that was used for library construction and sequencing. BG and SH produced and sequenced the Nanopore libraries. CG, AVa, MB, NB, CV, AG and ATRV produced and sequenced the low-coverage Illumina libraries. NP, CV-C, ADG and M-FS performed the genome assembly and automated gene prediction. CV-C, MM-H and TG analyzed and wrote the *phylome and horizontal gene transfer* note. PB-P, GF, SC, HC and FC analyzed and wrote *the global analysis of metabolic pathways* note. NP analyzed and wrote the *digestive enzymes* and the *detoxification and insecticide resistance* notes. PC analyzed and wrote the *development* note. CV-C analyzed and wrote the *cuticle protein genes* note. CV-M, CV-C, NP, JM, LB, AB, WZ, FM, AVi and AZ-R analyzed and wrote the *innate immune system* note. NM, CM, ASB and EJ-J analyzed and wrote the *odorant receptors* note. TC, CB, AVa and RR produced the data for the *epigenetic pathways* note. TC, CB, AVa, GR, CV-C, CV and RR, analyzed and wrote the *epigenetic pathways* note. MGF, CG, ED, RR, SB, GF, NM, CV-M and NP produced the figures. CG, RR, JMS, JR, RH and AFAS annotated and analyzed the TE content while MGF analyzed the TE RNAseq data. NP, CV-C, CG, CV, RR, AL and AH wrote the manuscript. All authors read and approved the final manuscript.

## Supplemental Material

Supplemental file S1: Supplemental notes, supplemental figures, and small tables. (PDF)

Supplemental table S2: Transposable elements annotation tables. (XLSX)

Supplemental table S3: STAR and TEtools mapping statistics. (XLSX)

Supplemental table S4: Summary of sequencing libraries produced for S. oryzae. (XLSX)

Supplemental table S5: Large supporting tables and datasets. (XLSX)

