## Supplemental file S1 for "The genome of the cereal pest *Sitophilus oryzae*: a transposable element haven"

[1] Univ Lyon, INSA Lyon, INRAE, BF2I, UMR 203, 69621 Villeurbanne, France.
[2] Institute for Integrative Systems Biology (I2SySBio), Universitat de València and Spanish Research Council (CSIC), València, Spain.
[3] Laboratoire de Biométrie et Biologie Evolutive, UMR5558, Université Lyon 1, Université Lyon, Villeurbanne, France.
[4] Department of Molecular Biology and Genetics, 526 Campus Rd, Cornell University, Ithaca, New York 14853, USA.
[5] KU Leuven, University of Leuven, Department of Human Genetics, Laboratory of Behavioral and Developmental Genetics, B-3000, Leuven, Belgium.
[6] ERABLE European Team, INRIA, Rhône-Alpes, France.
[7] INRAE, Sorbonne Université, CNRS, IRD, UPEC, Université de Paris, Institute of Ecology and Environmental Sciences of Paris, Versailles, France.
[8] Instituto de Ciencias de la Ingeniería, Universidad de O'Higgins, Rancagua, Chile.
[9] Life Sciences, Barcelona Supercomputing Centre (BSC-CNS), Barcelona, Spain.
[10] Mechanisms of Disease. Institute for Research in Biomedicine (IRB), Barcelona, Spain.
[11] Institut Catalan de Recerca i Estudis Avançats (ICREA), Barcelona, Spain.
[12] Laboratório de Bioinformática, Laboratório Nacional de Computação Científica, Petrópolis, Brazil.
[13] Institut de Génomique Fonctionnelle de Lyon (IGFL), Université de Lyon, Ecole Normale Supérieure de Lyon, CNRS UMR 5242, Lyon, France.
[14] Institute for Systems Biology, Seattle, WA, USA.
[15] Foundation for the Promotion of Sanitary and Biomedical Research of Valencian Community (FISABIO), València, Spain.
[16] IGEPP, INRAE, Institut Agro, Université de Rennes, Domaine de la Motte, 35653 Le Rheu, France.

[†] Present address: Institute of Evolutionary Biology (IBE), CSIC-Universitat Pompeu Fabra, Barcelona, Spain.
[‡] Present address: Human Genetics, McGill University, Montreal, QC, Canada.
[§] Present address: LSTM, Laboratoire des Symbioses Tropicales et Méditerranéennes, IRD, CIRAD, INRAE, SupAgro, Univ Montpellier, Montpellier, France.
[£] Present address: Global Health Institute, School of Life Sciences, Ecole Polytechnique Fédérale de Lausanne (EPFL), Lausanne 1015, Switzerland.
[ǁ] Present address: School of BioSciences, The University of Melbourne, Parkville, VIC 3010, Australia.
[#] Present address: Department of Evolutionary Ecology, Institute for Organismic and Molecular Evolution, Johannes Gutenberg University, 55128 Mainz, Germany.

[$] Authors contributed equally to this work.
[*] Authors contributed equally to this work.

### Table of contents


**1. Phylome and horizontal gene transfer** **3**

1.1 Introduction 3

1.2 Methods 3

1.3 Results and discussion 6

**2. Global analysis of metabolic pathways** **12**

2.1 Introduction 12

2.2 Methods 12

2.3 Results and discussion 13

**3. Digestive enzymes** **20**

3.1 Introduction 20

3.2 Methods 21

3.3 Results and discussion 21

**4. Development** **25**

4.1 Introduction 25

4.2 Methods 25

4.3 Results and discussion 26

**5. Cuticle protein genes** **30**

5.1 Introduction 30

5.2 Methods 30

5.3 Results and discussion 31

**6. Innate Immune system** **33**

6.1 Introduction 33

6.2 Methods 34

6.3 Results and discussion 34

**7. Detoxification and Insecticide resistance** **46**

7.1 Introduction 46

7.2 Methods 46

7.3 Results and discussion 47

**8. Odorant receptors** **49**

8.1 Introduction 49

8.2 Methods 50

8.3 Results and discussion 50

**9. Epigenetic pathways** **53**

9.1 Introduction 53

9.2 Methods 54

9.3 Results and Discussion 56

**10. Supplemental figures** **62**

**References** **68**

### 1. Phylome and horizontal gene transfer

Carlos VARGAS-CHAVEZ, Marina MARCET-HOUBEN and Toni GABALDON

#### 1.1 Introduction

Even between closely related organisms there is an enormous amount of variability in the sizes of their gene families. These changes are relevant given that the gain or loss of even a single gene can be involved in the adaptive divergence between species. To identify these changes we generated the full phylome for *Sitophilus oryzae*. A phylome represents the complete collection of all gene phylogenies in a genome. It can be used to uncover the evolutionary relationships between all the proteins in a genome and the proteomes of other species of interest. Additionally, to identify rapidly evolving gene families along the *S. oryzae* lineage we used CAFE [1]. With the gene family data and an ultrametric phylogeny, CAFE can be used to estimate gene gain and loss rates taking in consideration the amount of assembly and annotation error in the input data.

#### 1.2 Methods

##### Phylome reconstruction

The phylome of *S. oryzae*, meaning the collection of phylogenetic trees for each gene in its genome, was reconstructed using an automated pipeline that mimics the steps one would take to build a phylogenetic tree. First a database of 17 species was built that included *S. oryzae* and 16 other arthropods (Table S1.1). Then a blastp search was performed against this database starting from each of the proteins included in the genome. Blast results were filtered using an e-value threshold of 1e-05 and an overlap threshold of 50%. The number of hits was limited by the 150 best hits for each protein. Then the multiple sequence alignment was performed for each set of homologous sequences. Three different programs were used to build the alignments (Muscle v3.8.1551 [2], mafft v7.407 [3] and kalign v2.04 [4] and the alignments were performed in forward and in reverse resulting in six different alignments. From these groups of alignments, a consensus alignment was obtained using M-coffee from the T-coffee package v12.0 [5]. Alignments were then trimmed using trimAl v1.4.rev15 (consistency-score cut-off 0.1667, gap-score cut-off 0.9) [6]. IQTREE v1.6.9 [7] was then used to reconstruct a maximum likelihood phylogenetic tree. Model selection was limited to 5 models (DCmut, JTTDCMut, LG, WAG, VT) with freerate categories set to vary between 4 and 10. The best model according to the BIC criterion was used. 1 000 rapid bootstraps were calculated. All trees and alignments were stored in phylomedb [8] with phylomeID 43 (http://phylomedb.org).

**Table S1.1.** List of the species used for phylome reconstruction.

| NCBI Tax ID | Species | Source |
|---|---|---|
| 7029 | *Acyrthosiphon pisum* | NCBI (GCF_005508785.1) |
| 7165 | *Anopheles gambiae* | QFO8 |
| 7460 | *Apis mellifera* | NCBI |
| 7091 | *Bombyx mori* | Ensembl Metazoa release 25 |
| 104421 | *Camponotus floridanus* | NCBI |
| 6669 | *Daphnia pulex* | Ensembl Metazoa release 25 |
| 77166 | *Dendroctonus ponderosae* | Uniprot |
| 121845 | *Diaphorina citri* | NCBI |
| 7227 | *Drosophila melanogaster* | QFO8 |
| 37546 | *Glossina morsitans* | VectorBASE |
| 7130 | *Manduca sexta* | i5k.nal.usda.gov |
| 7425 | *Nasonia vitripennis* | Ensembl Metazoa release 25 |
| 121224 | *Pediculus humanus* | Ensembl Metazoa release 25 |
| 51655 | *Plutella xylostella* | NCBI (GCF_000330985.1) |
| 7048 | *Sitophilus oryzae* | NCBI (GCF_002938485.1) |
| 32264 | *Tetranychus urticae* | - |
| 7070 | *Tribolium castaneum* | Ensembl Metazoa release 25 |

A species tree was reconstructed using a concatenation method, and 184 single copy protein families were concatenated into a single multiple sequence alignment. IQTREE was then used to reconstruct the species tree [7]. The alignment contained 102 539 positions. The model selected for tree reconstruction was LG+F+R7. Additionally, duptree [9] was used to reconstruct a second species tree using a super tree method. All trees built during the phylome reconstruction process were used to reconstruct the species tree.

Proteins likely to be transposable elements (TEs) were detected by running a HMMER [10] search with 99 pfam domains that have been related to transposases. 1 411 proteins were identified as having one or more of these Pfam domains and trees containing those proteins were removed from the set of trees to analyze (3 031 trees were removed). Gene trees were rooted using a species to age dictionary in which a preferred list of outgroups was put. Then duplication nodes were inferred using the species overlap algorithm in which for each node in the tree species on both sides of the node are compared, and if there is an overlap, the node is considered to be a duplication node, otherwise it is considered a speciation node. Duplication nodes are then assigned to the species tree considering that the duplication happened at the common ancestor of all the species found in the node. The total number of duplications is then divided by the number of trees that have this node in their tree. Only duplications with a rapid bootstrap value of 90 are considered. GO term enrichment for each group of duplications was calculated using a python adaptation of FatiGO [11] (corrected p-value $< 0.01$). Species specific expanded families were calculated by clustering sets of species-specific paralogs using a UPGMA clustering approach.

**Estimating gene gain and loss rates**

EGGNOG-MAPPER was used to identify orthologs among the selected species using the diamond mode and the arthropoda (artNOG) dataset. All 1 345 genes with one-to-one orthologues in all 11 coleopterans included in the analysis (*Aethina tumida*, *Anoplophora glabripennis*, *Agrilus planipennis*, *Dendroctonus ponderosae*, *Diabrotica virgifera*, *Leptinotarsa decemlineata*, *Nicrophorus vespilloides*, *Onthophagus* *taurus*, *Photinus pyralis*, *S. oryzae* and *Tribolium castaneum*) were selected and their alignments concatenated to generate a species tree using ETE-BUILD [12] following the phylomedb4 gene tree workflow and the sptree_raxml_85 species tree workflow. The tree was converted to an ultrametric topology using the ape package [13] from R [14]. This ultrametric tree was used along with the gene families' data to estimate gene losses and gains using CAFE [1] after having estimated the error rate and corrected for it. Using the gene loss and gain rate the ancestral state gene counts are inferred and a p-value is calculated to evaluate the relevance of the gene family changes along each branch.

#### 1.3 Results and discussion

For the analysis of the phylome, a total of 13 519 trees were reconstructed, and those that contained putative transposons were discarded. This reduced the number of trees to analyse to 10 488 (see methods). We detected a total of 48 expanded gene families that contained in total 437 proteins specifically in *S. oryzae*. The largest of such expansions contained 30 proteins predicted to have a conserved THAP domain (PF05485) and associated to the GO term nucleic acid binding (GO:0003676). The next two largest groups, containing 28 and 21 paralogs respectively both contained proteins with zinc finger domains which likely indicate that the three groups are formed by transcription factors.

Using the phylome, we also explored which GO terms were enriched in duplications at other nodes in the tree leading to *S. oryzae*. GO terms involved in perception of smell (GO:0007608) and olfactory receptors and odorant binding were enriched in duplications in nodes belonging to the species of the Cucujiformia infraorder. This indicates an on-going trend to duplicate genes involved in olfaction in this group of species. This is particularly visible for the Odorant Receptor (OR) family, in which a large number of duplications specific to the Curculionidae family notably occurred in one particular lineage of the OR phylogeny (see Supplemental Note 8). Terms related to taste (GO:0008527, GO:0050909 and GO:0050912) were also enriched in duplications in the node leading to the Cucujiformia infraorder but they were not enriched in *S. oryzae* species specific duplications. In the duplications in the nodes leading to the Curculionidae family and specific for *S. oryzae* we observed an enrichment for cellulase activity (GO:0008810), which might be involved with an ancestral horizontal gene transfer event in the Curculionidae ancestor and further species specific expansions (see below and Supplemental Note 3 on Digestive enzymes).

Using CAFE to analyse coleopterans, we found 109 rapidly evolving gene families along the *S. oryzae* lineage, all of which are rapid expansions (Table S1.2). Among all species included in our analysis, *S. oryzae* had the third highest average expansion rate (0.088 genes per million years) after *D. virgifera* and *P. pyralis* (Figure S1.1). More than half of the expanded families with assigned putative function (43/84) are likely involved with TEs (Table S1.3, TE related proteins highlighted). We also observed a major expansion in a family of putative transcription factors with zinc finger domains, similar to what we had observed using the phylome. Lineage specific expansions of zinc-finger proteins have been described in other organisms such as *D. melanogaster* [15] and it has been suggested that these expansions might allow the evolution of novel functions [16]. Additionally, we observed a moderate expansion in a juvenile hormone-inducible protein family which could lead to an accelerated development in symbiotic insects [17]. There was also an expansion in a Cytochrome P450 family (+10) and two galactosyltransferase-like families (+7 and +4), which might be involved in insecticide resistance [18]. Finally, we observed a moderate expansion of immune effectors (+4) which are likely antimicrobial peptides that confer increased resistance against fungi or other pathogens (See Supplemental Note 6 on the Innate Immune system [19]).

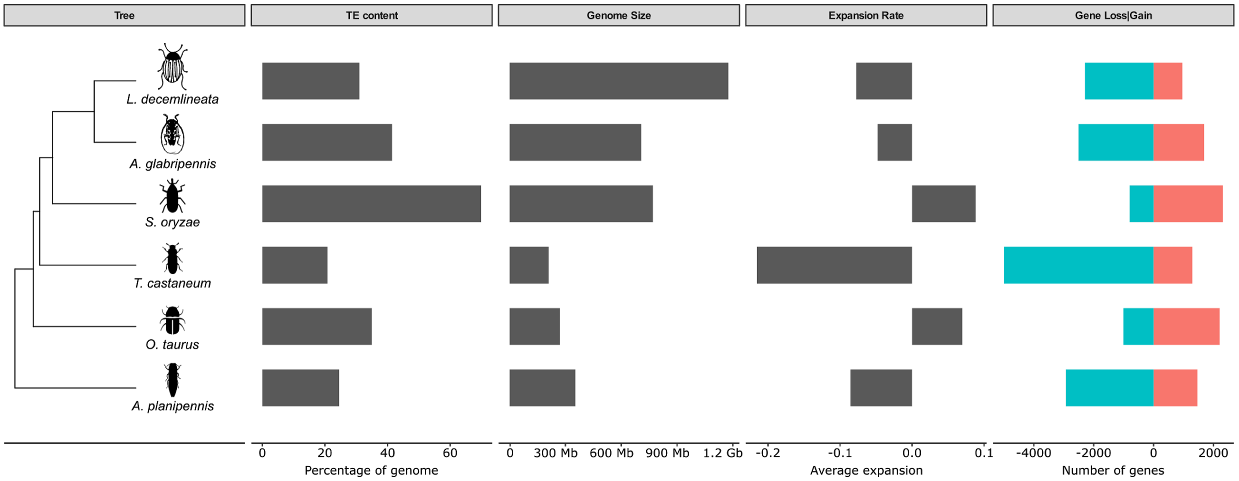


**Figure S1.1. Gene gain and loss in six coleopteran species.** The phylogeny of the species analyzed is shown along with the fraction of the genome spanned by TEs and each species' genome size. Average expansion refers to the mean number of genes gained or lost per family where negative values indicate lost genes and positive values gained genes.

**Table S1.2. Summary of gene gain and loss for all coleopteran species analyzed in this study.**

| Species | Expanded families | Genes gained | Expansion rate | Contracted families | Genes lost | Contraction rate | No change | Average Expansion rate |
|---|---|---|---|---|---|---|---|---|
| ***S. oryzae*** | 1022 (109) | 2315 | 2.27 | 790 (0) | 799 | 1.01 | 15348 | 0.088345 |
| ***D. ponderosae*** | 1007 (15) | 1410 | 1.4 | 1313 (13) | 1401 | 1.07 | 14840 | 0.00052448 |
| ***L. decemlineata*** | 649 (59) | 962 | 1.48 | 2169 (53) | 2293 | 1.06 | 14342 | -0.0775641 |
| ***D. virgifera*** | 1689 (399) | 4012 | 2.38 | 1174 (12) | 1210 | 1.03 | 14297 | 0.163287 |
| ***A. glabripennis*** | 784 (96) | 1689 | 2.15 | 2489 (3) | 2509 | 1.01 | 13887 | -0.0477855 |
| ***A. tumida*** | 1472 (16) | 2016 | 1.37 | 4678 (5) | 4749 | 1.02 | 11010 | -0.159266 |
| ***T. castaneum*** | 595 (23) | 1295 | 2.18 | 4949 (1) | 4994 | 1.01 | 11616 | -0.215559 |
| ***O. taurus*** | 1118 (87) | 2207 | 1.97 | 957 (9) | 1009 | 1.05 | 15085 | 0.0698135 |
| ***N. vespilloides*** | 509 (40) | 1116 | 2.19 | 1314 (15) | 1419 | 1.08 | 15337 | -0.0176573 |
| ***P. pyralis*** | 2909 (167) | 6594 | 2.27 | 1076 (2) | 1117 | 1.04 | 13175 | 0.319172 |
| ***A. planipennis*** | 954 (18) | 1464 | 1.53 | 2632 (22) | 2931 | 1.11 | 13574 | -0.0854895 |

**Table S1.3. List of the expanded gene families in *S. oryzae*.** In red, gene families with functions linked to transposable elements.

| Genes gained | Eggnog annotation |
|---|---|
| 77 | Pao retrotransposon peptidase |
| 49 | Inherit from KOG: Retrotransposon protein |
| 43 | Inherit from KOG: Zinc finger protein |
| 40 | Endonuclease/Exonuclease/phosphatase family |
| 36 | Endonuclease/Exonuclease/phosphatase family |
| 30 | reverse transcriptase |
| 28 | - |
| 24 | Reverse transcriptase (RNA-dependent DNA polymerase) |
| 23 | to Tigger transposable element-derived protein 6 |
| 21 | Pfam:DDE |
| 19 | Inherit from euNOG: reverse transcriptase |
| 19 | Inherit from opiNOG: Pao retrotransposon peptidase |
| 19 | Protein of unknown function (DUF3609) |
| 18 | Endonuclease/Exonuclease/phosphatase family |
| 18 | Inherit from meNOG: to H28G03.4 Hydra magnipapillata |
| 17 | Inherit from meNOG: protein Hydra magnipapillata |
| 12 | - |
| 11 | Endonuclease/Exonuclease/phosphatase family |
| 11 | Inherit from meNOG: protein Hydra magnipapillata |
| 11 | Inherit from meNOG: protein Hydra magnipapillata |
| 11 | Plant transposon protein |
| 11 | - |
| 10 | Inherit from biNOG: cytochrome P450 |
| 10 | Inherit from KOG: Retrotransposon protein |
| 10 | Pfam:DDE |
| 10 | Reverse transcriptase (RNA-dependent DNA polymerase) |
| 9 | Inherit from KOG: transposon protein |
| 9 | - |
| 8 | Inherit from biNOG: general transcription factor II-I repeat domain-containing protein |
| 8 | Inherit from biNOG: Reverse transcriptase (RNA-dependent DNA polymerase) |
| 8 | Inherit from opiNOG: to reverse transcriptase |
| 8 | Juvenile hormone-inducible protein |
| 8 | Pfam:DDE |
| 8 | Plant transposon protein |
| 8 | to Y54G2A.42 |
| 8 | - |
| 7 | Galactosyltransferase |
| 7 | Inherit from artNOG: YqaJ-like viral recombinase domain |
| 7 | Inherit from euNOG: reverse transcriptase |
| 7 | Inherit from KOG: transposon protein |
| 7 | Inherit from meNOG: protein Hydra magnipapillata |
| 7 | MADF |
| 7 | Plant transposon protein |
| 7 | Reverse transcriptase (RNA-dependent DNA polymerase) |
| 7 | ZnF_BED |
| 7 | - |
| 7 | - |
| 7 | - |
| 7 | - |
| 6 | Endonuclease/Exonuclease/phosphatase family |
| 6 | Inherit from artNOG: Endonuclease-reverse transcriptase HmRTE-e01 |
| 6 | Inherit from biNOG: piggyBac transposable element derived |
| 6 | Inherit from opiNOG: protein Hydra magnipapillata |
| 6 | Inherit from opiNOG: to reverse transcriptase |
| 6 | Pfam:DUF889 |
| 6 | Phage integrase family |
| 6 | Phage integrase family |
| 6 | Plant transposon protein |
| 6 | pol-like protein |
| 6 | Protein of unknown function (DUF3421) |
| 6 | reverse transcriptase |
| 6 | Reverse transcriptase (RNA-dependent DNA polymerase) |
| 6 | - |
| 6 | - |
| 6 | - |
| 6 | - |
| 6 | - |
| 6 | - |
| 5 | Alcohol dehydrogenase transcription factor Myb/SANT-like |
| 5 | cuticular protein |
| 5 | Inherit from biNOG: harbinger transposase derived 1 |
| 5 | Inherit from biNOG: piggyBac transposable element derived |
| 5 | Inherit from biNOG: SCAN domain containing 3 |
| 5 | Inherit from COG: Retrotransposon protein |
| 5 | jerky protein homolog-like |
| 5 | Matrixin |

| | |
|---|---|
| 5 | Pao retrotransposon peptidase |
| 5 | Reverse transcriptase (RNA-dependent DNA polymerase) |
| 5 | to Y54G2A.42 |
| 5 | - |
| 5 | - |
| 5 | - |
| 5 | - |
| 5 | - |
| 4 | Aldo/keto reductase family |
| 4 | ec 3.2.1.20 |
| 4 | galactosyltransferase activity |
| 4 | Inherit from biNOG: piggyBac transposable element derived |
| 4 | Inherit from biNOG: Thaumatin family |
| 4 | Inherit from meNOG: protein F54H12.3, partial Hydra magnipapillata |
| 4 | jerky protein homolog-like |
| 4 | Pfam:DDE |
| 4 | PHD |
| 4 | Plant transposon protein |
| 4 | pol-like protein |
| 4 | Protein of unknown function (DUF3609) |
| 4 | - |
| 4 | - |
| 4 | - |
| 3 | Acyltransferase family |
| 3 | Inherit from biNOG: piggyBac transposable element derived |
| 3 | Inherit from opiNOG: to reverse transcriptase |
| 3 | MADF |
| 3 | Odorant receptor |
| 3 | Odorant receptor |
| 3 | to conserved |
| 3 | to conserved |
| 3 | - |
| 3 | - |

**Horizontally transferred genes**

Horizontally transferred genes result from the movement of genetic material between organisms. Using a combination of tools including DARKHORSE [20] and HGT-FINDER [21] we identified hundreds of candidates; however, after manually curating the list of candidates, most of them were discarded given that the majority are likely retroviral mediated transfers. The putative donors of the remaining 31 candidates were identified (Table S1.4), and most of such candidates have a digestive-related function with 24 being putative plant cell wall degrading enzymes (see Supplemental Note 3 on Digestive enzymes). We also identified homologs for these genes in other members of the Cucujiformia suborder, suggesting that the transfers took place before the divergence of the clade. Regardless of the ideal conditions for gene transfers, we did not identify any HGT event potentially deriving from either *Wolbachia* or *S. pierantonius, S. oryzae*´s two current endosymbionts. Given the recent acquisition of *S. pierantonius* [22] perhaps there has not been enough time for a HGT event to have occurred. However, the lack of HGT events from *Wolbachia* is more surprising given that *Wolbachia*-like sequences have been found in numerous arthropods [23] including symbiotic models like tsetse flies [24] and aphids [25]. Interestingly, we also observed several transfers from an Enterobacteriales donor. This group includes, among other endosymbionts, Candidatus *Nardonella*, the ancestral symbiont of *S. oryzae*.

**Table S1.4. List of horizontally transferred candidate genes in *S. oryzae*.** 18 candidates had been previously described in Pauchet *et al.* [26]. Two candidates marked with an asterisk represent highly similar but not identical sequences to those previously described.

| ID | Function | Donor | Found in | ID from Pauchet ***et al.*** (26) |
|---|---|---|---|---|
| XP_030746517.1 | cellulose 1,4-beta-cellobiosidase | Streptomycetaceae | Phytophaga | ADU33251.1 |
| XP_030746518.1 | cellulose 1,4-beta-cellobiosidase | Streptomycetaceae | Phytophaga | ADU33252.1 |
| XP_030757664.1 | cyclase family protein | Bacteria | Polyphaga | - |
| XP_030763223.1 | endoglucanase-like | Fungi | Phytophaga | - |
| XP_030763224.1 | endoglucanase-like | Fungi | Phytophaga | - |
| XP_030747083.1 | endoglucanase-like | Fungi | Phytophaga | ADU33246.1 |
| XP_030751361.1 | endoglucanase-like | Fungi | Phytophaga | ADU33247.1 |
| XP_030751155.1 | endoglucanase-like | Fungi | Phytophaga | ADU33248.1 |
| XP_030751166.1 | endoglucanase-like | Fungi | Phytophaga | ADU33249.1 |
| XP_030763222.1 | endoglucanase-like | Fungi | Phytophaga | ADU33250.1 |
| XP_030763233.1 | endoglucanase-like | Fungi | Phytophaga | ADU33250.1* |
| XP_030753212.1 | glycoside hydrolase family 32 protein | Enterobacterales | Polyphaga | - |
| XP_030753449.1 | glycoside hydrolase family 32 protein | Enterobacterales | Polyphaga | - |
| XP_030757274.1 | methylated-DNA--[protein]-cysteine S-methyltransferase | Bacteria | Polyphaga | - |
| XP_030748053.1 | pectinesterase | Enterobacterales | Curculionidae | ADU33259.1 |
| XP_030746612.1 | pectinesterase | Enterobacterales | Curculionidae | ADU33260.1 |
| XP_030746614.1 | pectinesterase | Enterobacterales | Curculionidae | ADU33261.1 |
| XP_030762872.1 | pectinesterase | Enterobacterales | Curculionidae | ADU33262.1 |
| XP_030762871.1 | pectinesterase | Enterobacterales | Curculionidae | ADU33263.1 |
| XP_030751830.1 | p-loop containing nucleoside triphosphate hydrolase | Fungi | Polyphaga | - |
| XP_030761554.1 | p-loop containing nucleoside triphosphate hydrolase | Fungi | Polyphaga | - |
| XP_030761555.1 | p-loop containing nucleoside triphosphate hydrolase | Fungi | Polyphaga | - |
| XP_030767685.1 | polygalacturonase-like | leotiomyceta | Phytophaga | ADU33253.1 |
| XP_030767687.1 | polygalacturonase-like | leotiomyceta | Phytophaga | ADU33254.1 |
| XP_030745830.1 | polygalacturonase-like | saccharomyceta | Phytophaga | ADU33255.1 |
| XP_030745832.1 | polygalacturonase-like | saccharomyceta | Phytophaga | ADU33255.1* |
| XP_030745511.1 | polygalacturonase-like | leotiomyceta | Polyphaga | ADU33256.1 |
| XP_030745833.1 | polygalacturonase-like | saccharomyceta | Phytophaga | ADU33257.1 |
| XP_030757439.1 | polygalacturonase-like | leotiomyceta | Phytophaga | ADU33258.1 |
| XP_030764144.1 | protein phosphatase PP2A regulatory subunit A-like | saccharomyceta | Polyphaga | - |
| XP_030761743.1 | uracil-DNA glycosylase | Bacteria | Polyphaga | - |

### 2. Global analysis of metabolic pathways

Patrice BAA-PUYOULET, Gérard FEBVAY, Stefano COLELLA, Hubert CHARLES and Federica CALEVRO

#### 2.1 Introduction

The automated functional annotations of the 20 947 predicted proteins from *S. oryzae* genome were specified using the CycADS pipeline and gathered in the dedicated database SitorCyc (see Methods). Automatic and manual annotations have been carried out to explore the metabolic network of *S. oryzae* and its endosymbiont *S. pierantonius* with a specific focus on central metabolism (*i.e.* amino acids, carbohydrates, fatty acids and nucleotides), vitamins and cofactors in order to determine the compounds that derive from the diet, and those that need to be shuttled between the associated partners. Our global genomic analysis of the *S. oryzae*/*S. pierantonius* association reveals that the specific losses of enzymes in *S. oryzae* are limited, whereas *S. pierantonius* genome has suffered much more degradations. Furthermore, this analysis shows host/symbiont collaboration, particularly for some essential vitamins biosynthetic pathways that require genome complementarities between these two organisms, and/or assimilation and exchange of dietary compounds.

#### 2.2 Methods

Automated functional annotations of predicted proteins from *S. oryzae* genome (NCBI Annotation Release 100) were specified using the CycADS pipeline [27] allowing the reconstruction of metabolic networks using the PATHWAYTOOLS software [28]. The SitorCyc database was generated and added to the ArthropodaCyc collection (http://arthropodacyc.cycadsys.org/) [29]. This allowed the comparisons with the 40 other arthropod species of the collection, and more precisely with the six other coleopteran species belonging to the Cucujiformia infraorder including the mountain pine beetle *Dendroctonus* *ponderosae* (Curculionidae), the Colorado potato beetle *Leptinotarsa decemlineata* (Chrysomelidae), the Western corn rootworm *Diabrotica virgifera* (Chrysomelidae), the Asian long-horned beetle *Anoplophora glabripennis* (Cerambycidae), the small hive beetle *Aethina tumida* (Nitidulidae) and the red flour beetle *Tribolium castaneum* (Tenebrionidae).

Starting from this metabolic network reconstruction, we focused on the major metabolic pathways, and specifically the ones involved in the biosynthesis of amino acids, energy metabolism, amino sugar and nucleotide sugar metabolism, as well as metabolism of cofactors and vitamins. The metabolic network of the endosymbiont *S. pierantonius* was reconstructed using the same method and we focused our analyses on the pathway interconnections between the associated partners.

#### 2.3 Results and discussion

The global metabolism analysis detected the presence of 1 387 different Enzyme Commission numbers (ECs) among the annotated proteins in *S. oryzae,* which is fully consistent with other Cucujiformia beetle genomes (whose EC repertoires range from the 1 308 of *D. ponderosae* to the 1 388 of *D. virgifera* (Table S2.1)).

**Table S2.1. Summary of the ArthropodaCyc annotation of metabolic genes in the Cucujiformia taxon.**

| Species | *Sitophilus oryzae* | *Dendroctonus ponderosae* | *Leptinotarsa decemlineata* | *Diabrotica virgifera* | *Anoplophora glabripennis* | *Aethina tumida* | *Tribolium castaneum* |
|---|---|---|---|---|---|---|---|
| **Order, Family** | Coleoptera, Curculionidae | Coleoptera, Curculionidae | Coleoptera, Chrysomelidae | Coleoptera, Chrysomelidae | Coleoptera, Cerambycidae | Coleoptera, Nitidulidae | Coleoptera, Tenebrionidae |
| **Gene set ID** | NCBI Sitophilus oryzae Annotation Release 100 | NCBI GCA_000355655 1.29 DendPond_male_1.0 | NCBI GCF_00050032 5.1 Ldec_2.0 | NCBI GCF_00301383 5.1 Dvir_v2.0 | NCBI GCF_00039028 5.2 Agla_2.0 | NCBI GCF_00193 7115.1 Atum_1.0 | NCBI GCF_0000023 35.3 Tcas5.2 |
| **CycADS Database ID** | SitorCyc | DenpoCyc | LepdeCyc | DiaviCyc | AnoglCyc | AettuCyc | TricaCyc |
| **Polypeptides** | 20,947 | 13,467 | 17,595 | 26,275 | 19,013 | 16,585 | 19,883 |
| **Pathways** | 308 | 297 | 308 | 303 | 309 | 310 | 300 |
| **Enzymatic reactions** | 3,334 | 3,156 | 3,288 | 3,300 | 3,315 | 3,282 | 3,289 |
| **Enzymes** | 5,189 | 3,461 | 4,473 | 5,372 | 5,003 | 4,392 | 4,984 |
| **Compounds** | 2,257 | 2,157 | 2,219 | 2,248 | 2,247 | 2,227 | 2,258 |
| **EC[1] present in the genome** | 1,387 | 1,308 | 1,352 | 1,388 | 1,370 | 1,378 | 1,382 |
| **EC unique to this genome[2]** | 40 | 13 | 17 | 29 | 13 | 16 | 18 |
| **EC missing only in this genome[2]** | 14 | 43 | 23 | 10 | 5 | 4 | 7 |

[1] "EC" refers to the number of proteins, as represented by their unique numerical designations within the Enzyme Commission (EC) classification system for enzymes and their catalytic reactions.
[2] in comparison with the other seven *Cucujiformia* genomes.

We then looked at the metabolic networks of the *S. oryzae*/*S. pierantonius* association. For the synthesis of organic nitrogen compounds like amino acids, insects, as all animals depend entirely on organic nitrogen supply. Comparatively to *Sodalis praecaptivus* (a closely related free-living bacterium [30], *S. pierantonius* has lost the nitrate and nitrite reductase activities, and is unable to assimilate nitrate and to reduce it into ammonia. Consequently, the symbiotic partners depend entirely on an external organic nitrogen supply from the diet, i.e., proteins of the wheat grain. *In silico* prediction of the amino acid biosynthetic pathways reveals that the *S. oryzae* genome possesses all the enzyme-encoding genes required for the biosynthesis of all non-essential amino acids. Like in other metazoan organisms, essential amino acid requirements should be satisfied by the diet and/or by the bacterial symbionts. Hence, *S. pierantonius* only retained the capability of supplying four essential amino acids (Thr, Phe, Lys and Arg), suggesting that the other six (Met, Val, Leu, Ile, Trp and His) must be obtained from the diet. In return, *S. pierantonius* is dependent on the host for the supply of the non-essential proline (Figure S2.1). Eventually, among the three distinct metabolic routes for the production of Ala in Bacteria, two appear to be absent in *S. pierantonius*. The route from cysteine desulfurase is complete, but the role of this pathway in generating the unique cellular supply of Ala has not been fully demonstrated. This analysis suggests that Ala biosynthesis could be limited and the endosymbiont may therefore be dependent on its host for the supply of this amino acid.

Starch is the most abundant component of cereal seeds. Indeed, the genome of *S. oryzae* contains highly active midgut α-amylases [31] (see below). Glucose and/or glucose-1-phosphate provided by starch digestion can then be used in the glycolysis, citrate cycle, and pentose phosphate pathways, for which all involved enzymes have been annotated in the *S. oryzae* genome. Glucose can be internalized in the bacteria thanks to a specific sugar transporting PhosphoTransferase System (PTS) [EC 2.7.1.199] that has been retained in the *S. pierantonius* genome. Glycolysis, pentose phosphate, and fatty acid biosynthesis pathways are complete in the endosymbiont, while the citric acid cycle is interrupted by the pseudogenization of the gene encoding aconitate hydratase [EC 4.2.1.3], rendering *S. pierantonius* dependent on its host supply for isocitrate (Figure S2.1).

The genomic annotation shows that in *S. oryzae*, as in other metazoans, salvage pathways should allow the incorporation of purines derived from the degradation of food directly into nucleotides. Additionally, *S. oryzae* has retained the complete pathway for the *de novo* purine nucleotide biosynthesis enabling inosine monophosphate (IMP) biosynthesis. Similarly, *S. oryzae* has retained the salvage and *de novo* pathways for pyrimidine nucleotide synthesis leading to either direct incorporation into nucleotides or production of uridine monophosphate (UMP). In *S. pierantonius*, the salvage pathways for nucleotide synthesis appear to be conserved; however, both *de novo* biosynthetic routes are interrupted due to pseudogenization. Therefore, the endosymbiont entirely depends on its host for nucleotide biosynthesis through precursor supply (IMP, and dihydroorotate or UMP) (Figure S2.1).

*In silico* analyses on *S. pierantonius* metabolic network suggests that biosynthetic pathways of three vitamins, B6, B1, and H (PLP-pyridoxine, thiamine, and biotin, respectively) are disrupted. The dietary requirements for PLP and thiamine are consistent with previous nutritional studies [32]. Conversely, we show here that biotin cannot be provided by the endosymbiont, contrary to what was previously suggested by metabolic complementation experiments [32]. While *S. pierantonius* has kept full pathways for pantothenate (vitamin B5), riboflavin (vitamin B2), and folate (vitamin B9) biosynthesis, these pathways all require the provision of precursors from the insect of its diet (*e.g.* valine and 4-Hydroxybenzoate from the diet and IMP and Coproporphyrinogen III from *S. oryzae*, respectively, Figure S2.1). Starting from these vitamins, *S. oryzae* is then able to perform the final reactions to synthesize active cofactors. Hence, for these three vitamins, host and bacterial pathways are highly interconnected and interdependent.

Like other insects, *S. oryzae* is unable to synthesize *de novo* nicotinamide adenine dinucleotide (NAD) and must obtain it from an external source, while *S. pierantonius* retains the NAD synthesis pathway from aspartate, and is thus able to provide this coenzyme to the weevil. On the other hand, the salvage pathway that rescues NAD seems to have been lost in the endosymbiont and potentially functional in *S. oryzae.* This suggests that nicotinamide is a dead-end product in *S. pierantonius* and that it must be shuttled for salvage into the weevil bacteriocyte. This finding is in agreement with previous studies demonstrating that mitochondrial enzymatic activities were higher in symbiotic than in aposymbiotic insects and that supplementation with pantothenate and riboflavin results in higher oxidative phosphorylation activity [33,34]. Indeed, increase of mitochondrial activity allows symbiotic, but not aposymbiotic, weevils to fly, which directly impacts insect behavior and dissemination [35].

The biosynthetic pathway for the production of lipoic acid, a fatty acid that can be used as a source of cyl groups in several enzyme systems, and the lipoate salvage pathway are possible both in *S. oryzae* and in *S. pierantonius*.

For the ubiquinone (or coenzyme Q) biosynthetic pathway, the final steps could be performed both by the weevil and the endosymbiont. The association seems therefore to only need a source of para-hydroxybenzoate (PHBA) to initiate the ubiquinone synthesis. This phenolic compound is widely distributed in plants [36].

Concerning tetrapyrroles (chemical compounds with four pyrrole rings in either a linear or a cyclic shape), the common precursor of all the family is 5-aminolevulinate and its biosynthetic pathways are retained both by *S. oryzae* and *S. pierantonius*. Additionally, the insect is able to synthesize heme, and thereafter cytochrome c. As the endosymbiont is unable to synthesize these important co-factor/co-enzyme, they must be provided by the host. *S. pierantonius* is capable of synthesizing siroheme, a co-factor of assimilatory sulfite reductase (NADPH) [EC 1.8.1.2] playing a major role in the sulfur assimilation pathway.

Lastly, *S. oryzae* has retained the vitamin A (retinol) biosynthetic pathway starting from β-carotene, a compound that can be found in the diet [37]. It is worth noting that the retinol biosynthetic pathway is absent in *S. pierantonius*.

Taken together, our metabolic analysis attests that *S. pierantonius* became highly dependent on its host at different levels and in a very short coevolutionary period. This illustrates the high genomic plasticity of the genus *Sodalis,* which is known to be associated with a broad spectrum of insect species [38]. Importantly, with the exception of the pathways absent in the majority of animal clades, including the ones involved in the biosynthesis of essential amino acids and vitamins, the host genome does not seem to have lost genes encoding basic metabolic functions. This could explain why the symbiont replacement occurring within the Dryophtorinae subfamily and leading to the diversification of the *Sitophilus* group was possible [22]. These *in silico* analyses provide evidence of the nesting of the two partners' metabolic networks at the genetic level. They also show that the *S. pierantonius* enzyme repertoire could metabolically boost its weevil host, then enabling it to rapidly adapt to human cereal crops, whether stored or in the field.

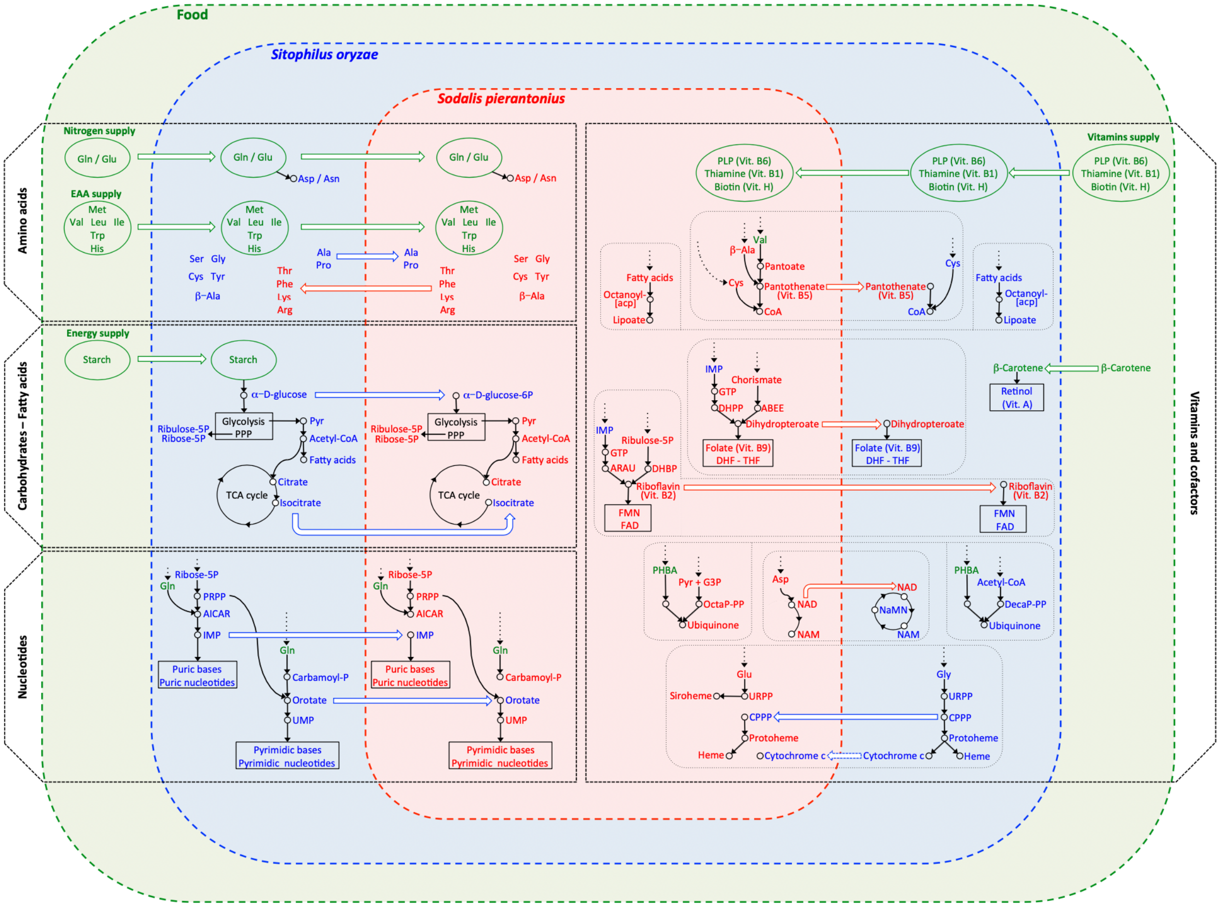


**Figure S2.1. Food supply and metabolic pathway interconnections between the insect *S. oryzae* and its bacterial symbiont, *S. pierantonius*.** The illustration shows the integrated metabolic pathways for the biosynthesis of amino acids, carbohydrates / fatty acids, nucleotides, and vitamins / cofactors between the weevil and its bacterial symbiont, relatively to the food supply. The metabolic compounds acquired through the diet are labelled in green, those produced via the holobiont endogenous metabolism in blue if they are synthesized by the insect,or in red if they are synthesized by the symbiont. The exchanges take the color of the origin of a specific compound (e.g. green for compounds coming from the diet). Compounds coming from diet are indicated only if the insect or its bacterial symbiont cannot synthesize them. Abbreviations: ABEE = 4-Aminobenzoate; AICAR = 5-Aminoimidazole-4-carboxamide ribotide;

ARAU = 5-Amino-6-(1-D-ribitylamino)uracil; CPPP = Coproporphyrinogen III; DecaP-PP = all-trans-Decaprenyl diphosphate; DHBP = L-3,4-Dihydroxybutan-2-one 4-phosphate; DHPP = 7,8-Dihydropterin pyrophosphate; EAA = Essential amino acids; FAD = Flavin adenine dinucleotide; FMN = Flavin mononucleotide; G3P = Glyceraldehyde 3-phosphate; GTP = Guanosine 5'-triphosphate; IMP = Inosine 5'-monophosphate; NAD = Nicotinamide adenine dinucleotide; NAM = Nicotinamide; NaMN = Nicotinic acid mononucleotide; OctaP-PP = all-trans-Octaprenyl diphosphate; PHBA = 4-Hydroxybenzoate; PLP = Pyridoxal 5'-phosphate; PPP = Pentose phosphate pathway; PRPP = 5-Phosphoribosyl 1-pyrophosphate; Pyr = Pyruvate; UMP = Uridine 5'-monophosphate; URPP = Uroporphyrinogen III.

### 3. Digestive enzymes

Nicolas PARISOT

#### 3.1 Introduction

The metabolic reconstruction of the *S. oryzae*/*S. pierantonius* association showed that *S. oryzae* needs to efficiently break down seed proteins to free amino acids in order to survive and develop on its strict diet (the wheat grain). Peptidase families are classified in nine groups according to their active site or their dependence from metal ions: Aspartic (A), Cysteine (C), Glutamic (G), Metallo (M), Asparagine (N), Serine (S), Threonine (T), Mixed (P), and Unknown (U) [39]. For Cucujiformia beetles, which include Curculionidae, cysteine peptidases are the major digestive proteinases [40,41]. More specifically, it has been shown that *S. oryzae* possesses midgut digestive proteinases ([42], see below), which free amino acids from food proteins (Figure S2.1). These amino acids, accounting for about 12-15% (w/w) of dry matter in cereal grain [43], are the major source of nitrogen, especially the proteinaceous diamino acid glutamine that is very abundant in the prolamins, a group of seed storage proteins characteristic of numerous monocotyledons.

Carbohydrates represent the other essential nutrients for *S. oryzae* to develop in cereal grains enriched with starch. Carbohydrate active enzymes (CAZymes) are involved in the biosynthesis, modification, binding and catabolism of oligo- and polysaccharides. CAZymes are categorized into five major classes: glycoside hydrolases (GH), polysaccharide lyases (PL), carbohydrate esterases (CE), glycosyltransferases (GT) and various auxiliary oxidative enzymes (CAZY Database, [44]). Here, we investigated the genomic basis of specialized phytophagy on cereal grains by *S. oryzae,* through annotation and comparative genomic analyses of major enzymes involved in grain utilization.

#### 3.2 Methods

Peptidase gene families were annotated using the MEROPS database [39]. Predicted protein sequences from all compared species were searched using BLASTP [45] against the MEROPS database with an E-value cutoff of 1E-15. Carbohydrate-active enzymes (cazymes) were annotated using the dbCAN2 meta server [46]. Predicted protein sequences from all compared species were searched using HMMER, DIAMOND and HotPep against the dbCAN HMMdb version 8. Only proteins with a positive hit in each of these three methods were considered as significant.

#### 3.3 Results and discussion

We screened *S. oryzae* for the presence of all known protease gene families and found a total of 769 protease coding genes from 79 peptidase gene families divided among six groups (Asparagine (N = 4), Aspartic (N = 31), Cysteine (N = 121), Metallo (N = 185), Serine (N = 388) and Threonine (N = 26) peptidases) (Table S3.1 and Supplemental Table 5: Table S3.3). The serine peptidase families S9 (N = 166), including prolyl oligopeptidases, and S1 (N = 113), including Trypsins and Chymotrypsins, have by far the highest number of peptidases. Cysteine peptidases of the C1 family were represented by 35 genes belonging mostly to cathepsins L (N = 17) and B (N = 7). An expansion of C1 cysteine peptidase genes has already been described in other coleopteran species such as *T. castaneum*, *T. molitor* or *Leptinotarsa decemlineata* [47–49]. Such expansion of protease encoding genes in *Sitophilus* may be explained by the arsenal of allelochemicals, including protease inhibitors present in the cereal grains leading to an arms race between insects and plants [50–52]. Grains also contain α-amylase inhibitors contributing to the host-plant resistance to insect pests [51]. Indeed, α-amylases are a family of simple carbohydrate (starch)-metabolizing enzymes essential for insect growth particularly for the insect pests of stored grains since starch is the most abundant component of grains. Previous studies have demonstrated the susceptibility of *S. oryzae* to α-amylase inhibitors of wheat [53,54]. In the classification of all carbohydrate-active enzymes (CAZy; [44]), α-amylases are one of the most frequently occurring glycoside hydrolases (GH). Using the dbCAN2 resource [46], we identified 133 GH assigned to 30 families, 116 glycosyltransferases (GT) assigned to 39 families, 32 redox enzymes with auxiliary activities (AA) assigned to 4 families and 10 carbohydrate esterases (CE) assigned to 4 families (Table S3.2 and Supplemental Table 5: Table S3.4). Additionally, 29 proteins with a carbohydrate-binding domain assigned to 5 families were detected including 16 chitin-binding proteins (CBM14). In insects, chitin-binding proteins are particularly found in the midgut peritrophic matrix where they are assumed to mediate the matrix barrier function [55,56]. Interestingly, we found that *S. oryzae* possesses a large array of plant cell wall degrading enzymes (PCWDE) including 23 cellulases (12 GH1, 1 GH9, 8 GH45, 2 GH48), 37 hemicellulases (3 GH2, 3 GH16, 3 GH27, 2 GH30, 8 GH31, 5 GH35, 9 GH38, 4 GH47) and 12 pectinases (6 GH28, 1 GH79 and 5 CE8). Previous work demonstrated that the acquisition of some of these PCWDE may however arise from lateral gene transfer [57–59].

#### Table S3.1. Comparison of the protease genes among coleopteran sequenced genomes.

| | *Sitophilus oryzae* | *Dendroctonus ponderosae* | *Leptinotarsa decemlineata* | *Diabrotica virgifera* | *Anoplophora glabripennis* | *Aethina tumida* | *Tribolium castaneum* |
|---|---|---|---|---|---|---|---|
| **Order, Family** | Coleoptera, Curculionidae | Coleoptera, Curculionidae | Coleoptera, Chrysomelidae | Coleoptera, Chrysomelidae | Coleoptera, Cerambycidae | Coleoptera, Nitidulidae | Coleoptera, Tenebrionidae |
| **Asparagine peptidases** | 4 | 3 | 3 | 3 | 4 | 3 | 3 |
| **Aspartic peptidases** | 31 | 9 | 19 | 68 | 18 | 8 | 25 |
| **Cysteine peptidases** | 121 | 115 | 144 | 154 | 133 | 132 | 117 |
| **Metallo peptidases** | 185 | 198 | 174 | 208 | 190 | 189 | 186 |
| **Serine peptidases** | 388 | 390 | 422 | 505 | 548 | 437 | 415 |
| **Threonine peptidases** | 26 | 21 | 30 | 31 | 24 | 30 | 24 |
| **Unknown peptidases** | 14 | 17 | 12 | 14 | 15 | 20 | 11 |
| **Total** | 769 | 753 | 804 | 983 | 932 | 819 | 781 |

#### Table S3.2. Summary of annotated CAZymes in *S. oryzae*.

| Enzyme class | Enzyme family | Number of proteins in *S. oryzae* |
|---|---|---|
| **Auxiliary Activities (AA)** | AA1 | 2 |
| | AA15 | 3 |
| | AA3 | 17 |
| | AA8 | 10 |
| **Carbohydrate Binding Modules (CBM)** | CBM13 | 8 |
| | CBM14 | 16 |
| | CBM39 | 1 |
| | CBM47 | 2 |
| | CBM48 | 2 |
| **Carbohydrate Esterases (CE)** | CE0 | 3 |
| | CE13 | 1 |
| | CE8 | 5 |
| | CE9 | 1 |
| **Glycoside Hydrolases (GH)** | GH0 | 2 |
| | GH1 | 12 |
| | GH116 | 1 |
| | GH13 | 12 |
| | GH133 | 1 |
| | GH152 | 6 |
| | GH16 | 3 |
| | GH18 | 13 |
| | GH2 | 3 |
| | GH20 | 12 |
| | GH22 | 2 |
| | GH27 | 3 |
| | GH28 | 6 |
| | GH29 | 4 |
| | GH30 | 2 |
| | GH31 | 8 |
| | GH32 | 2 |
| | GH35 | 5 |
| | GH37 | 5 |
| | GH38 | 9 |
| | GH45 | 8 |
| | GH47 | 4 |
| | GH48 | 2 |
| | GH56 | 1 |

| | | |
|---|---|---|
| | GH63 | 1 |
| | GH79 | 1 |
| | GH84 | 1 |
| | GH85 | 1 |
| | GH89 | 2 |
| | GH9 | 1 |
| GlycosylTransferases (GT) | GT1 | 27 |
| | GT10 | 2 |
| | GT105 | 3 |
| | GT13 | 1 |
| | GT14 | 1 |
| | GT16 | 1 |
| | GT2 | 6 |
| | GT20 | 1 |
| | GT21 | 1 |
| | GT22 | 4 |
| | GT23 | 1 |
| | GT24 | 1 |
| | GT25 | 2 |
| | GT27 | 8 |
| | GT29 | 1 |
| | GT3 | 1 |
| | GT31 | 9 |
| | GT32 | 3 |
| | GT33 | 1 |
| | GT35 | 1 |
| | GT39 | 2 |
| | GT4 | 3 |
| | GT41 | 1 |
| | GT43 | 2 |
| | GT47 | 3 |
| | GT49 | 4 |
| | GT54 | 1 |
| | GT57 | 2 |
| | GT58 | 1 |
| | GT61 | 1 |
| | GT64 | 3 |
| | GT65 | 1 |
| | GT66 | 2 |
| | GT68 | 1 |
| | GT7 | 4 |
| | GT76 | 1 |
| | GT8 | 4 |
| | GT90 | 2 |
| | GT92 | 3 |

**Large supplementary tables (Supplemental Table 5):**

**Table S3.3. List of all protease genes annotated in *S. oryzae*.**

**Table S3.4. List of all CAZYme genes annotated in *S. oryzae*.**

### 4. Development

Patrick CALLAERTS

#### 4.1 Introduction

Most of our insight into the gene regulatory networks that control insect embryogenesis and patterning stems from work on the genetic model *D. melanogaster* [60]. Contrary to *D. melanogaster*, which has a *long germ* embryogenesis, most insects have different modes of embryogenesis, namely *short* and *intermediate* embryogenesis [61]. *Short germ* embryogenesis is considered the ancestral mode and in recent years, the red flour beetle, *T. castaneum*, has emerged as an experimentally tractable model to study genetic mechanisms regulating *short germ* embryogenesis [62–64]. Previous work from Tiegs and Murray [65] has shown that *S. oryzae* embryogenesis belongs to the *short germ* mode and is very similar to *T. castaneum*. Therefore, we used the combined insight into the developmental gene regulatory networks of *D. melanogaster* and *T. castaneum* as the basis for the current annotation of developmental genes in *S. oryzae*.

#### 4.2 Methods

In order to annotate developmental genes encoded in the *S. oryzae* genome, we used reciprocal best BLASTP analyses. We first took advantage of the automatic annotation of the *S. oryzae*'s genome to identify proteins with similarity to developmental gene-encoded proteins of interest. These putative *S. oryzae* proteins were then compared (BLASTP) against the *D. melanogaster* reference proteins. In a complementary analysis, a complete collection of *D. melanogaster* developmental gene-encoded proteins was compared against the *S. oryzae* genome. When both BLASTP analysis yielded the same *S. oryzae*/*D. melanogaster* homolog protein pairs, these were annotated as such. In cases where there were discrepancies between the two, manual curation was done with focus on unique protein identifying amino acids to assign the most likely homolog. The other putative homologs were then listed as "homolog-like".

#### 4.3 Results and discussion

Homologs for all major signaling pathway genes were identified in *S. oryzae* (Supplemental Table 5: Table S4.1). The pathways that were annotated include Wnt signaling (core canonical, planar cell polarity, other pathway-associated genes), TGFbeta signaling (BMP and Activin branches), Notch signaling, Hippo signaling, RTK signaling (core, EGFR signaling and regulators, FGFR signaling, PVR signaling), JAK-STAT signaling, JNK pathway signaling, and Hedgehog signaling. We did not find a homolog for *naked cuticle* (Wnt signaling). In addition, we observed quite frequently that homologs for signaling pathway ligands could not unequivocally be identified. Single EGFR (*Keren*) and FGFR (*FGF-like*) ligands were identified. No homologs were identified for *gurken* and *vein* (EGFR signaling), *unpaired 1-3* and *eye transformer* (JAK-STAT signaling). Given the conservation of the receptors and the signaling cascades, this finding suggests that not the primary sequence, but conceivably the tertiary structure of the ligands is essential for binding to the relevant receptors. Consequently, more extensive sequence divergence may occur complicating identification of ligand homologs.

Regarding the embryonic patterning genes, we were unable to identify homologs for a few key genes of different coordinate gene groups as defined in *D. melanogaster*. We found no homologs for *bicoid* and *swallow* (anterior group), for *oskar* (posterior group) and for *trunk* (terminal group). A putative homolog for *nanos* was detected albeit with low support, thus awaiting further confirmation. In summary, the annotated *S. oryzae* genes suggest that anteroposterior patterning in *S. oryzae* may be regulated in a manner very similar to what is observed in *T. castaneum*. The absence of *trunk*, a terminal patterning gene, from the *S. oryzae* genome is remarkable in light of the fact that it is found in the *T. castaneum* genome [66]. All genes involved in dorsoventral patterning are however highly conserved in *S. oryzae*.

Many of the genes implicated in *D. melanogaster* oogenesis, and deposition and localization of maternal gene products seem conserved in *S. oryzae* suggesting that some of the functions could be conserved as well. The absence of *oskar*, a gene with important roles in *D. melanogaster* germline development, is consistent with the fact that maternally synthesized polar granules and early specification germ cells (as seen in *D. melanogaster*) are missing from many insect species, including *T. castaneum* [47]. Instead of two genes, we found a single homolog for *wunen* and *wunen2*, genes that act during germ cell migration [67].

Regarding segmentation, the *S. oryzae* genome harbours all gap, segment polarity, and pair-rule genes except *fushi tarazu*. The conservation of all these genes suggests that the segmentation mechanisms of *S. oryzae* do not differ from other arthropods. An interesting novelty was identified in *T. castaneum* in the form of the gene *mille-pattes,* which we also identified in *S. oryzae*. This gene was classified as a gap gene because of its cross-regulatory interactions with other gap genes. Remarkably, this gene encodes four peptides, and its knockdown leads to transformation of abdominal segments into thoracic segments [68]. As expected, a full complement of homeotic HOX genes was identified comparable to what is found in *D. melanogaster* and *T. castaneum* [69,70].

All signaling pathways and key transcription factors implicated in *D. melanogaster* embryonic organogenesis are conserved in *S. oryzae*. Even though our insight in *T. castaneum* embryonic organogenesis is currently still limited, these results suggest that many of the fundamental genetic mechanisms that control embryonic organogenesis may be conserved in *S. oryzae*. One remarkable observation is that we did not find homologs for *miranda* and *partner of numb*, two genes implicated in asymmetric division of neuroblasts during nervous system development. Albeit speculative, this could indicate that the mode of neuroblast division in *S. oryzae* embryogenesis is distinct from what is seen in *D. melanogaster*.

Appendage development in *T. castaneum* does not depend on imaginal discs as in *D. melanogaster*. Nevertheless, development of appendages (antenna, mouth parts, legs) is regulated by essentially the same developmental pathways. These include *wingless* and *dpp* signaling pathways, and the transcription and nuclear factors *homothorax*, *extradenticle*, *distalless*, *aristaless,* and *dachshund* [69,71,72]. All these genes are conserved in *S. oryzae* suggesting that the gene regulatory networks controlling appendage development are also conserved.

All genes of the retinal determination gene network are conserved in *S. oryzae*, so it seems very likely that early head and eye patterning will be strongly conserved. The fact that we could not identify a *sevenless* homolog may indicate that genes and gene networks relevant later in photoreceptor development may have undergone changes.

Insect size and developmental transitions are controlled by insulin and mTOR signaling on the one hand and ecdysteroid and juvenile hormone signaling on the other hand [73,74]. We have identified homologs of all essential genes, which would be consistent with them having very similar roles in *S. oryzae*. The only clear difference is at the level of insulin signaling ligands. Our analysis reveals strong support for three homologs of Ilp2 and Ilp5, whereas additional Ilp-like peptides were not found. However, this is inconclusive as it may well be due to significant sequence divergence, similar to what was seen when comparing *Acyrthosiphon pisum* with *D. melanogaster* [75].

One final observation was that several genes for which two homologs are present in the *Drosophila* genome only have a single counterpart in the *S. oryzae*'s genome (*Drosophila* genes *echinoid*/*friend of echinoid*, *thisbe*/*pyramus*, *bric-a-brac1/2*, *ladybird early*/*late*, *wunen*/*wunen-2*, *zerknültt*/*zen2*, *teashirt*/*tiptop*). We speculate that many of these may reflect duplication events specific for the *Drosophila* lineage and that the gene complement as observed in *S. oryzae* is the more ancestral status. Interesting in that regard is also that the *Tribolium* genome encodes two *zerknüllt* homologs (*Tc-zen1* and

*Tc-zen2*) [76] suggesting that further independent duplication events can occur within the Coleoptera. It can be expected that results of the iBeetle genome-wide RNAi screen to identify gene functions in embryonic and postembryonic development and in physiology will facilitate a more detailed functional annotation of the *S. oryzae* developmental genes [62,77].

**Large supplementary table (Supplemental Table 5):**

**Table S4.1. List of all development related genes in *S. oryzae.***

### 5. Cuticle protein genes

Carlos VARGAS-CHAVEZ

#### 5.1 Introduction

*S. oryzae* represents one of the greatest threats to postharvest agricultural products in terms of resistance to insecticides and the cuticle represents the first physical barrier to topical insecticides [78]. Like other beetles, weevils have a strong cuticle that protects them from the penetration of toxins and/or pathogens, physical trauma and water loss, while also displaying a wide range of mechanical properties. The cuticle is used to form both a strong and rigid armor, via modified wings called elytra, and a light and flexible yet resistant pair of wings that enable flight [79]. A vast diversity of cuticular proteins (CPs) has been documented in arthropoda ranging from a repertoire of 66 to 98 CPs in hymenopterans to between 118 and 305 CPs in dipterans [80]. In the case of coleopterans, the CPs repertoire has been explored in a few species.

#### 5.2 Methods

We downloaded the full proteomes for seven species from NCBI genome database: *S. oryzae* - GCF_002938485.1, *D. ponderosae* - GCF_000355655.1, *A. glabripennis* - GCF_000390285.2, *A. tumida* - GCF_001937115.1, *T. castaneum* - GCF_000002335.3, *D. virgifera* - GCF_003013835.1, *L. decemlineata* - GCF_000500325.1 and used the CUTPROTFAM-PRED web server [80] to identify and classify CPs in the full proteomes.

#### 5.3 Results and discussion

Besides detoxification strategies, the cuticle represents a physical barrier to topical insecticides allowing an increased tolerance to insecticides [78], therefore we explored the cuticular protein repertoire of *S. oryzae* and identified 152 CPs (Table S5.1). In comparison, we identified between 135 and 256 in other beetles. While *S. oryzae* was in the median of the total number of CPs, it had an increased number of members of the CPAP1 family (25 in *S. oryzae* versus a median of 10 in all assessed beetles). While a direct link between these proteins and an increased tolerance to insecticides has not been described, it has been shown that some of these proteins are essential for the pupal-to-adult molt in *T. castaneum* [81,82]. Additionally, given the divergent expression patterns in the different cuticle-forming tissues, these proteins might be involved in the development of cuticular tissues particular to *S. oryzae* or other weevils such as the rostrum. We did not observe that the total number of CPs followed the taxonomy of the beetles but instead it might be an adaptation to their diverse lifestyles.

**Table S5.1. Comparison of the cuticular protein genes among coleopteran sequenced genomes.**

| | ***Sitophilus oryzae*** | ***Dendroctonus ponderosae*** | ***Leptinotarsa decemlineata*** | ***Diabrotica virgifera*** | ***Anoplophora glabripennis*** | ***Aethina tumida*** | ***Tribolium castaneum*** |
|---|---|---|---|---|---|---|---|
| **Order, Family** | Coleoptera, Curculionidae | Coleoptera, Curculionidae | Coleoptera, Chrysomelidae | Coleoptera, Chrysomelidae | Coleoptera, Cerambycidae | Coleoptera, Nitidulidae | Coleoptera, Tenebrionidae |
| **CPAP1** | 25 | 12 | 14 | 18 | 12 | 15 | 15 |
| **CPAP3** | 7 | 10 | 10 | 12 | 9 | 11 | 7 |
| **CPCFC** | 3 | 4 | 1 | 3 | 5 | 6 | 2 |
| **CPF** | 2 | 2 | 4 | 2 | 3 | 4 | 5 |
| **CPLCA** | 0 | 0 | 0 | 0 | 1 | 0 | 1 |
| **CPLCG** | 1 | 0 | 1 | 1 | 1 | 5 | 1 |
| **CPR_RR-1** | 49 | 35 | 53 | 47 | 64 | 103 | 42 |
| **CPR_RR-2** | 59 | 64 | 86 | 92 | 62 | 97 | 61 |
| **Tweedle** | 6 | 8 | 8 | 5 | 7 | 15 | 3 |
| **Total** | 152 | 135 | 177 | 180 | 164 | 256 | 137 |

### 6. Innate Immune system

Carole VINCENT-MONEGAT, Carlos VARGAS-CHAVEZ, Nicolas PARISOT, Justin MAIRE, BERANGER Louis, BONNAMOUR Aymeric, ZAMOUM Waël, Florent MASSON, Aurélien VIGNERON, Anna ZAIDMAN-REMY

#### 6.1 Introduction

The innate immune system is the first line of defence against invasion by microbial pathogens in both insects and mammals [83]. The innate immune system of insects is divided into humoral defenses that include the production of soluble effector molecules, and cellular defenses such as phagocytosis and encapsulation that are mediated by hemocytes [84]. Here, we focused on the humoral immune responses including the production of AntiMicrobial Peptides (AMPs) that protect against a broad array of infectious agents, such as bacteria, fungi, viruses and even eukaryotic parasites [85]. These defense responses are activated by Pattern Recognition Receptors (PRRs) that detect and bind to conserved microbial structures called Microbial-Associated Molecular Patterns (MAMPs), such as bacterial lipopolysaccharide (LPS) and peptidoglycans (PG). The three main signaling transduction pathways leading to the production of AMPs are Toll, JAK/STAT and IMmune Deficiency (IMD) [86].

Several holometabolous insects, including *D. melanogaster* (Diptera) [87], *A. gambiae* (Diptera) [88], *G. morsitans* (Diptera) [89], *T. castaneum* (Coleoptera) [90] and *M. sexta* (Lepidoptera) [91] are traditional models for genomic and functional investigations of insect innate immunity. The data obtained in these various models have attested the conservation of innate immunity in holometabolous insects so far. However, compared to these models, *S. oryzae* presents a specificity due to its symbiotic relationship with the nutritional endosymbiont *S. pierantonius*. Several sap-feeding insects harbouring endosymbionts have, for instance, lost the entire signaling IMD pathway, supposedly allowing them to tolerate the permanent association with their Gram-negative endosymbiont [92,93]. In this study, we annotated the immune-related genes of *S. oryzae* with the objective of determining whether the main pathways regulating humoral responses in holometabolous insects were conserved in *S. oryzae*, or whether its association with a Gram-negative endosymbiont had led to a significant evolution of its immune gene repertoire, at the genomic level.

#### 6.2 Methods

For the annotation of immune genes encoded in the *S. oryzae* genome, we used bidirectional Blastp analysis. Briefly, the set of predicted *S. oryzae* proteins was compared to the immune gene database coming from 4IN (http://4in.cycadsys.org), an interactive database for Insect Innate Immunity, that we developed in the laboratory. Only insects whose genome has been completely sequenced, annotated, and on which preliminary studies of the innate immune response have been performed, were selected: *Aedes aegypti, Anopheles gambiae, Acyrthosiphon pisum, Bombyx mori, Camponotus floridanus, Drosophila melanogaster, Dendroctonus ponderosae, Glossina morsitans, Manduca sexta, Nasonia vitripennis, Pediculus humanus, Plutella xylostella, Solenopsis invicta and Tribolium castaneum*. Hits showing ≥ 30 % of identity over at least 100 amino acids and an E-value ≤ 5e-30 were considered as significant. Due to their short length, AMPs were annotated using a dedicated procedure. Briefly, the entire set of predicted *S. oryzae* proteins was blasted (Blastp) against the AMPs database coming from DRAMP [94] and 4IN databases (http://4in.cycadsys.org). A manual curation of putative *S. oryzae* AMPs was then performed, using the physico-chemical properties of each candidate. Additionally, we blasted (tBlastn) a set of well-known insect AMPs (from DRAMP and 4IN) to the assembly.

#### 6.3 Results and discussion

In the fruit fly *D. melanogaster*, microbial recognition leads to signal production via three major pathways: Toll, IMmune Deficiency (IMD) and Janus Kinase/Signal Transducers and Activators of Transcription (JAK-STAT), each pathway being activated in response to particular pathogens, viral infection or tissue damage [95].

A graphical summary of these pathways as conserved in *S. oryzae* is reported in Figure S6.1.

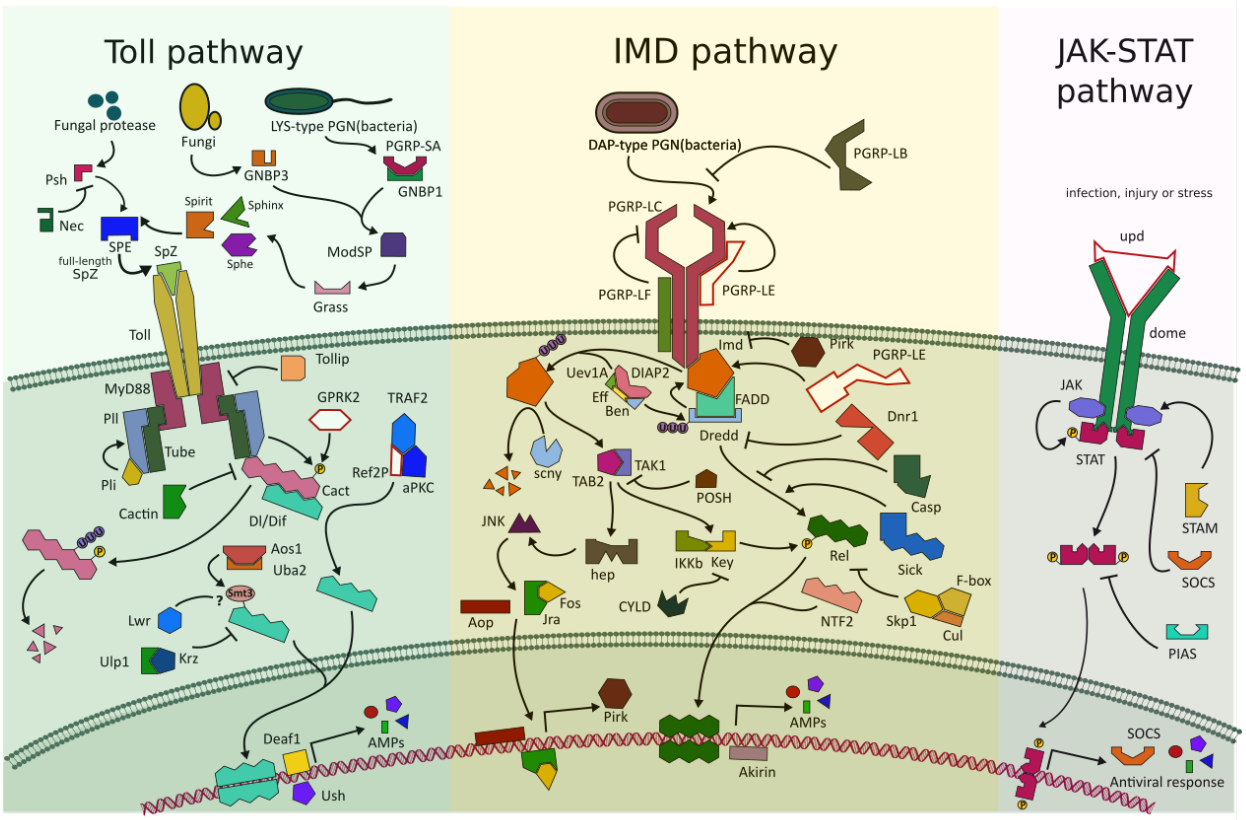


**Figure S6.1. Toll, IMD and JAK-STAT pathways of *S. oryzae***. All members were manually verified by blast. The name of each member corresponds to *Drosophila* protein names. Missing members are shown in white.

The IMD pathway is triggered by bacterial diaminopimelate (DAP)-type PG recognition, a cell wall component of most Gram-negative bacteria as well as Gram-positive bacteria from the *Bacillus* and *Listeria* genera [87,96]. In *D. melanogaster,* the main receptors associated with this pathway are the PeptidoGlycan Recognition Proteins (PGRP)-LC and PGRP-LE, which bind DAP-type PG and initiate the signalling cascade by activating the Imd protein [97–99]. Specifically, the transmembrane receptor PGRP-LC recognizes circulating polymeric DAP-type PG as well as a monomeric fragment, the tracheal cytotoxin (TCT) [100,101], while TCT located within insect cells is detected by the intracellular PGRP-LE [102–104]. In *S. oryzae*, a gene encoding the receptor PGRP-LC was identified, and demonstrated to be essential for circulating-PG recognition and consequent IMD pathway activation [105]. However, no *pgrp-le* ortholog was found in the genome. Contrasting with many members of the PGRP family that are relatively well conserved across insects, PGRP-LE has a more complexe profil of conservation. While it has so far not been identified in species such as *A. gambiae*, *B. mori, G. morsitans* or *A. mellifera,* it is present in the genomes of several coleopteran, such as *T. castaneum* [90] and *T. molitor* [106]. In the case of endosymbiotic insects, including tsetse fly that harbors the intracellular *Wigglesworthia glossinidia* [89] and *S. oryzae* (this work and [105]), the lack of these intracellular TCT receptors could be a common feature that avoids the induction of local immune responses against endosymbionts.

Once bound to PGN, the receptor likely dimerizes or multimerizes and an intracellular signal is transmitted to the adaptor protein Imd. Imd contains a death domain that recruits FADD (TAK1 activator) and Dredd (a caspase). Active TAK1 triggers the JNK pathway (through Hep, JNK, Jra, Fos, AOP) and Relish (rel) phosphorylation (through IKKb and Kenny (key)), which in turn upregulates AMP-encoding gene expression [99]. In *S. oryzae*, all signaling members were identified, consistently with a recent study showing the IMD pathway is functional [107] (Table S6.1).

**Table S6.1. Identification of the main genes belonging to the IMD pathway in *S. oryzae* based on the annotation available for insects**. Only best hit per gene were reported in this Table (for a comprehensive list of orthologs, see Supplemental Table 5: Table S6.2)

| Query | Name | Pathway | ID sequences producing best significant alignments | E-value |
|---|---|---|---|---|
| **LOC115876451** | akirin | IMD pathway | GLEAN_14377 | 3.33E-84 |
| **LOC115882365** | ben | IMD pathway | GLEAN_01755 | 1.75E-109 |
| **LOC115880562** | cad | IMD pathway | GLEAN_07576 | 3.22E-43 |
| **LOC115888419** | casp | IMD pathway | ENN74653 | 0 |
| **LOC115874148** | Cul | IDM pathway | ENN72810 | 0 |
| **LOC115877056** | CYLD | IMD pathway | ENN80894 | 0 |
| **LOC115891591** | Diap2 | IMD pathway | GLEAN_01189 | 0 |
| **LOC115889325** | dnr1 | IMD pathway | ENN72057 | 0 |
| **LOC115878704** | Dredd | IMD pathway | ENN79559 | 3.75E-136 |
| **LOC115880045** | eff | IMD pathway | NV14758 | 1.54E-108 |
| **LOC115876310** | Fadd | IMD pathway | ENN77263 | 2.00E-24 |
| **LOC115884520** | IKKb | IMD pathway | ENN74989 | 2.85E-155 |
| **LOC115876629** | imd | IMD pathway | GLEAN_10851 | 3.87E-36 |
| **LOC115879224** | key | IMD pathway | GLEAN_12666 | 0 |
| **LOC115885445** | Npc2 | IMD pathway | ENN74610 | 3.69E-56 |
| **LOC115885165** | Ntf2 | IMD pathway | GLEAN_12876 | 4.48E-83 |
| **LOC115878866** | pirk | IMD pathway | AJG05482.1 | 0 |
| **LOC115890600** | POSH | IMD pathway | XP_019769090.1 | 0 |
| **LOC115886763** | rel | IMD pathway | GLEAN_11191 | 0 |
| **LOC115885873** | scny | IMD pathway | GLEAN_02978 | 0 |
| **LOC115888857** | sick | IMD pathway | XP_019772630.1 | 0 |
| **LOC115885781** | Tab2 | IMD pathway | ENN76471 | 6.87E-142 |
| **LOC115885735** | Tak1 | IMD pathway | ENN76602 | 0 |
| **LOC115878176** | UEV1a | IMD pathway | GLEAN_10380 | 2.96E-100 |

The Toll pathway is involved in the response against some Gram-positive cocci and against fungi. In *D. melanogaster*, the pathway is triggered by extracellular Lys-type PG recognition *via* PGRP-SA, GNBP1, by fungal β-glucans recognition *via* GNBP3, and by the detection of fungal protease activity by Persephone [108–110], followed by several protease cascades converging in the activation of the Späetzle Processing Enzyme (SPE). Several candidates are proposed for the serine-proteases involved in the cascades in *S. oryzae* (ModSP, Grass, Sphinx, Spirit, Spheroid (Sphe), SPE and Persephone (Psh)), however establishing a clear one-to-one relation was not possible for some of them [111]. Following SPE activation, Spätzle (Spz) is cleaved, allowing it to bind the extracellular region of the Toll receptor, which initiates the intracellular signaling cascade: dimerization of Toll receptors then homotypic interaction with MyD88. MyD88, Tube and Pelle (Pll) bind together to form a signaling complex via their death domains. Then, the complex induces the translocation of their downstream transcription factors, Dif and Dorsal (Dl) into the nucleus, resulting in the activation of AMP-encoding genes upregulation [112]. Both Spätzle-like proteins and Toll receptors exhibit strong amplification in *S. oryzae* (10 and nine orthologs respectively), as described in other insects [94,95,113] (Supplemental Table 5: Table S6.3). On the other hand, most of the components of the intracellular pathway are highly conserved and only Gprk2, Ref(2)P and Dif appear to be missing (Table S6.4 and Figure S6.1). Similar to *C. floridanus* and *A.mellifera* [114], a single gene encoding Dorsal is present in *S. oryzae*, but no ortholog of Dif was found.

**Table S6.4. Identification of the main genes belonging to the Toll pathway in *S. oryzae* based on the annotation available for insects.** Genes with no matching with *D. melanogaster* and other insect homolog are shown in red. Only best hit per gene were reported in this Table (for a comprehensive list of orthologs, see Supplemental Table 5: Table S6.2)

| Query | Name | Pathway | ID sequences producing best significant alignments | E-value |
|---|---|---|---|---|
| **LOC115890560** | aos1 | TOLL pathway | ENN75245 | 0 |
| **LOC115883435** | aPKC | TOLL pathway | GB47743 | 0 |
| **LOC115887424** | cact | TOLL pathway | ENN76157 | 1.36E-114 |
| **LOC115889882** | cactin | TOLL pathway | ENN71807 | 0 |
| **LOC115885191** | Deaf1 | TOLL pathway | ENN76751 | 0 |
| | dif | TOLL pathway | | |
| **LOC115888177** | dl | TOLL pathway | GLEAN_07697 | 1.77E-161 |
| **LOC115886736** | GNBP | TOLL pathway | ENN83076 | 0 |
| | GPRK2 | TOLL pathway | | |
| **LOC115889345** | grass | TOLL pathway | GLEAN_09090 | 3.00E-115 |
| **LOC115876845** | krz | TOLL pathway | GLEAN_01639 | 0 |
| **LOC115881934** | lwr | TOLL pathway | ENN74894 | 1.30E-117 |
| **LOC115890075** | mbo | TOLL pathway | ENN71557 | 0 |
| **LOC115876215** | modSP | TOLL pathway | ENN81797 | 0 |
| **LOC115884632** | myd88 | TOLL pathway | ENN75680 | 1.51E-112 |
| **LOC115884774** | pli | TOLL pathway | GLEAN_09672 | 0 |
| **LOC115877325** | pll | TOLL pathway | ENN77083 | 0 |
| **LOC115881027** | psh | TOLL pathway | ENN71261 | 1.00E-121 |
| | ref(2)P | TOLL pathway | | |
| **LOC115889684** | smt3 | TOLL pathway | ENN79776 | 5.00E-58 |
| **LOC115888508** | Spe | TOLL pathway | AAEL000028 | 1.00E-88 |
| **LOC115877093** | sphinx | TOLL pathway | ENN74004 | 1.55E-106 |
| **LOC115883682** | spirit | TOLL pathway | PHUM452840 | 9.25E-91 |
| **LOC115885648** | Spz | TOLL pathway | ENN81604 | 9.38E-41 |
| **LOC115888019** | tamo | TOLL pathway | GLEAN_15946 | 1.98E-171 |
| **LOC115881356** | toll | TOLL pathway | GLEAN_04438 | 0 |
| **LOC115888469** | tube | TOLL pathway | ENN70294 | 0 |
| **LOC115876397** | tollip | TOLL pathway | GLEAN_07125 | 1.05E-144 |
| **LOC115878153** | uba2 | TOLL pathway | ENN71609 | 0 |
| **LOC115877476** | ulp1 | TOLL pathway | ENN83368 | 0 |
| **LOC115885819** | Ush | TOLL pathway | GLEAN_13689 | 0 |

The JAK-STAT pathway is involved in pathogen surveillance mechanisms in insects, especially in hematopoiesis and cellular immunity, viral response, and gut immunity [115]. In *D. melanogaster*, extracellular proteins of the Unpaired family bind to Domeless (dome), causes receptor dimerization, and recruits STAM and JAK, which in turn phosphorylates itself and then STAT. We did not find in *S. oryzae* an Unpaired (upd) ortholog, consistently with the data in *M. sexta* and *T. castaneum* [90]. After phosphorylation, the STAT dimer translocates into the nucleus to induce antiviral gene expression. SOCS (a JAK inhibitor) and PIAS (protein inhibitor of activated STAT) may down-regulate the pathway. Except for the ligand, orthologs of all the pathway components are present in *S. oryzae* (Figure S6.1). While this absence of Unpaired (Upd) may mean that the JAK-STAT pathway is not functional in *S. oryzae*, we cannot exclude that the non-identification of an ortholog of the ligand Unpaired in the genome of *S. oryzae* could result from high variation in the sequence of the cytokine-like protein, as suggested in Zou *et al.* 2007 [90].

In *D. melanogaster,* immune signaling triggers the production of a multitude of effectors, so-called immune effectors. In the cereal weevil, three main groups of immune effectors were identified: AMPs, lysozymes and thaumatins. AMPs have small molecular weights and broad-spectrum activities against bacteria, fungi and viruses [116]. They generally consist of twelve to fifty amino acids and are divided into subgroups based on their amino acid composition and structure. The hydrophobic part of the molecule generally covers more than 50% of amino acids residues. The secondary structure of AMPs follows several categories. For example, we distinguished the family of Cecropins, characterized by linear peptides with alpha-helix that lack cysteine residues, the family of Defensins, characterized by six to eight conserved cysteine residues, a stabilizing array of three or four disulfide bridges and three domains consisting in a flexible amino-terminal loop, and the family of Glycine-rich AMPs characterized by an overrepresentation of Proline and/or Glycine residues. Sixteen sequences matching five different categories of AMPs were identified using the automated genome annotation and supplementary manual annotation (Table S6.5, see Methods). We found well-known peptides (cecropin, diptericin, sarcotoxin, coleoptericin and defensin) as well as many members of the Glycine-rich AMPs, including a new candidate probably specific to our model (Glycine-rich AMPs-like).

**Table S6.5. Identification of the main genes belonging to immune effectors in *S. oryzae* based on the annotation available for insects.**

| GeneID | Name | Category of immune effector |
|---|---|---|
| **LOC115875079** | Acanthoscurrin-1-like | a novel Gly-rich AMPS |
| **LOC115875523** | Acanthoscurrin-2-like | a novel Gly-rich AMPS |
| **LOC115875524** | Acanthoscurrin-2-like | a novel Gly-rich AMPS |
| **LOC115891322** | Cecropin | Alpha-helical linear AMPs |
| **LOC115886926** | CAMP-like | Cathelicidins |
| **LOC115888712** | Defensin | Disulfide bonds and beta-hairpin AMPs |
| **LOC115874620** | Coleoptericin A | Glycine-rich AMPs |
| **LOC115874703** | Coleoptericin B | Glycine-rich AMPs |
| **LOC115877460** | Diptericin-1 | Glycine-rich AMPs |
| **LOC115877462** | Diptericin-2 | Glycine-rich AMPs |
| **LOC115877463** | Diptericin-3 | Glycine-rich AMPs |
| **LOC115877465** | Diptericin-4 | Glycine-rich AMPs |
| **LOC115877461** | Diptericin-like partial | Glycine-rich AMPs |
| **LOC115884869** | holotricin-3-like | Glycine-rich AMPs |
| **LOC115888387** | Sarcotoxin | Glycine-rich AMPs |
| **LOC115884866** | Glycine-rich AMPs-like | Glycine-rich AMPs |
| **LOC115875419** | putative defense protein Hdd11 | Insect defense protein |
| **LOC115883884** | Luxuriosin | Serine protease inhibitor |
| **LOC115885658** | Thaumatin-like protein | Thaumatin |
| **LOC115885681** | Barietin | Toxin |

Lysozymes also act against bacteria, by targeting PGN, especially of Gram-positive bacteria. Lysozymes are muramidases hydrolysing the β-1,4-glycosidic linkage between N-acetylmuramic acid and N-acetylglucosamine in the PG [117]. They are well known immune effectors, widespread in a huge variety of organisms from viruses to animals and plants. Insect lysozyme was described for the first time as the main antimicrobial factor by Mohrig and Messner [118] and the vast majority of known insect lysozymes are both c-type and i-type lysozymes [119]. Two c-type lysozymes [120] and three i-type lysozymes [121] were identified in *S. oryzae* (Table S6.6).

**Table S6.6.** Identification of the lysozyme-like in *S. oryzae* based on the annotation available for insects.

| Gene ID | Superfamily | Description |
|---|---|---|
| **LOC115882935** | Lyz-like Superfamily | C-type invertebrate lysozyme |
| **LOC115884241** | Lyz-like Superfamily | I-type lysozyme |
| **LOC115889788** | Lyz-like Superfamily | C-type invertebrate lysozyme |
| **LOC115891623** | Lyz-like Superfamily | I-type lysozyme |
| **LOC115891647** | Lyz-like Superfamily | I-type lysozyme |

Thaumatins are described as antifungal proteins with a glucanase function [122], although their precise mechanism of action is still unknown. A thaumatin sequence had been identified earlier in *S. oryzae* [123,124], and is confirmed in this study.

We also identified a new putative immune effector named Viresin-like (Table S6.7). These peptides have been characterized in *Heliothis virescens* as a defense response to bacterium [125]. In other insects, these sequences are characterized as homologs to insect pheromone-binding family A10/OS-D.

**Table S6.7. List of genes encoding a viresin-like protein in *S. oryzae.***

| GeneID | Name |
|---|---|
| **LOC115884092** | viresin-1-like |
| **LOC115884091** | viresin-2-like |
| **LOC115884088** | viresin-3-like |
| **LOC115880538** | viresin-4-like |
| **LOC115877659** | viresin-5-like |
| **LOC115877658** | viresin-6-like |
| **LOC115877656** | viresin-7-like |

In conclusion, the whole-genome analyses have revealed a high conservation of key cellular and humoral immune pathways in *S. oryzae*. We identified 840 immune related genes belonging to several categories including microbial recognition, signalling pathways (Toll, JAK-STAT, IMD and JNK), AMPs, phagocytosis, melanisation, encapsulation, cytoskeleton immune proteins, antiviral defence, coagulation, hematopoiesis and other immune responses (Supplemental Table 5: Table S6.2).

Many studies on hemimetabolous species living in association with symbiotic bacteria have described highly degraded immune signaling pathways especially the IMD pathway, for example in aphids, whiteflies, psyllids, triatomines, Pediculidae or Cimicidae [92,93,126–128]. It has been proposed that in their long co-evolution with endosymbiotic bacteria, these hemimetabolous species have lost several genes of the IMD pathway, which could participate in the preservation of their endosymbionts from the host immune responses.

However, this hypothesis may need to be refined in light of various recent studies. First, in some species the loss of the IMD pathway had to be re-evaluated. For example, while it was first claimed that the IMD pathway was largely depleted from the genome of *Rhodnius prolixus* (hemipteran), Salcedo-Porras *et al.* have recently identified most of the missing orthologs of the IMD pathway and demonstrated that the IMD pathway is present and inducible in this species [129]. Hence, the authors suggested that the evolution of tolerance mechanisms towards bacterial symbionts in hemimetabolous insects does not systematically involve the depletion of immune-related genes. Other tolerance mechanisms, yet to be identified, must ensure the maintenance of the extracellular obligatory nutritional symbiont of *R. prolixus*, *Rhodococcus rhodnii* [130].

Second, the depletion of the IMD pathway in hemimetabolous insects is sometimes un-correlated with the presence of obligate symbionts. For example, *Phylloxera* lacks obligate symbionts, and its genome apparently also lacks an intact IMD pathway; although the possibility that divergent genes would participate in a functional pathway, as shown for *R. prolixus* [131], remains open. If confirmed in this species and in other hemimetabolous species that do not require obligate symbionts, the loss of the IMD pathway could be correlated to other adaptations of these insects to their environment. For instance it was proposed that feeding on diets poor in bacteria, such as plant phloem or mammalian blood, alleviated the evolutive pressure on the IMD pathway. In contrast, the Toll pathway would be better conserved due to its essential function during embryonic development. Higher conservation of the Toll pathway could also reflect an acute pressure of fungal infections on some species, *e.g.* species such as *S. oryzae* living on cereal grains where fungi are present in high quantity.

Last, other hypotheses have been proposed to explain the loss of IMD in many hemimetabolous species and not in holometabolous insects, such as a functional merging of the Toll and IMD pathways in hemimetabolous species [132], or an important function of IMD during metamorphosis in holometabolous species, which would maintain pressure on the conservation of a functional IMD pathway along these species' evolution.

In contrast to many hemimetabolous insects, we show here that the IMD immune signaling pathway is complete in *S. oryzae.* This is consistent with previous work on holometabolous insects living in association with endosymbiotic bacteria, such as the carpenter ant *C. floridanus* [114] and the tsetse fly *G. morsitans* [133]. Notably, while the majority of the immune repertoire is present in *C. floridanus*, a low number of PGN recognition proteins and AMPs have been identified, although in this case this could also reflect a lower pressure on immune genes due to the existence of protective social immunity. This suggests that the adaptation has been selected at the level of these few specific genes, resulting in the adaptation of the pathway to the endosymbiotic situation (and/or to the presence of social immunity in the case of ants), instead of its total inactivation.

Similarly, in *Sitophilus*, previous studies have shown that the AMP ColA has evolved toward a function of endosymbiont control through bacterial growth inhibition, thereby preventing endosymbiont escape out of the bacteriome [134]. Interestingly, we demonstrated that the expression of this AMP in the bacteriome was regulated by the IMD pathway, revealing a function of this pathway not only in the immune response to pathogenic infection, but also in the control of endosymbionts [107]. Imd was shown to also regulate the expression of *pgrp-lb*, which we showed to be important for the maintenance of homeostasis in the context of endosymbiosis [135]. Intriguingly, although *pgrp-lb* is well conserved in many insects, its specific function in the cereal weevil immune homeostasis relies on specific isoforms generated by alternative splicing and additional exons that are not shared with other species. Hence, a more complete understanding of this “tweaking” of Imd in an endosymbiotic context will require both a global knowledge of the genes conserved in the genome in various species, but also a finer analysis of the evolution of their regulatory and coding sequences in specific associations.

**Large supplementary tables (Supplemental Table 5):**

**Table S6.2. List of immunity-related genes in *S. oryzae*.**

**Table S6.3. Description of GNBP, Spz, Toll and Serine protease orthologs of immune activators of Toll pathway.**

### 7. Detoxification and Insecticide resistance

Nicolas PARISOT

#### 7.1 Introduction

Fumigation using phosphine, hydrogen phosphide gas ($PH_3$), is by far the most widely used treatment for the protection of stored grains against insect pests due to its ease of use, low cost and universal acceptance as a residue-free treatment [136,137]. However, high-level resistance to this fumigant has been reported in *S. oryzae* from different countries [138–146]. While early studies demonstrated that only 5% of tested samples were resistant to phosphine [147], the frequency of resistance has reached more than 75% in developing countries and up to 100% in Brazil [148].

Insect resistance to phosphine poses a serious threat to effective pest management as there is currently no practical replacement for phosphine that can match its range of advantages. Thus, understanding of the molecular basis underlying insecticide resistance is of utmost importance.

#### 7.2 Methods

Gene families associated with insecticide resistance and detoxification were identified by searching for matches to relevant InterPro [149], Pfam [150] and PROSITE [151] domains in the available coleopteran Cyc databases from ArthropodaCyc [29]. The gene families examined included cytochrome P450s, carboxyl/choline esterases, glutathione S-transferases, UDP-glycosyltransferases, ABC transporters, Cys-loop ligand-gated ion channels and voltage-gated sodium channels. Protein domains used for the searches are listed in Supplemental Table 5: Table S7.2.

#### 7.3 Results and discussion

*S. oryzae* harbors a large arsenal of genes associated with detoxification and resistance to insecticide and more generally to toxins (including plant allelochemicals). These include the major detoxifying enzymes and xenobiotic transporters (cytochrome P450s (CYPs) and carboxyl/choline esterases (CCEs) in phase I direct metabolism, glutathione S-transferases (GSTs) and UDP-glycosyltransferases (UGTs) in phase II conjugation, and ATP-binding cassette (ABC) transporters in phase III excretion) and the known receptors for the main groups of insecticides (Cys-loop ligand-gated ion channels and voltage-gated sodium channels) (Table S7.1 and Supplemental Table 5: Table S7.2). CYPs are a superfamily of enzymes (monooxygenases) that have a highly diverse array of functions including synthesis and metabolism of hormones and pheromones, as well as detoxification through metabolic breakdown of exogenous substrates such as insecticides [152]. Altogether, the results suggest that the *S. oryzae* repertoire of detoxification and insecticide resistance genes is similar to other coleopterans. Studies of the molecular bases of insecticide resistance mechanisms in *S. oryzae* are emerging [153–156] and we believe that this genomic resource will contribute to the understanding of these processes. Moreover, an emerging trend in the study of insecticide resistance mechanisms lies in detoxification mechanisms modulated by symbiotic associations between bacteria and insects [157].

**Table S7.1. Comparison of the detoxification and insecticide resistance associated genes among coleopteran sequenced genomes.**

| | *Sitophilus oryzae* | *Dendroctonus ponderosae* | *Leptinotarsa decemlineata* | *Diabrotica virgifera* | *Anoplophora glabripennis* | *Aethina tumida* | *Tribolium castaneum* |
|---|---|---|---|---|---|---|---|
| **Order, Family** | Coleoptera, Curculionidae | Coleoptera, Curculionidae | Coleoptera, Chrysomelidae | Coleoptera, Chrysomelidae | Coleoptera, Cerambycidae | Coleoptera, Nitidulidae | Coleoptera, Tenebrionidae |
| **Cytochrome P450 (CYP)** | 148 | 96 | 93 | 185 | 126 | 112 | 126 |
| **Carboxyl/choline esterases (CCE)** | 67 | 57 | 100 | 118 | 109 | 58 | 50 |
| **Glutathione S-transferases (GST)** | 47 | 43 | 39 | 51 | 43 | 49 | 46 |
| **UDP-glucuronosyltransferases (UGT)** | 31 | 24 | 43 | 50 | 54 | 60 | 40 |
| **ABC transporters (ABCs)** | 65 | 85 | 117 | 124 | 70 | 70 | 77 |

**Large supplementary table (Supplemental Table 5):**

**Table S7.2. List of all detoxification and insecticide-related genes annotated in *S. oryzae*.**

### 8. Odorant receptors

Nicolas MONTAGNE, Camille MESLIN, André DA SILVA BARBOSA, Emmanuelle JACQUIN-JOLY

#### 8.1 Introduction

The evolution of the sense of smell is expected to play an important role in the adaptation of insects to diverse ecological niches, as these animals notably use olfactory cues for finding food, mating partners and oviposition sites [158]. *Sitophilus* spp. are known to use kairomones for host detection [159,160] as well as aggregation pheromones [161,162]. The main gene family responsible for the detection of odorants in insect olfactory organs is the odorant receptor (OR) family. Insect ORs are 7-transmembrane domain receptors that form heteromeric complexes with an obligate and universal co-receptor named Orco [163]. They evolved from gustatory receptors, most likely in a common ancestor of winged insects [164]. OR genes have experienced a highly dynamic evolution in insects, and their number greatly varies among insect lineages, ranging from 10 in the human body louse to more than 300 in several ant species [165]. Whereas OR gene repertoires have now been annotated in numerous species, response spectra of insect ORs have been thoroughly investigated only in a handful of species, i.e. the fruit fly *Drosophila melanogaster* [166], the malaria mosquito *Anopheles gambiae* [167,168], the moths *Spodoptera littoralis* [169] and *Helicoverpa armigera* [170] and the ant *Harpegnathos saltator* [171,172]. Whereas Coleoptera represents the largest insect order, only six coleopteran ORs have been functionally characterized to date: three in the cerambycid beetle *Megacyllene caryae* [173], two in the bark beetle *Ips typographus* [174] and one in the red palm weevil *Rhynchophorus ferrugineus* [175], the latter two species belonging to the family Curculionidae. As of today, OR gene repertoires have been manually annotated from ten coleopteran genomes [176], showing that the number of OR genes is highly variable between coleopteran families. Here, we report the annotation of the full repertoire of candidate OR genes in *S. oryzae*.

#### 8.2 Methods

OR amino acid sequences annotated from other coleopteran genomes (1 036 sequences) were used as a query against the *S. oryzae* genome using tBLASTn, with an e-value threshold of 1e-10. Fifty scaffolds presented a hit with at least one of the query sequences. To define precise intron/exon boundaries, the same OR amino acid sequences were then aligned on these scaffolds using Scipio [177], Exonerate [178] and Genewise [179], with default parameters. These alignments were used to generate gene models in WebApollo. Gene models were manually curated based on homology with other coleopteran OR genes. The data set used to build the OR phylogeny contained amino acid sequences of *S. oryzae* candidate ORs plus sequences annotated from the genomes of *Agrilus planipennis* (Buprestidae), *Nicrophorus vespilloides* (Silphidae), *Tribolium castaneum* (Tenebrionidae), *Leptinotarsa decemlineata* (Chrysomelidae) and *Dendroctonus ponderosae* (Curculionidae), removing only those annotated as pseudogenes [176]. Amino acid sequences were aligned using Muscle [2] and the maximum-likelihood phylogeny was built using PhyML 3.0 [180]. The best-fit model of protein evolution was determined using SMS [181]. Node support was assessed by carrying out a hierarchical likelihood-ratio test [182].

#### 8.3 Results and discussion

We annotated 100 candidate OR genes in *S. oryzae* (named SoryORs), including the gene encoding the co-receptor Orco. Of these genes, 46 were predicted to encode a full-length sequence. The global size of the SoryOR gene repertoire is in the range of what has been described in other species of the coleopteran suborder Polyphaga (between 46 in the emerald ash borer *A. planipennis* and >300 in *T. castaneum*) and close to the number of OR genes annotated in the closely related species *D. ponderosae* (85 genes, [176]).

In the phylogeny (Figure S8.1), all SoryORs clustered within the 9 coleopteran OR subfamilies recently described [176]. As already observed for *D. ponderosae*, we found no SoryOR belonging to groups 3 and 6, and only a few SoryORs belonging to groups 4 and 5A. On the other hand, 40 SoryORs clustered within group 7, representing the highest number of OR genes found in this subfamily, for any coleopteran species. Some of these SoryOR genes likely resulted from recent duplications, as they were found in tandem arrays in some scaffolds (e.g. SoryOR32-33-34 and SoryOR52-53-54-55). Interestingly, group 7 contains the pheromone receptors characterized in the Curculionidae *I. typographus* [174] and *R. ferrugineus* [175], whose pheromone compounds are structurally related to the *S. oryzae* pheromone. One can speculate that the pheromone receptors of *S. oryzae* probably belong to this clade. Most of these 40 SoryORs show orthologous relationships only with ORs from *D. ponderosae*, thus suggesting that they result from an expansion specific to the family Curculionidae (Figure S8.1). We also identified a relatively large number of SoryORs belonging to group 1 (15 SoryORs, including 6 SoryOR genes in tandem array on scaffold NW_022147046.1) and group 2B (6 SoryORs). These two groups are the ones containing the three pheromone receptors characterized in the cerambycid beetle *M. caryae* (two in clade 2B and one in clade 1 [173]).

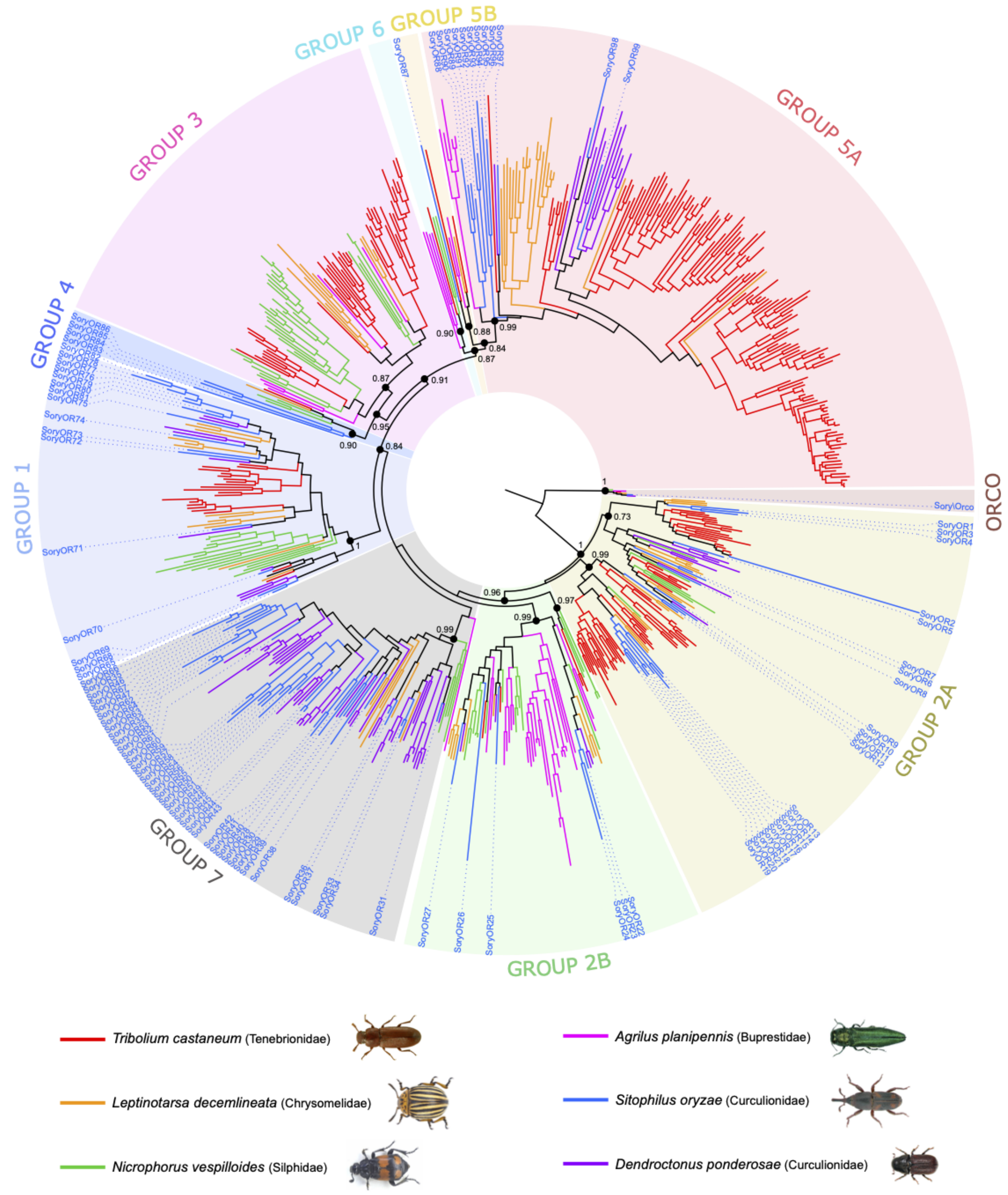


**Figure S8.1. Maximum-likelihood phylogeny of coleopteran ORs.** The amino acid dataset included OR sequences from *S. oryzae* and 5 other coleopteran species [149]. The tree was rooted using the OR co-receptor clade as the out-group. Groups 1 to 7 refer to the coleopteran OR sub-families described previously [149]. Support values indicated correspond to the result of the likelihood ratio-test (aLRT values).

### 9. Epigenetic pathways

Théo CHANCY, Caroline BLANC, Agnès VALLIER, Gautier RICHARD, Carlos VARGAS-CHAVEZ, Cristina VIEIRA and Rita REBOLLO

#### 9.1 Introduction

TEs are usually silenced by host epigenetic mechanisms as chromatin remodeling factors, DNA methylation and small RNA molecules [183]. Given the high propensity of TEs in *S. oryzae*'s genome, we have undertaken to annotate genes belonging to each of these pathways. Very few studies have focused on the epigenetic regulation of TE sequences in Coleoptera, nor on the epigenetic dynamics in such insects [184,185].

Small RNAs can be generally categorized into microRNAs (miRNAs), small interfering RNAs (siRNAs) and piwi interacting RNAs (piRNAs). Each small RNA class possess a different biogenesis pathway, involving specific RNA binding proteins: the endoribonuclease DICER, and the RNA induced silencing complex (RISC) are involved in miRNA (DICER 1) and siRNA (DICER2) silencing in *D. melanogaster,* along with two argonaute proteins, AGO1, and AGO2, respectively [186,187]. Other small RNA synthesis key components (PIWI, AGO3 and AUB) are associated with piRNA biogenesis [187,188] which are able to silence TEs post-transcriptionally, but also to guide repressive histone post-translational modification complexes to TE sequences [189,190]. While in dipterans the piRNA silencing pathway is germline specific, most non-dipteran insects analyzed to date show functional piRNA pathways in both the soma and the germline [191].

While histone post-translational modifications are rather ubiquitous, large groups of insect species are mostly devoid of DNA methylation, including the model species *D. melanogaster* [184,192,193]. Within Coleoptera, *A. glabripennis*, *L. decemlineata, Aleochara curtula, Nicrophorus vespilloides, Onthophagus taurus, A. planipennis* and *Pyractomena borealis* harbour DNA methylation, while *Dendroctonus* *ponderosae, T. castaneum* and *G. marinus* have lost it [184,193,194], suggesting a patchy distribution of DNA methylation in the Coleoptera order. The DNA methylation toolkit comprises two methyltransferases (DNMT1 for DNA methylation maintenance, and DNMT3 for *de novo* DNA methylation) and a variable number of methyl binding domain proteins (MBDs). In insects, DNMT1 is sufficient to maintain a functional DNA methylation pathway, as seen with the Coleopterans *A. glabripennis* and *L. decemlineata* [184]. TET proteins are also part of the DNA methylation toolkit by playing a crucial role in CpG DNA demethylation in animals [195], and in 6mA demethylation in *D. melanogaster* as a DNA N6-methyladenine (6mA) demethylase (DMAD) [196]. While DNA methylation has been associated with TE repression in mammals and plants [184,191,197–199], no direct evidence has been observed in insects, and only the desert locust, *Schistocerca gregaria*, shows DNA methylation present within TE copies [109].

Here we annotated genes involved in chromatin remodelling, DNA methylation, and small RNA pathways. In addition we provide the first *S. oryzae* methylome, and confirm genes central to the piRNA pathway are transcriptionally expressed in germline tissues.

#### 9.2 Methods

**Annotation of epigenetic writers, erasers and readers**

In order to annotate the epigenetic toolkit in *S. oryzae*'s genome, we took advantage of previously annotated insect genomes (*D. melanogaster, A. mellifera* and*, T. castaneum*), and retrieved all genes described to be involved in DNA methylation and small RNAs (Table S9.1). We used blastp and tblastx [45] to search for protein homologs in *S. oryzae.* For chromatin remodelling factors, we have simply searched for *S. oryzae* genes annotated as histone methyltransferases, histone deacetylases, histone acetyltransferases and histone demethylases.

**Bisulfite sequencing (BS-seq) and DNA methylation analysis**

*S. oryzae* laboratory strain (Bouriz) were reared on wheat grains at 27.5 °C and at 70% relative humidity. Ovaries were dissected in diethylpyrocarbonate-treated Buffer A (25 mM KCl, 10 mM MgCl2, 250 mM sucrose, 35 mM Tris/HCl, pH = 7.5). DNA was extracted from *S. oryzae* ovaries using a classical phenol-chloroform extraction. Bisulfite-sequencing and library construction was done in technical duplicates within the framework of an epigenetic workshop (ReaCTION-INRAE, publication in preparation). Briefly, 500 ng of DNA were mixed with 50 ng of unmethylated lambda phage (Sigma) in 40 µL RNAse and DNAse free water. Epitect® Fast Bisulfite Conversion Kit was used on this DNA pool following manufacturer's recommendations to convert unmethylated cytosines to thymine (Low-concentration samples protocol). Subsequently to unmethylated cytosine conversion, DNA fragments have been double size selected using SPRIselect magnetic beads (Beckman Coulter, double size selection protocol): in a 50 µL volume of cytosine converted DNA sample (completed using low-EDTA TE), size selection has been performed following manufacturer's recommendation with 0.7x sample volume SPRI beads (35 µL) and subsequently with 1.1x sample volume SPRI beads (55 µL) to remove small and large DNA fragments. Sequencing library construction of the size selected samples was then performed using Accel-NGS® Methyl-seqDNA library kit and Accel-NGS® Methyl-Seq Set A Indexing Kit (Swift Biosciences). Libraries were sequenced in an Illumina Hiseq3000 sequencer in paired-end mode at 100 bp size (PRJNA681724). We obtained ≈63M reads per replicate, representing around 30X of *S. oryzae*'s genome. Reads were cleaned with TrimGalore ( --paired -q 30 --clip_R2 18 --clip_R1 9), and mapping, deduplication and methylation extraction was performed using Bismark 0.22.1 (-N 1 -X 900 [200]) against *S. oryzae*'s genome. Around 57% of reads were uniquely mapped against *S. oryzae*'s genome according to bowtie2 [201]. We took advantage of *S. pierantonius* naturally unmethylated genome to calculate cytosine conversion rate (99.7%). Thanks to CpG coverage profiles obtained from Bismark 0.22.1 [200], we pooled both replicates CpG coverage information and filtered for cytosines with coverage higher than five reads. Meta-graphs were obtained using deepTools 3.3.0 [202] while intersection between genomic regions and methylated cytosines were performed with Bedtools v2.27.1 [203]. TE CpG methylation was calculated by mapping bisulfite reads against TE consensus sequences and averaging CpG methylation per family.

**Somatic and germline expression of piRNA biogenesis genes**

*S. oryzae* laboratory strain (Bouriz), were reared on wheat grains at 27.5 °C and at 70% relative humidity. Seven day old adult midgut (anterior and posterior), ovaries and testis were dissected in diethylpyrocarbonate-treated Buffer A (25 mM KCl, 10 mM MgCl2, 250 mM sucrose, 35 mM Tris/HCl, pH = 7.5). Total RNA was extracted using the RNAqueous - Micro kit (Ambion). DNA was removed with DNAse treatment and RNA quality was checked with Nanodrop (Thermo Scientific). Complementary DNA (cDNA) was produced with the iScript™ cDNA Synthesis Kit (BioRad) following the manufacturer's instructions and starting with 500 ng total RNA. Differential gene expression was assessed by quantitative real-time PCR with a CFX Connect Real-Time PCR Detection System (Bio-Rad) using the LightCycler Fast Start DNA Master SYBR Green I kit (Roche Diagnostics), as previously described [105]. Data were normalized using the ratio of the target cDNA concentration to the geometric average of two housekeeping genes: glyceraldehyde 3-phosphate dehydrogenase (LOC115881082) and malate oxidase (LOC115886866). Primers were designed to amplify fragments of approximately 90 bp. Primers are available upon request. Graphs and statistical tests were performed on GraphPad Prism 9.

#### 9.3 Results and Discussion

***S. oryzae* has a functional DNA methylation toolkit**

*S. oryzae* harbours one full length DNMT1.1 and a second DNMT1.2 gene that seems to be missing a DNMT1 replication foci domain and a zinc finger domain. We verified the presence of both genes in the genome, and DNMT1.2 might stem from an assembly error as we could not amplify this loci by PCR. We have also detected one copy of DNMT2, also known as TRDMT1 (an RNA methyltransferase, Table S9.1A), but no DNMT3 was detected. Furthermore *S. oryzae* has two full length MBD2/3, and two genes that harbor similarities to MBD4 and MBD5 (Table S9.1). Hence, *S. oryzae* harbours a potentially functional DNA methylation toolkit, despite the lack of DNMT3 [184]. TET genes are involved in active DNA demethylation in mammals and honey bees [204,205], but also as 6mA demethylase in dipterans that lack DNA methylation [196,206]. We were able to detect two genes encoding TET proteins, as opposite to the single TET protein often described in other insects [207].

In order to verify if such DNA methylation machinery is indeed active, we have performed BS-seq on *S. oryzae's* ovaries. We were able to uncover ≈1% of CpG methylation according to Bismark reports [200], while no DNA methylation is observed in CHH and CHG contexts (Figure S9.1A). Distribution of CpG methylation shows enrichment around gene transcriptional start sites (TSSs) and exons (Figure S9.1B and C). We have also searched for DNA methylation within TE copies. Unfortunately BS-seq is not appropriate to study DNA methylation within repeated sequences as it can only take into account uniquely mapping reads. Our preliminary analysis shows overall low DNA methylation within TE consensus sequences (S9.1D). Nevertheless, some DNA, LTR and Unknown families harbor higher CpG methylation (> 20%) suggesting TE sequences might be the target of CpG methylation in *S. oryzae*. Such analysis should be confirmed by new methods using long reads and allowing for TE copy methylation analysis [183]. In conclusion, *S. oryzae* harbors a functional DNA methylation machinery resulting in ≈1% of CpG methylation in ovaries, mainly observed within exons and close to gene TSSs.

***S. oryzae*'s piRNA pathway is potentially active in germ and somatic cells**

The complete miRNA and siRNA biogenesis pathways seem complete in *S. oryzae* as we detected AGO1, AGO2, DICER1, DICER2 and DROSHA. Regarding the piRNA pathway, we were able to retrieve AGO3 and SIWI, as observed in non-dipteran insects [191], but also Armi, Hen1, Spindle-E, Zucchini and Vasa. We wondered if the piRNA pathway could be active in both soma and germline tissues, and henceforth performed a quantitative RT-PCR targeting genes involved in piRNA biogenesis in somatic and germline tissues of *S. oryzae* (Figure S9.2). *S. oryzae* carries intracellular symbiotic bacteria in ovarian apexes and midgut caeca which could impact the activity of small RNA pathways. Henceforth, we have dissected germline and somatic tissues devoid of bacteria (testis, ovaries without apexes and posterior midguts), and containing bacteria (ovarian apexes and anterior midguts). All genes assayed, with the exception of Hen1, show the same expression pattern, with upregulation in ovaries, especially ovarian apexes where oocytes and the endosymbiont *S. pierantonius* are present, compared to testis and somatic tissues (midgut harboring intracellular bacteria, and posterior midgut devoid of any bacteria). Henceforth, despite the presence of SIWI, known to participate in piRNA biogenesis in somatic and germ tissues [191], *S. oryzae* presents higher expression of piRNA related genes in ovaries than in somatic tissues or the male germline. Further analyses are necessary in order to verify if piRNAs are able to control TE expression in the germline or soma.

**Survey of chromatin remodeling factors**

Contrary to DNA methylation, there are several writers and readers of histone modifications. By mining NCBI annotations we were able to find 28 histone acetyltransferases, 7 histone deacetylases, 26 histone methyltransferases, and two histone demethylases.

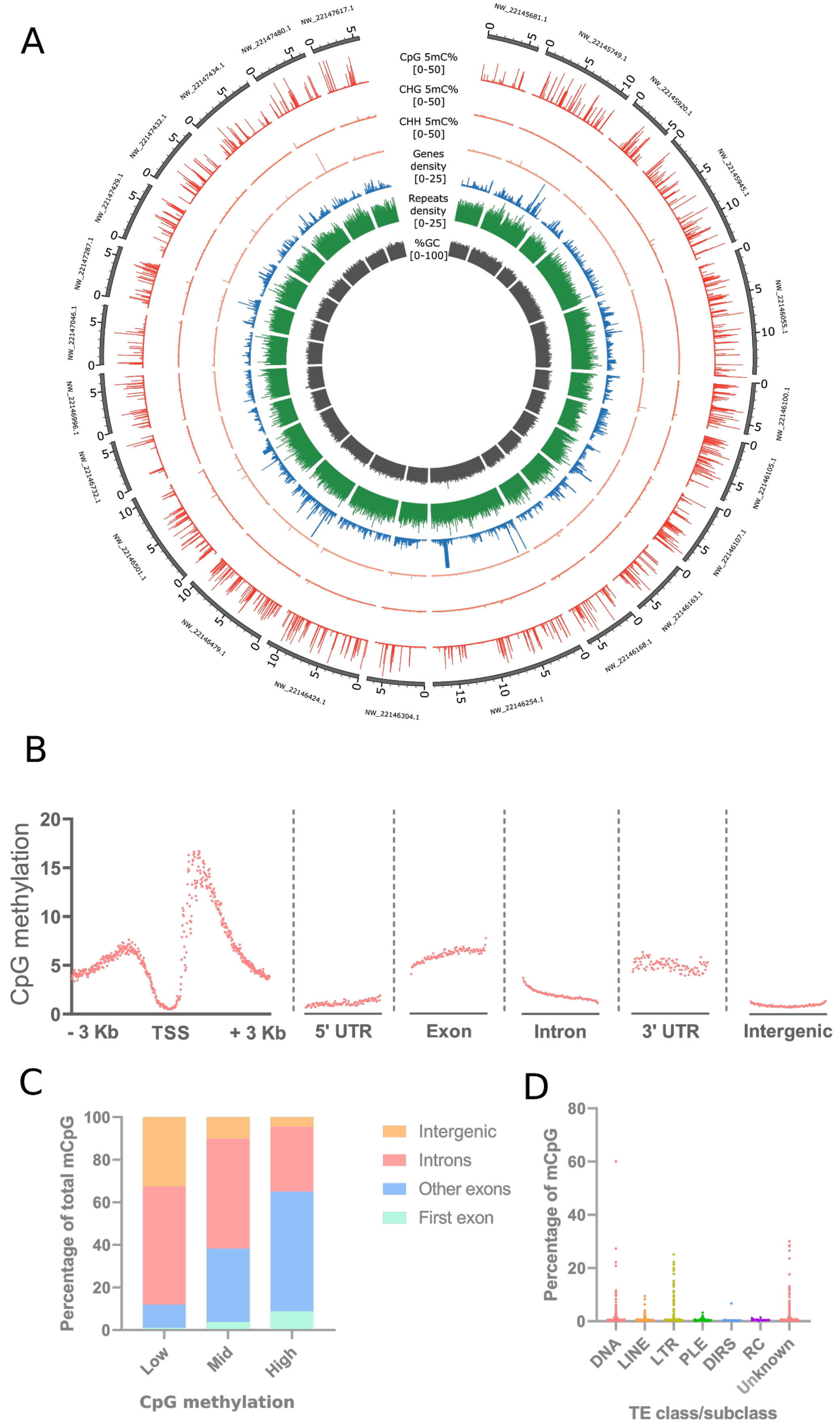
A
CpG 5mC%
[0-50]
CHG 5mC%
[0-50]
CHH 5mC%
[0-50]
Genes
density
[0-25]
Repeats
density
[0-25]
%GC
[0-100]
B
CpG methylation
- 3 Kb
TSS
+ 3 Kb
5' UTR
Exon
Intron
3' UTR
Intergenic
C
Percentage of total mCpG
Low
Mid
High
CpG methylation
Intergenic
Introns
Other exons
First exon
D
Percentage of mCpG
DNA
LINE
LTR
PLE
DIRS
RC
Unknown
TE class/subclass

**Figure S9.1. DNA methylation in *S. oryzae.*** A. Circos plot representing, for the 24 largest scaffolds, from the external to internal parts of the plot, using genomic signals binned in 500bp windows: name of the scaffold and genomic location (dark grey lines), methylation levels in the CpG (red), CHG (orange), CHH (light orange) context averaged between the two replicates and averaged across cytosines in each bin ; density of genes (blue) and repeated elements (green) (sum of bed features per bin) ; average GC content per bin (grey). 5mC are detected only in the CpG context. B. Distribution of CpG methylation across genomic features. CpG methylation is depleted around TSSs and 5' untranslated regions (UTRs), but high in exons and 3' UTRs. C. Proportion of low (1-30%), mid (30-70%) and high (>70%) CpG methylation across genomic features. High CpG methylation is observed at exons. D. Average CpG methylation within TE consensus sequences. Most TE class/subclasses show low CpG methylation. Within DNA, LTR and unknown subclasses, a few TE families harbord average mCpG higher than ≈20%, suggesting TEs could be the target of CpG methylation in *S. oryzae.*

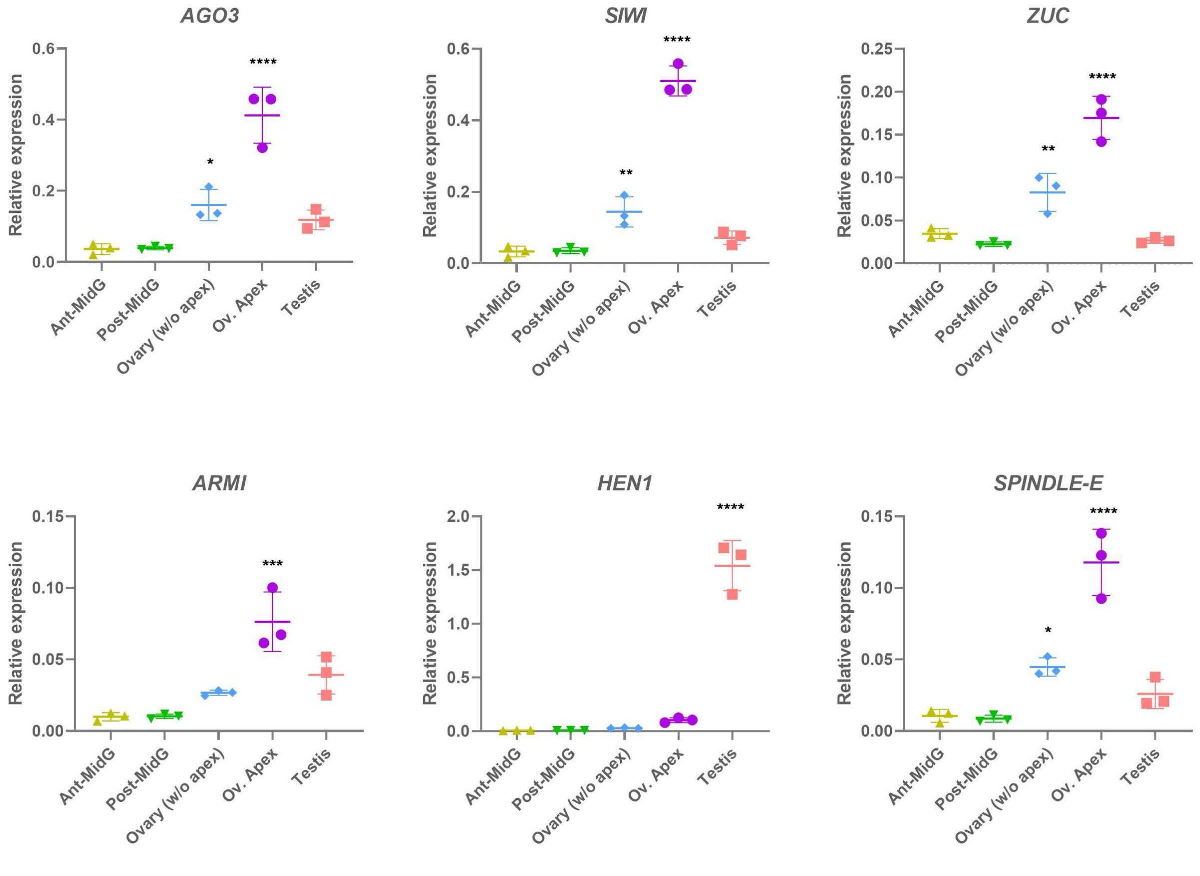


**Figure S9.2. piRNA associated genes transcription survey in somatic and germ tissues.** Ant-MidG: anterior midgut, contains symbiotic bacteria. Post-MidG: posterior midgut, no symbiotic bacteria. Ovary (w/o apex): entire ovarian chambers without the apexes. Ov. Apex: ovarian apexes, where symbiotic bacteria are present. AGO3: argonaute 3, ZUC: zucchini, ARMI: armitage. Most piRNA associated genes show the same transcriptional profile: higher steady-state expression level in germ tissues, especially ovarian apexes. Anova with tukey. Asterisks denote p-value significance (**** p-value <0.001, *** p-value between 0.0001 to 0.001, ** p-value 0.001 to 0.01 and lastly * p-value between 0.01 and 0.05.

**Table S9.1. List of genes related to epigenetic mechanisms in *S. oryzae*.**

| Protein | Gene ID | Coordinates |
|---|---|---|
| AGO1 | LOC115881833 | NW_022146439.1 (685290..770283, complement) |
| AGO2 | LOC115885402 | NW_022146953.1 (1989339..2007681) |
| AGO3 | LOC115879776 | NW_022146265.1 (2451081..2472039, complement) |
| SIWI | LOC115878363 | NW_022146179.1 (936219..1005229, complement) |
| Dicer-1 | LOC115887113 | NW_022147143.1 (323842..341798, complement) |
| Dicer-2 | LOC115880804 | NW_022146347.1 (1863075..1998078, complement) |
| Drosha | LOC115876974 | NW_022146106.1 (1327841..1354553, complement) |
| DNMT1.1 | LOC115882389 | NW_022146479.1 (4131629..413967) |
| DNMT1.2 (tested by PCR - assembly error) | LOC115883131 | NW_022146520.1 (1329669..1379208) |
| DNMT2 | LOC115883430 | NW_022146616.1 (392145..393538, complement) |
| MBD2/3 | LOC115876162 | NW_022146076.1 (33426..38183, complement) |
| MBD2/3 | LOC115883257 | NW_022146561.1 (9412..14911) |
| MBD | LOC115879534 | NW_022146261.1 (2314950..2368917, complement) |
| MBD | LOC115874592 | NW_022145945.1 (3890683..3923221, complement) |
| TET | LOC115876270 | NW_022146094.1 (1249792..1329131, complement) |
| TET | LOC115883839 | NW_022146731.1 (7..3200, complement) |
| Armitage | LOC115890098 | NW_022147483.1 (1926014..1943769, complement) |
| Hen1 | LOC115884554 | NW_022146866.1 (1641791..1662885, complement) |
| Spindle-E | LOC115886157 | NW_022147050.1 (469728..497481, complement) |
| Vasa | LOC115889396 | NW_022147432.1 (3732419..3755509) |
| Zucchini | LOC115891431 | NW_022145817.1 (225625..226543) |

### 10. Supplemental figures

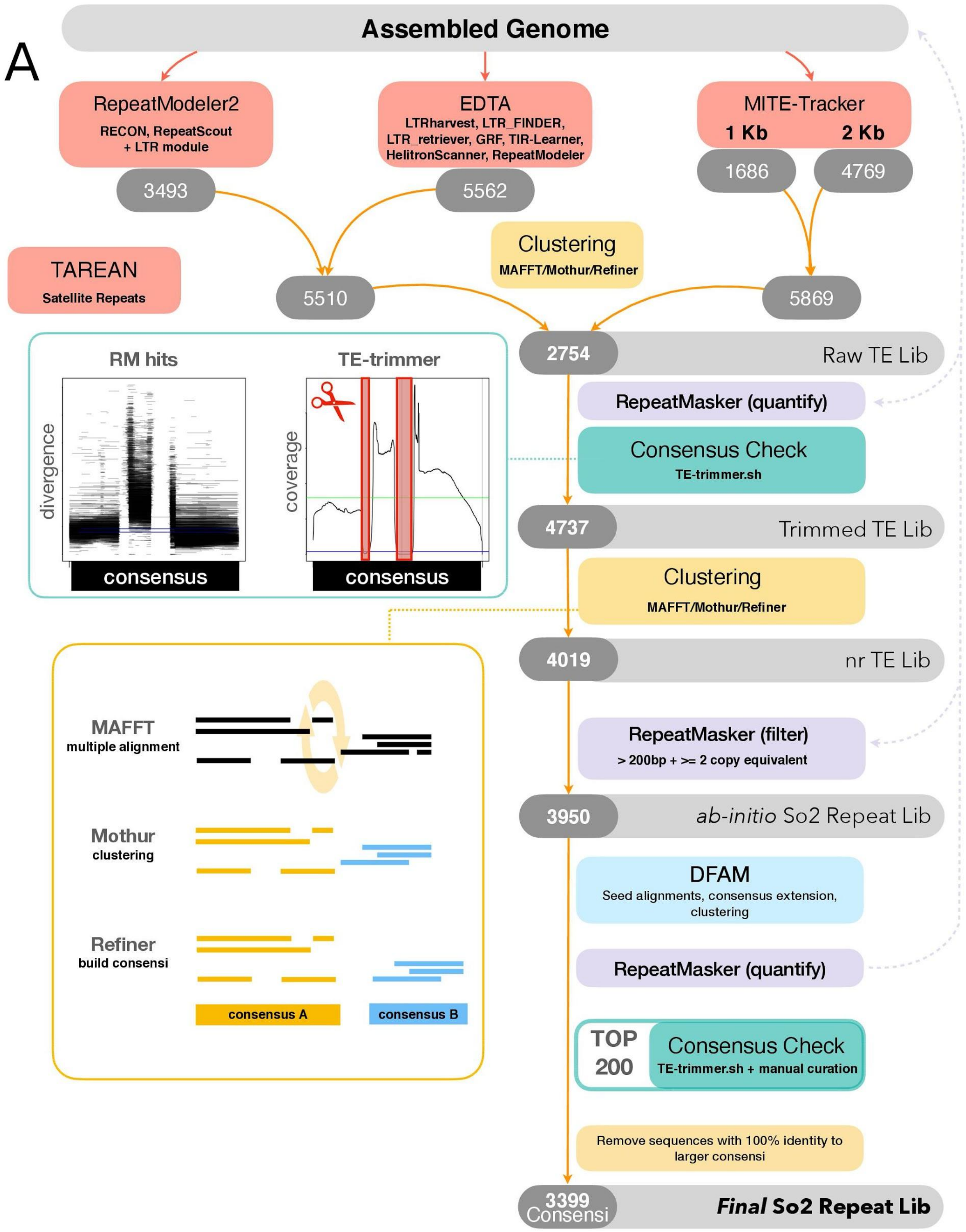

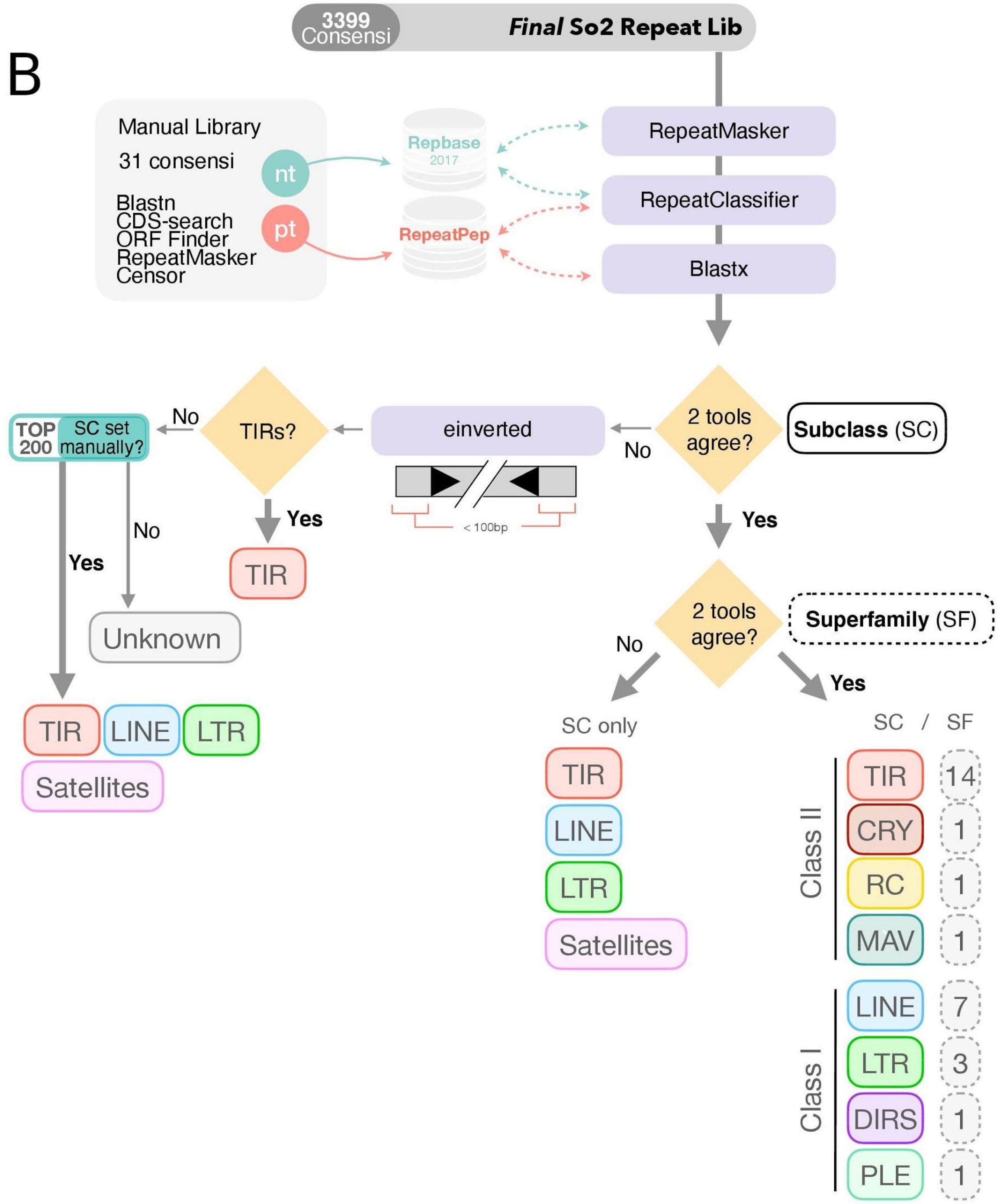


**Supplemental Figure S1. Scheme of TE annotation.** A. Detection and assembly of repeated elements. RepeatModeler 2, EDTA and MITE-tracker were run on the genome assembly sequence. RepeatModeler

2 and EDTA include a variety of modules (indicated in the red boxes) dedicated to assemble repeats and TE families. MITE-tracker was also used to specifically detect repeats with terminal inverted motifs such as DNA/TIR and MITE elements, using two window sizes of 1 and 2 kb. Because the consensus sequences made by these programs likely overlap to some degree, we performed successive clustering rounds using MAFFT/Mothur/Refiner (see Methods). Upon clustering of the initial consensi pool, 2 754 sequences were screened for support using RepeatMasker against the reference genome. The results were analyzed with the script TE-trimmer.sh, which scan consensi for support among the repeat masker hits. Drop in coverage below 5% of the average consensus coverage leads to a split of the original sequence. Over-splitted consensi were then recovered by performing a clustering round. After another instance of RepeatMasker, consensi with no hits or less than the equivalent of two full consensi in length in the genome, were removed from the library. At this point, two Satellites consensi identified by Tarean and manually curated were incorporated to the library. B. Classification of the repeat library. We classified as much consensi as possible up to the superfamily level. First, 31 high quality consensi were manually curated and classified. Then the *ab-initio* automatic library (A) was analyzed with a combination of structural, homology or machine learning tools. Upon completion, TEs are classified if two or more programs agree on the subclass (TIR, MITE, CRYPTON, LTR, SINE, LINE, RC, Maverick). Further, subfamilies can be given following the same rule.

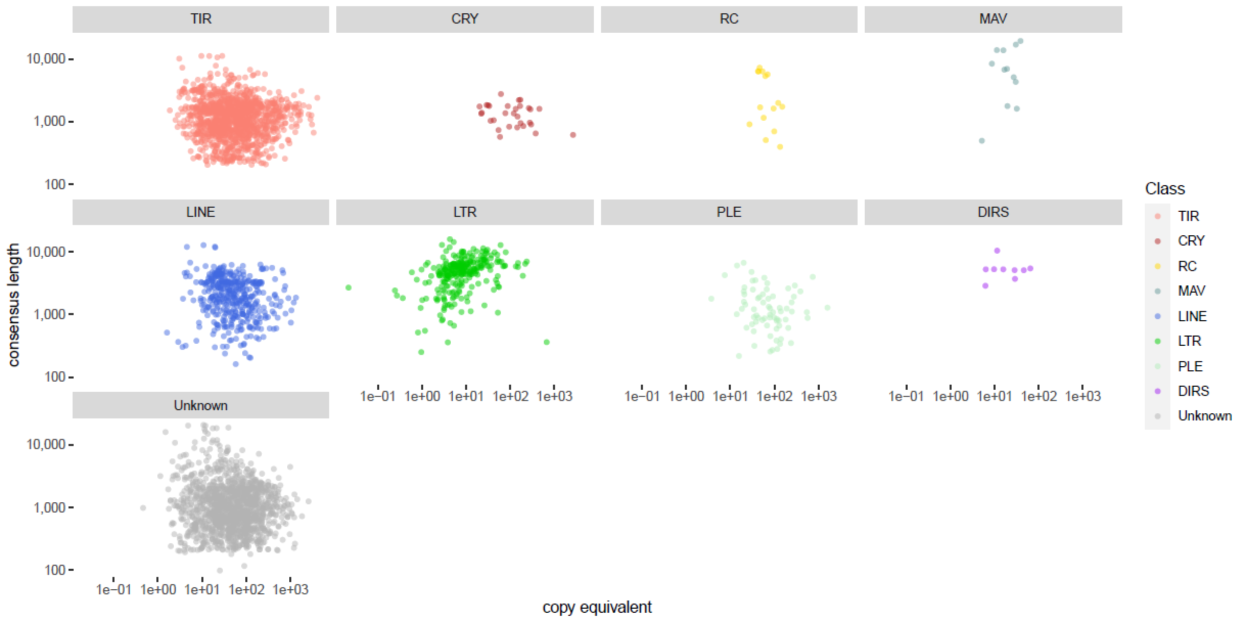


**Supplemental Figure S2. Consensus length as function of the copy equivalent distribution.** While DNA, LINE and Unknown elements show a gradual distribution of TE consensus size, LTR families are contained at a higher molecular size.

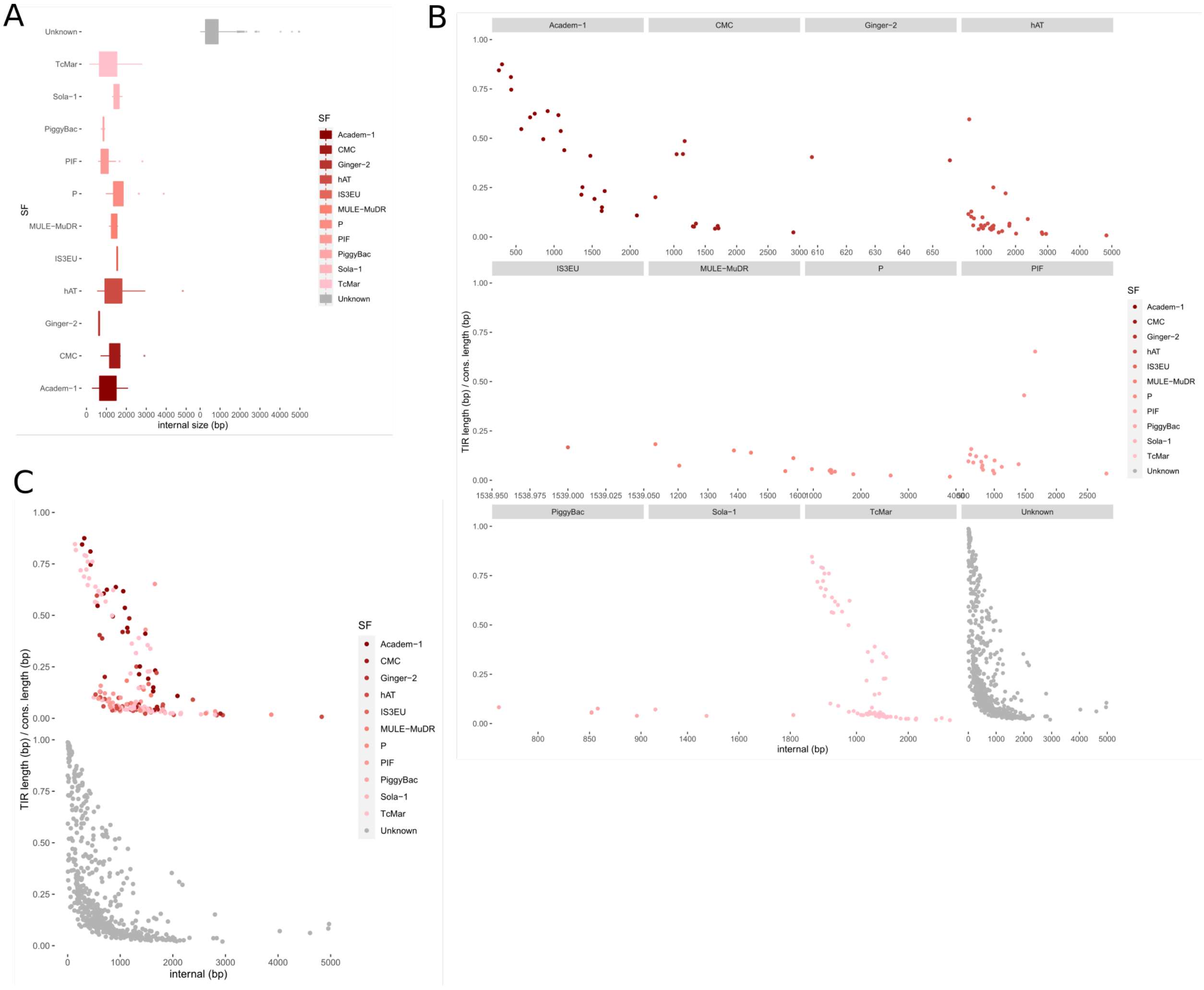


**Supplemental Figure S3. A. TIR element's internal size per superfamily.** Most internal sequences (without TIR sequences) of Unknown DNA elements are short. B. Comparison of TIR length and internal size for all DNA superfamilies. C. Proportion of TIRs (%) vs internal sequence length (bp).

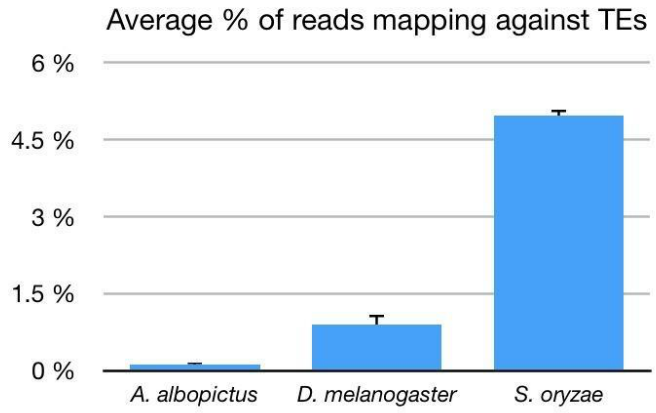


**Supplemental Figure S4. Expression of TE families in *S. oryzae*, *D. melanogaster* and *A. albopictus* based on RNAseq poly-A enriched reads.**
